# The temporal transcriptomic signature of cartilage formation

**DOI:** 10.1101/2022.04.01.486765

**Authors:** Roland Á. Takács, Judit Vágó, Szilárd Póliska, Peter N. Pushparaj, László Ducza, Patrik Kovács, Eun-Jung Jin, Richard Barrett-Jolley, Róza Zákány, Csaba Matta

## Abstract

Chondrogenesis is a multistep process, in which cartilage progenitor cells generate a tissue with distinct structural and functional properties. Although several approaches to cartilage regeneration rely on the differentiation of implanted progenitor cells, the temporal transcriptomic landscape of *in vitro* chondrogenesis in different models has not been reported. Using RNA sequencing, we examined differences in gene expression patterns during cartilage formation in micromass cultures of embryonic limb bud-derived progenitors. Principal component and trajectory analyses revealed a progressively different and distinct transcriptome during chondrogenesis. Differentially expressed genes (DEGs), based on pairwise comparisons of samples from consecutive days were classified into clusters and analysed. We confirmed the involvement of the top DEGs in chondrogenic differentiation using pathway analysis and identified several chondrogenesis-associated transcription factors and collagen subtypes that were not previously linked to cartilage formation. Transient gene silencing of *ATOH8* or *EBF1* on day 0 attenuated chondrogenesis by deregulating the expression of key osteochondrogenic marker genes in micromass cultures. These results provide detailed insight into the molecular mechanism of chondrogenesis in primary micromass cultures and present a comprehensive dataset of the temporal transcriptomic landscape of chondrogenesis, which may serve as a platform for new molecular approaches in cartilage tissue engineering.

**GRAPHICAL ABSTRACT:** 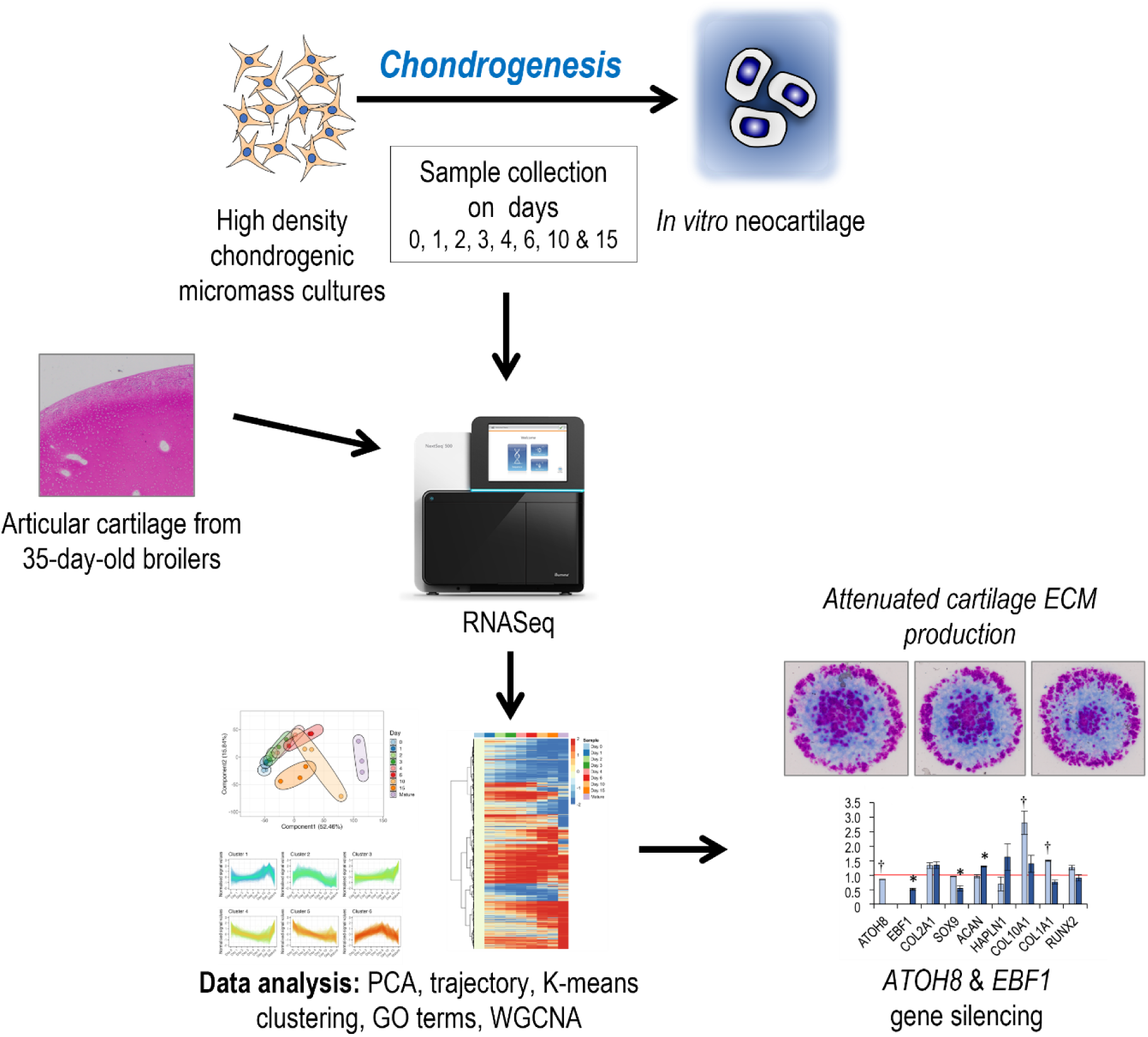
This study provides insights into the mechanisms of cartilage formation. Gene expression patterns during cartilage formation in micromass cultures were analysed using RNA sequencing. Silencing *ATOH8* or *EBF1* attenuates chondrogenesis.

**Key points:**

• We examined the global gene expression patterns during in vitro chondrogenesis.
• Using WGCNA, we created a module of genes with patterns similar to those of SOX9, ACAN, and COL2A1.
• We identified ATOH8 and EBF1 transcription factors with a yet unexplored role in chondrogenesis.

## INTRODUCTION

During morphogenesis of the vertebrate appendicular skeleton, progenitor cells derived from the embryonic mesoderm undergo specification, proliferation, condensation, and nodule formation (1). The cells in these nodules, termed limb bud mesenchymal progenitors (LMPs), specify into the osteochondrogenic lineage and form the supporting and connective tissues of the developing limb: bone, tendon, and cartilage. Young chondroblasts continue to proliferate and form the structures of the future skeleton. During this process, chondroblasts begin to produce a cartilage-specific extracellular matrix (ECM) that gradually changes as the tissue matures. Collagen is the most abundant macromolecule in the ECM and accounts for approximately two-thirds of the dry weight of cartilage (2). Collagens stabilise the cartilage matrix and provide tensile and shear strength. Many different types of collagen molecules are expressed in articular cartilage; however, the backbone polymeric framework during development is a copolymer of collagen types II, IX, and XI. Over 90% of the collagens in hyaline cartilage ECM is type II collagen (3). However, very little is known about the other types of collagens synthesised by differentiating chondrocytes during chondrogenesis. Knowledge of the molecular structure of the collagen framework of articular cartilage and its development, remodelling, and maturation through various pericellular, territorial, and interterritorial domains is critical for understanding the mechanisms of its degradation in disease and possible regeneration.

The canonical pathway for chondrogenesis in bone primordia involves terminal chondrocyte differentiation, followed by hypertrophy, apoptosis, and the induction of bone formation. This default pathway is blocked in articular cartilage, resulting in permanent cartilage (4). Although its unique composition allows joints to move with little friction and acts as a shock absorber, articular cartilage has little or no ability to regenerate following an injury or disease. Therefore, diseases or disorders affecting articular cartilage often result in progressive long-term pain and disability (5). Therefore, significant efforts have been made to develop novel approaches, often based on a combination of biomaterials and stem cells, to enhance intrinsic cartilage repair, regenerate new cartilage to repair focal defects, or restore the joint surface (6).

Mesenchymal stem cells (MSCs) are promising candidates for cartilage tissue engineering owing to their potential to differentiate into chondrocytes (7, 8). However, increasing evidence suggests that the phenotype of chondrocytes differentiated from MSCs for cartilage repair is unstable (9). It is plausible that the canonical differentiation pathway of bone marrow-derived MSCs is toward a hypertrophic phenotype, raising concerns about their applicability in tissue engineering of cartilage, as hypertrophy of chondrocytes in neocartilage may ultimately lead to apoptosis, cartilage mineralisation, and/or ossification (10). However, lineage-tracing data have recently provided evidence that hypertrophic chondrocytes can transdifferentiate into osteoblasts, osteocytes, and other lineages, such as adipocytes and pericytes (11). Nevertheless, adult MSCs are probably primed for endochondral ossification and the chondrogenic state is transient. Therefore, to improve cartilage tissue engineering strategies, it may be logical to use embryonic progenitor cells to improve cartilage tissue engineering strategies (12).

The chicken limb bud-derived 3-dimensional micromass assay uses embryonic LMPs and represents a widely used, adaptable, and relatively simple *in vitro* model of the early stages of skeletal development (12). LMPs in high density micromass cultures self-organise to produce a nodular pattern, which is a prerequisite for chondrogenic differentiation (13). This model is particularly useful for studying early events of chondrogenesis. Several molecular pathways regulate in this complex process (14). However, the time course of transcriptomic events involved in the regulation of these pathways during *in vitro* chondrogenesis has not been fully explored. Recent studies have investigated this process by differential gene expression analysis on microarray platforms (15) and next-generation sequencing (NGS) in adult MSCs differentiated into chondrocytes in pellet cultures (6). However, as described above, the primary micromass model established from embryonic LMPs more consistently recapitulates chondrogenesis of hyaline cartilage *in vivo*. Therefore, we performed a quantitative study on the transcriptomic landscape of chondrogenesis using this model. RNA sequencing (RNA-seq) was performed at eight key time points (between culture days 0 and 15, the latter representing mature micromasses with differentiated chondrocytes) during *in vitro* chondrogenesis in micromass cultures established from freshly isolated LMPs. In this study, we also included chondrocytes obtained from mature articular cartilage from the knee joint. Bioinformatics analysis of the transcriptome revealed several dominant gene signatures associated with chondrogenesis. We identified the transcripts of several “minor” collagen types in differentiating micromass cultures not previously reported in cartilage, and found that the LMP-based micromass model was superior to chondrogenic MSC cultures in terms of cartilage ECM gene expression profiles. We also detected the expression of potentially novel chondrogenic transcription factors (TFs) and confirmed that atonal homolog 8 (ATOH8) and early B-cell factor 1 (EBF1) are important for chondrogenic differentiation because their transient silencing impaires chondrogenesis. The results of this study provide a deeper understanding of the biology of the chondrogenic lineage during key stages of cartilage formation, which is essential for future therapeutic interventions.

## MATERIAL AND METHODS

### Experimental Design

We used transcriptome profiling by RNA-seq to identify changes in global gene expression as chicken primary limb bud mesenchymal progenitors (LMPs) differentiate into mature chondrocytes (Fig. 1). The limb bud-derived chondrifying micromass model is a well-established method used in our laboratory (16, 17). Samples for RNA-seq were collected on key days of chondrogenic differentiation from micromass cultures (days 0, 1, 2, 3, 4, 6, 10, and 15) and subjected to high-throughput mRNA sequencing on an Illumina platform. Total RNA was extracted from the articular cartilage of knee joints of 35-day-old broilers and subjected to RNA-seq analysis. Raw sequencing data were aligned to the chicken reference genome version GRCg6a (Ensembl release 106^1^). Genes with a Benjamini-Hochberg adjusted *p*-value of less than 0.05 and a log2 fold change (LFC) cut-off of ±2.0 were considered differentially expressed. Network analysis and enrichment of Gene Ontology (GO) terms were performed using R, Cytoscape, and Ingenuity Pathway Analysis (IPA) software (Qiagen, USA).

**Figure 1.**
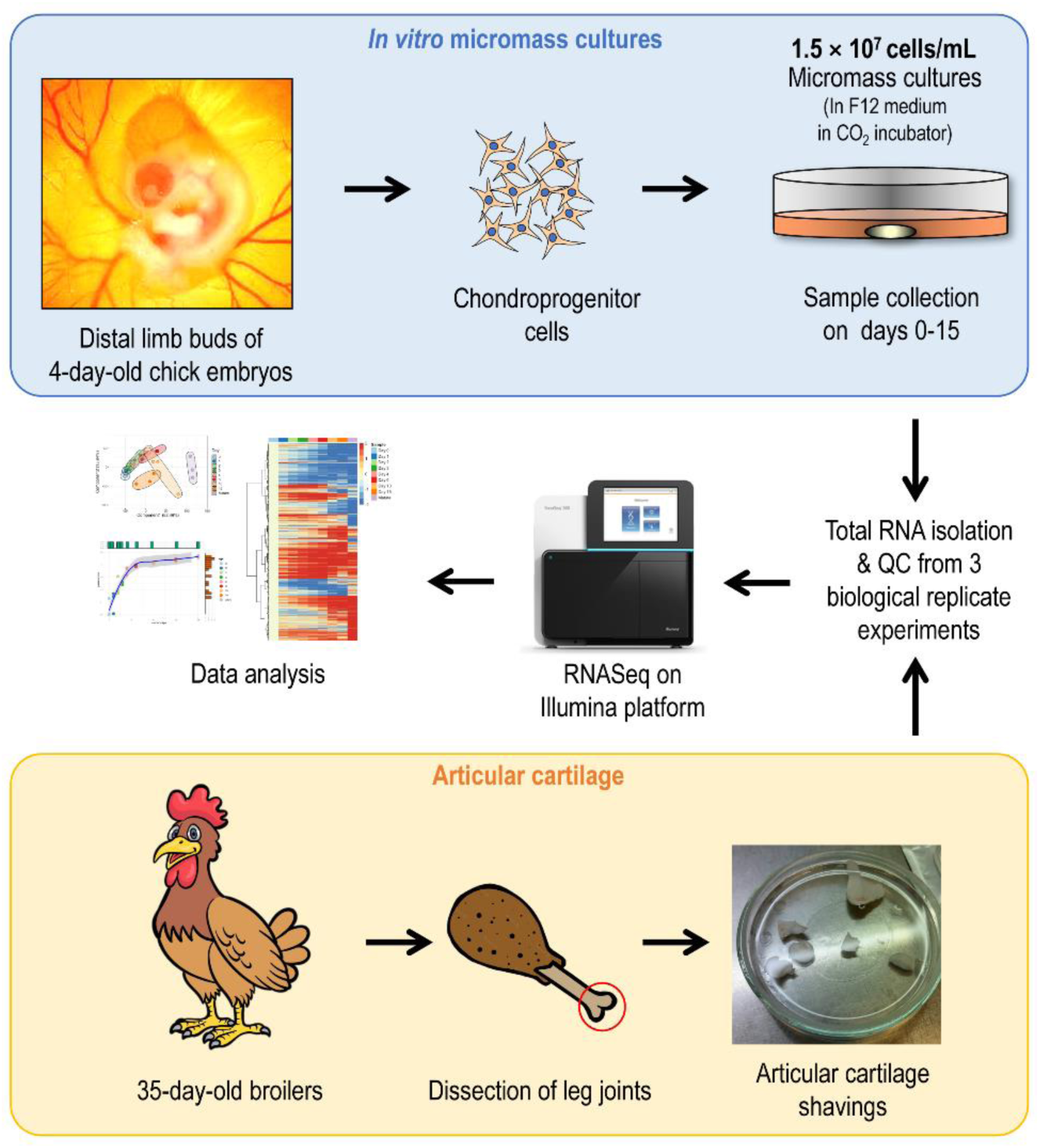
Experimental workflow. Limb bud-derived 3D micromass cultures grown in plasticware (grey cylinder) in F12 medium (light orange) are represented by the yellow oval in the upper right corner. QC, quality check.

### Cell Cultures

Primary chondrifying micromass cell cultures (high density cultures, HDCs) were used as an *in vitro* experimental model for hyaline cartilage formation (18). LMP cells in HDCs first proliferated and then differentiated into matrix-producing chondroblasts during the first three days of culture, and a well-detectable hyaline cartilage-specific metachromatic ECM was formed on the 6^th^ day of culture, which became even more abundant by day 15. HDCs were established as previously described (16,19,20). Briefly, chondroprogenitor cells were isolated from the developing limb buds of chicken embryos at Hamburger–Hamilton stages 22–24 (21). The *in vitro* work on early stage (4.5-day-old) chicken embryos did not require approval from the Ethics Committee of the University of Debrecen. To obtain a sufficiently high yield of primary chondrogenic cells, the limb buds of approximately 100 embryos were collected for each experiment. Distal parts of the fore and hind limb buds of embryos were pooled and dissociated in 0.25% trypsin-EDTA (Sigma-Aldrich, Merck, St. Louis, MO, USA; pH 7.4) at 37 °C for 1 h. The digestion of limb buds was terminated by adding an equal volume of foetal bovine serum (FBS; Gibco, Thermo Fisher Scientific, Gaithersburg, MD, USA). Single-cell suspensions of chondrogenic LMP cells were obtained by filtering through a 20-μm pore-size plastic filter (Millipore, Merck, Billerica, MA, USA). After centrifugation at 800 × *g* for 10 min at room temperature, the cells were resuspended in Ham’s F12 culture medium (Euroclone, Pero, Italy) supplemented with 10% FBS at a concentration of 1.5 × 10^7^ cells/mL. Droplets of various numbers (100 µL; 12–12 droplets for days 0 and 1, 10 droplets for day 2, 8–8 droplets for days 3 and 4, and 6–6 droplets for the remaining days) were inoculated into 150-mm Petri dishes (Eppendorf, Hamburg, Germany), and the cells were allowed to attach to the surface for 2 h in a CO2 incubator (5% CO2 and 90% humidity). Finally, 1.5 mL of Ham’s F12 supplemented with 10% FBS, 0.5 mM L-glutamine, and 1% penicillin/streptomycin (TEVA, Debrecen, Hungary) was added to the colonies. The day of inoculation was considered day 0 of the culture. Cultures were maintained at 37 °C in a CO2 incubator for 1, 2, 3, 4, 6, 10, or 15 days. Cultures designated “day 0” were kept in the incubator for 2 h after being flooded with F12 medium and then harvested. The culture medium was changed every two days.

### Monitoring Cell Culture Morphology, ECM Production, and Differentiation

Additional primary chondrifying cell cultures were inoculated onto the surface of 30-mm round cover slips (3×30-μL droplets) (Menzel-Gläser, Menzel GmbH, Braunschweig, Germany) placed in 35-mm Petri dishes (Eppendorf). Hyaline cartilage-specific ECM production in cultures from days 0 to 15 was qualitatively analysed using metachromatic staining. For Alcian blue (AB) staining, cultures were washed with phosphate-buffered saline (PBS), fixed with a 4:1 mixture of absolute ethanol and formalin for 30 min, and stained with 1% AB (Merck) for 15 min in 3% acetic acid and 1% HCl (pH 1). Excess stain was removed by washing with PBS (modified based on (22)). For Safranin-O staining, formalin-fixed micromasses were stained with 0.1% Safranin-O (Merck) for 15 min, and excess stain was removed by washing with water (modified based on (23)). Dimethyl methylene blue (DMMB; pH 1.8; Sigma-Aldrich, Merck) was applied to the cultures, as previously described (20, 24). Photomicrographs of the stained specimens were obtained using an Olympus BX53 camera on a Nikon Eclipse E800 microscope (Nikon Corporation, Tokyo, Japan). The optical density values of DMMB-stained specimens were determined from HDCs in three independent, biological replicate experiments using a MATLAB image analysis application, where cartilage nodules rich in metachromatic cartilage ECM were defined by an approximate range of values in the RGB colour space, and the pixels were counted. Values were normalised to day 6 values. Photomicrographs show visual representations of three cultures on each culture day in three independent experiments.

### Collagen Type I and II Immunohistochemistry

Immunoperoxidase reactions were carried out on micromass cell cultures seeded onto glass coverslips to demonstrate collagen type I and II expression on designated days of culturing (*i.e*., day 0 (2 h of incubation after being flooded with F12 medium), and then on days 1, 2, 3, 4, 6, 10, and 15). The cultures were fixed with 4% paraformaldehyde (Sigma-Aldrich, Merck) for 30 min and washed with distilled water. Endogenous peroxidase activity was blocked with 1% H2O2 in phosphate buffered saline (PBS) for 10 min. After washing with PBS, non-specific binding sites were blocked with Tris-phosphate buffered saline (TPBS) supplemented with 10% normal goat serum (Vector Labs, Newark, CA, USA) and 3% bovine serum albumin (BSA) (Amresco, VWR, Avantor Sciences, Radnor, PA, USA) for 50 min at room temperature. Thereafter, the samples were incubated with rabbit anti-collagen type I or type II polyclonal antibodies (Novus Biologicals; Bio-Techne; Cat. no. NB600-408 and NB600-844, respectively) at dilutions of 1:500 and 1:250 (respectively) in TPBS supplemented with 1% BSA and 3% goat serum at 4°C overnight. The following day, after extensive washing in PBS, the samples were transferred to biotinylated anti-rabbit IgG (Vector Laboratories; Cat. no. BA-1000-1.5) for 1 h at a dilution of 1:200 in TPBS, followed by additional washing steps. Thereafter, the cultures were treated with extravidin (Sigma-Aldrich, Merck) for 1 h at a dilution of 1:500 in TPBS. After washing in TRIS buffer, immunoreaction was visualised with 3,3’-diaminobenzidine (DAB) chromogen reaction (Sigma-Aldrich, Merck) and background-stained with haematoxylin (Amresco). Finally, the slides were mounted using Pertex medium (VWR, Avantor Sciences) and coverslipped. Photomicrographs were captured using an Olympus BX53 camera and a Nikon Eclipse E800 microscope (Nikon Corporation). Reactions were performed using three biological replicates (n=3).

Collagen type I and II deposition into the ECM of micromass cultures was semi-quantitatively determined using the colour deconvolution plug-in of Fiji software, as published by others (25). The empty parts of all images were set as white, and false-positive staining was excluded. During colour deconvolution, DAB labelling and haematoxylin nuclear staining were digitally isolated. Then, images were further analysed to obtain mean grey values, with arbitrarily set lower and upper thresholds. The mean grey values were converted into optical density values using step-tablet calibration. The relative amounts of deposited collagen types I and II were calculated as the average optical density values of DAB labelling normalised by the average optical density values of haematoxylin staining from four images per micromass cell culture. Values were normalised to day 6 values.

### Articular Cartilage Shavings

Full depth articular cartilage shavings were harvested from the knee joints of 35-day-old broilers as described in detail previously (26). Ethics license number: HB/15-ÉLB/03743-2/2022. Shavings were minced into small chips (∼1–2 mm^3^) using sterile scalpels, and 100 mg of cartilage chips was subjected to enzymatic digestion with 1.2% collagenase type II (Merck) in DMEM overnight at 37°C (26). Some shavings were routinely fixed, paraffin-embedded, sectioned, and stained with HE, acidic DMMB (3% acetic acid, pH 1.8), and neutral DMMB to visualise microscopic morphology.

### Total RNA Extraction

On the designated days of culture, HDCs were washed twice with physiological NaCl and then stored at ‒80 °C. Chondrocytes obtained from collagenase type II digested articular cartilage were collected by centrifugation at 5,000×*g* at 4°C for 10 min (26). For total RNA isolation, micromass cultures and pelleted articular chondrocytes were dissolved in TRI Reagent (Applied Biosystems, Foster City, CA, USA). Samples were mixed with 20% chloroform and centrifuged at 4 °C at 10,000 × *g* for 20 min. After incubation at −20 °C for 1 h in 500 μL of RNase-free isopropanol, the pellet of total RNA was dissolved in RNase-free water (Promega, Madison, WI, USA) and stored at −80 °C.

### RNA Sequencing

To obtain global transcriptome data, high-throughput mRNA sequencing analysis was performed using the Illumina sequencing platform. The quality of the total RNA samples was checked using an Agilent BioAnalyzer (Santa Clara, CA, USA) and the Eukaryotic Total RNA Nano Kit according to the manufacturer’s protocol (26). For micromass cultures, samples with an RNA integrity number (RIN) value of >7 were used for library preparation. Articular cartilage samples with an RIN > 6 were considered acceptable. RNA-seq libraries were prepared from total RNA using the Ultra II RNA Sample Prep Kit (New England BioLabs, Ipswitch, MA, USA), according to the manufacturer’s protocol. Briefly, poly-A RNAs were captured with oligo-dT conjugated magnetic beads then the mRNAs were eluted and fragmented at 94 °C. First-strand cDNA was generated by random priming reverse transcription and double-stranded cDNA was generated after the second-strand synthesis step. After repairing the ends, A-tailing, and adapter ligation steps, the adapter-ligated fragments were amplified by enrichment PCR, and sequencing libraries were generated. Sequencing runs were executed on an Illumina NextSeq500 instrument using single-end 75 sequencing cycles, generating an average of approximately 20 million raw reads per sample (26).

### RNA-seq Data Analysis

Raw sequencing data (fastq) were quantified using *Salmon* (27) and reads were aligned against a build of the Ensembl GRCg6a transcriptome, including decoy sequences, to improve accuracy. Subsequent bioinformatic analyses were conducted in R (version 4.1.1, except where stated otherwise). Reads were imported to *DESeq2* (28) with *tximport* (29) and normalised before all remaining analyses. Principal component analysis (PCA) was performed using the *prcomp* package and plotted with *ggplot2* (30) (along with all other bioinformatics plots, except where stated) and *ggforce* (31) packages. Differential gene expression was calculated for each contrasts indicated in Table 1 directly with *DESeq2*. The resulting differentially expressed genes (DEG) expression tables were limited by a Benjamini–Hochberg-adjusted false discovery rate (FDR) ≤ 0.05, and an LFC of ± 2. Heatmaps (*pheatmap*) were produced using only the top 80% of expressed genes across the samples and were additionally centred/standardised by column. Hierarchical clustering within the heatmaps was performed using the default *hclust* method. To calculate K-means clustering, DESeq2 normalised counts were row-centered by subtracting the median expression for each gene. K-means was calculated using the R-base function. For each cluster, STRING networks were calculated using the *STRINGDB* package (32), and cluster coefficients were calculated using *iGraph* (33). Gene ontology (GO) enrichment scores were calculated per cluster (and separately per day for DEGs) using the *Goseq* package (34).

**Table 1.** A. Summary of differentially expressed genes (DEGs) by comparing consecutive time points during chondrogenesis, as well as mature chondrocytes from articular cartilage of 35-day-old broilers. Total and unique entities (and their percentage) between pairwise comparisons, and their proportions are shown. **B.** An overview of the number of down- and upregulated genes between consecutive time points as well as mature chondrocytes, along with total and unique DEGs (and their percentage) between pairwise comparisons.**A.**

| Comparison | Total DEGs | Unique DEGs | Percentage |
| --- | --- | --- | --- |
| Day 0 vs. 1 | 590 | 279 | 47.29% |
| Day 1 vs. 2 | 42 | 13 | 30.95% |
| Day 2 vs. 3 | 44 | 8 | 18.18% |
| Day 3 vs. 4 | 14 | 4 | 28.57% |
| Day 4 vs. 6 | 42 | 10 | 23.81% |
| Day 6 vs. 10 | 550 | 241 | 43.82% |
| Day 10 vs. 15 | 408 | 109 | 26.72% |
| Day 15 vs. Mature | 2877 | 2305 | 80.12% |

**B.**
| Comparison | Total downregulated DEGs | Unique downregulated DEGs | Percentage |
| --- | --- | --- | --- |
| Day 0 vs. 1 | 132 | 101 | 76.52% |
| Day 1 vs. 2 | 1 | 1 | 100.00% |
| Day 2 vs. 3 | 6 | 4 | 66.67% |
| Day 3 vs. 4 | 2 | 1 | 50.00% |
| Day 4 vs. 6 | 22 | 10 | 45.45% |
| Day 6 vs. 10 | 264 | 192 | 72.73% |
| Day 10 vs. 15 | 324 | 264 | 81.48% |
| Day 15 vs. Mature | 1577 | 1502 | 95.24% |

| Comparison | Total upregulated DEGs | Unique upregulated DEGs | Percentage |
| --- | --- | --- | --- |
| Day 0 vs. 1 | 458 | 373 | 81.44% |
| Day 1 vs. 2 | 41 | 22 | 53.66% |
| Day 2 vs. 3 | 38 | 16 | 42.11% |
| Day 3 vs. 4 | 12 | 4 | 33.33% |
| Day 4 vs. 6 | 20 | 10 | 50.00% |
| Day 6 vs. 10 | 286 | 171 | 59.79% |
| Day 10 vs. 15 | 84 | 66 | 78.57% |
| Day 15 vs. Mature | 1300 | 1108 | 85.23% |

Enrichment bias was corrected by fitting a length/probability weight function to the gene length data for each gene, and over-representation *p*-values were then BH corrected for multiple comparisons. Gene lengths were obtained from the *tximport* object.

Trajectory analysis was conducted using *Monocle* 3 (which required R version 4.2) (35, 36). This tool was developed for single-cell RNA trajectory analysis but is also explicitly appropriate for a range of assays^2^. The DESeq2 normalised reads for each of the 27 samples were sliced to retain the top 10,000 genes (mean over the entire dataset), and genes with no known symbols were excluded. When there were duplicate symbols, only the first was retained. Samples were pre-processed with PCA and then clustered and ordered using Monocle 3 built-in routines, evoking uniform manifold approximation and projection (UMAP) dimension reduction. Pseudotime was inferred using *Monocle 3* with reverse graph embedding (37).

Transcription factors were identified by cross-referencing the expressed genes (top 50% of mean counts) to the go_id term GO:0003700 (and its children go_ids) using a combination of *BiomaRt* (38) and *Go.db* (39). The enrichment background/universe was defined as the complete set of genes expressed across all samples. The genes involved in chondrogenesis, cartilage development, and homeostasis were retrieved by aggregating the following GO terms: GO:1990079 cartilage homeostasis; GO:0051216 cartilage development; GO:0060536 cartilage morphogenesis; GO:0001502 cartilage condensation; and GO:0061975 articular cartilage development. LFC lists, by pairwise comparisons, were exported to IPA for identification of significant diseases and functions.

Co-expression data were analysed using the Weighted Gene Correlation Network Analysis (WGCNA) package (40). Our workflow followed that of Huynh *et al*. (6). The trait data *SOX9*, *ACAN,* and *COL2A* were derived from our transcriptomic datasets. The three most highly correlating modules (“*sienna3”*, “*thistle2”*, and “*pink*”) with 622 genes were aggregated and the top ∼500 genes were taken forward for further analysis and visualisation in Cytoscape (41).

The entire dataset has been deposited and published in the BioProject database (http://www.ncbi.nlm.nih.gov/bioproject/). BioProject IDs: PRJNA817177 (day 0–day 6 micromass cultures); PRJNA938813 (day 10 and day 15 micromass cultures; mature articular chondrocytes).

### Transient Gene Silencing

The effects of transient gene silencing of *ATOH8* and *EBF1* on chondrogenic differentiation were studied using siRNA electroporation of freshly isolated LMPs. siRNAs for *ATOH8* (Silencer™ Select, Design ID #: ABMSIJ9) and *EFB1* (Silencer™ Select, Design ID #: ABI1N1L), as well as a non-targeting control (Silencer™ Select Negative Control No. 1 siRNA; Cat. no. 4390843), were custom-designed (except the non-targeting control) and purchased (Life Technologies, Thermo Fisher Scientific). Electroporation was performed as previously described (42), with modifications to the original protocol. We generated a single-cell suspension of LMPs, as described in the ‘Cell Cultures’ section. Instead of Ham’s F12 culture medium, the cells were resuspended in a modified sucrose electroporation buffer prepared by adding 272 mM sucrose (Sigma-Aldrich, Merck) to Opti-MEM™ Reduced Serum Medium (Life Technologies, Thermo Fisher). siRNA (2 μg) was added to the cell suspension and mixed prior to transferring the entire volume to a 2.0 mm gap sterile electroporation cuvette (Cat. no. 732-1136, VWR, Avantor Sciences). A BTX ECM 830 Square Wave Electroporation System (Harvard Apparatus) was used to pulse the LMPs with the following settings: 3–400 V pulses 150 µs in length at 100 ms intervals. Following electroporation, cells were incubated at 4°C for 10 min and gradually warmed to allow them to sink to the bottom of the cuvette. The lower four-fifths of the suspension were removed using a micropipette and pelleted by centrifugation. Cells were then counted and resuspended at a density of 1.5 × 10^7^ cells/mL in 37°C Ham’s F12. 30- μL droplets of the cell suspension were inoculated onto the surface of 30-mm round cover slips (3 × 30-μL droplets) (Menzel-Gläser) placed in 35-mm Petri dishes (Eppendorf). As described above, the cultures were then kept in the incubator for 2 h prior to flooding with F12 medium. The culture medium was changed every two days.

For the evaluation of cartilage ECM production, colonies were fixed on culture day 6, stained with DMMB (pH 1.8), photographed, and analysed using a MATLAB script, as described above. Values were normalised to non-targeting control values. Relative cell numbers (inferred from mitochondrial activity) were analysed by MTT assay, as previously reported (16). Briefly, MTT reagent (5 mg/1 mL PBS) was added to the cells cultured in 24-well plates on culture day 6. Cells were incubated for 2 h at 37°C, and following the addition of MTT solubilizing solution, optical density was measured at 570 nm (Chameleon, Hidex, Turku, Finland). Optical density readings of the experimental groups (*ATOH8* and *EBF1* gene-silenced cultures) were normalised to those of the non-targeting controls and are shown as percentage changes.

Gene expression studies of osteo/chondrogenic marker genes were carried out as described previously (43) with the following modifications. Cultures were harvested 2, 4, or 6 days post-electroporation. Total RNA was isolated from micromass cultures using the TRI Reagent (Applied Biosystems) according to the manufacturer’s instructions. Total RNA (1 μg) was reverse-transcribed into complementary cDNA using the High-Capacity cDNA Reverse Transcription Kit (Thermo Fisher Scientific) according to the manufacturer’s protocol. The expression patterns of osteo/chondrogenic marker genes were determined by RT-qPCR and relative quantification using the 2^–ΔΔCt^ method. Primer pairs were obtained from Integrated DNA Technologies (Coralville, IA, USA). For the sequences of the custom-designed primer pairs, please see Table S1 in Supplementary file 1. SYBR Green-based RT-qPCR reactions were set up using GoTaq qPCR Master Mix (Promega) and 20 ng of input cDNA per each 10-μL reaction. Reactions were run on a QuantStudio 3 Real-Time PCR System (Thermo Fisher Scientific) using the following standard thermal profile: activation and initial denaturation at 95°C for 2 min, followed by 40 cycles of denaturation at 95°C for 3 s, annealing and extension at 60°C for 30 s, and then final extension at 72°C for 20 s. Data were collected during the extension step. Amplification was followed by a melt curve stage analysis (denaturation at 95°C for 15 s, annealing at 55°C for 1 min, dissociation at 0.15°C/s increments between 55°C and 95°C). Data were collected at each increment during the dissociation. Amplification data were analysed using the QuantStudio Design and Analysis Software (version 1.5.1), and exported data were processed using Microsoft Excel (version 2108). Six reference genes were analysed for stability at each time point as follows: hypoxanthine phosphoribosyltransferase 1 (*HPRT1*), peptidylprolyl isomerase A (*PPIA*), ribosomal protein L4 (*RPL4*), 60S ribosomal protein L13 (*RPL13*), ribosomal protein S7 (*RPS7*), and tyrosine 3-monooxygenase/tryptophan 5- monooxygenase activation protein zeta (*YWHAZ*). BestKeeper (44), a Microsoft Excel-based tool, was used to determine the optimal normalising gene among the chosen reference genes based on pairwise correlations. Real-time qPCR data for each gene of interest were normalised to *RPS7*, the most stable reference gene.

## RESULTS

### Global Gene Expression Patterns During Chondrogenesis Indicate Progressive Changes in the Transcriptome

To identify the genes involved in chondrogenic differentiation in limb bud-derived LMPs, we performed global transcriptomic analysis of micromass cultures at eight different time points, representing the key stages of the process *in vitro* (see the workflow in Fig. 1). Articular cartilage obtained from knee joints of 35-day-old broilers was also included in the analysis. Chondrogenic differentiation of the micromass cultures was confirmed by the increasing metachromatic staining pattern of the cartilage ECM, as revealed by DMMB, AB, and Safranin-O staining (Fig. 2), and the reduced proliferation rate of the cells in the micromass cultures over time (see Fig. S1 in Supplementary file 1). Cartilage matrix production was also monitored by immunohistochemical analysis of collagen types I and II deposition (Fig. 3), showing a gradual increase in collagen incorporation into the ECM starting from day 4, especially in the case of collagen type II.

**Figure 2.**
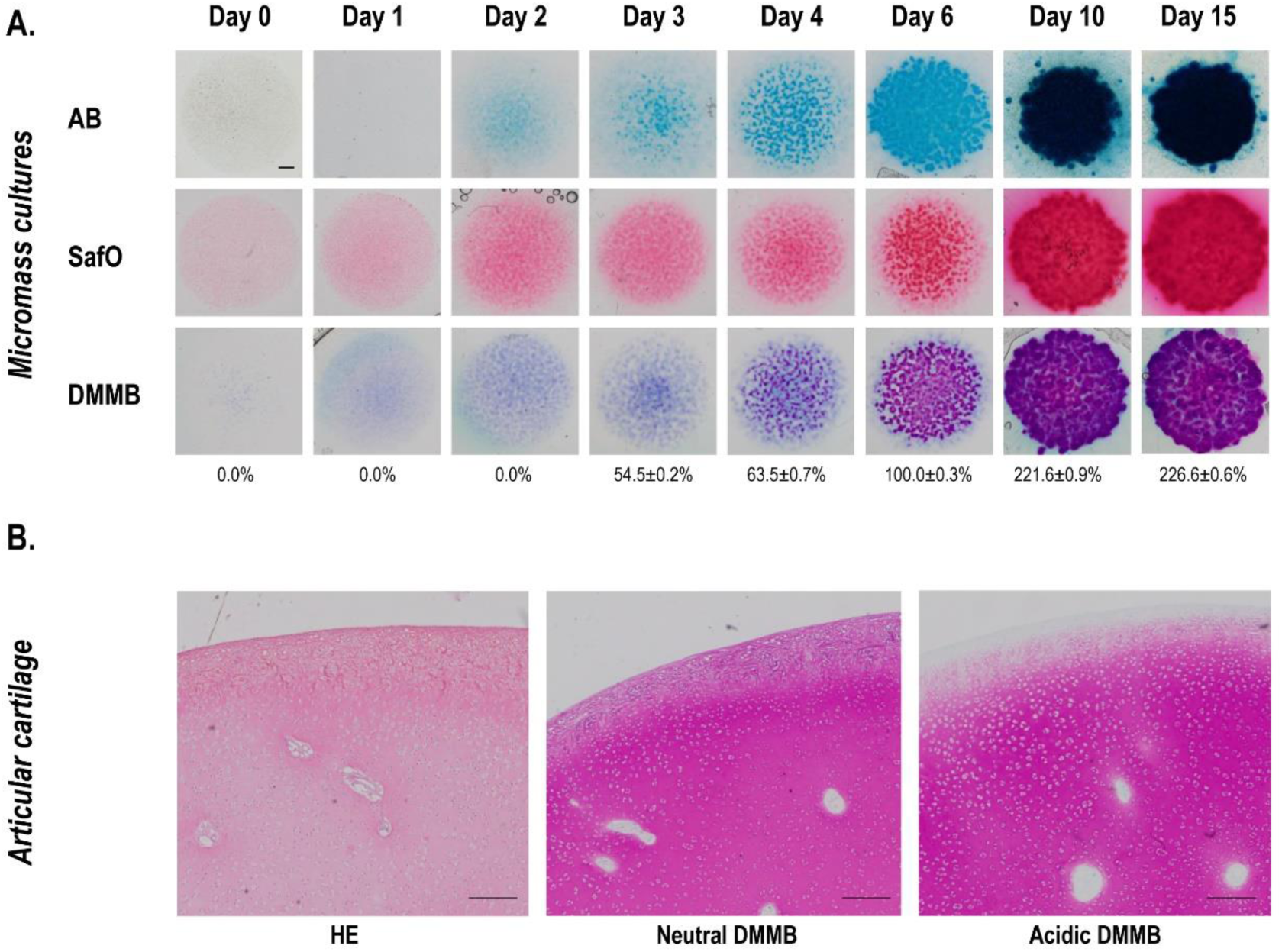
Light microscopy analysis of micromass cultures during the course of chondrogenic differentiation (days 0–15), and articular cartilage obtained from the knee joint of 35-day-old broilers. (**A**) Photomicrographs of alcian blue (AB), safranin-O (SafO) and dimethyl methylene blue (DMMB) stained cultures are shown. Original magnification was ×4. Scale bar, 1 mm. Values below images of DMMB stained cultures reflect results obtained using a MATLAB-based image analysis of metachromatic areas. Data are expressed as mean ± SEM, compared to day 6 (100%). (**B**) Photomicrographs of histological sections of chicken articular cartilage following haematoxylin-eosin (HE), neutral DMMB, and acidic (pH 1.8) DMMB staining. Original magnification was ×4. Scale bar, 1 mm. Representative data out of 3 biological replicates.

**Figure 3.**
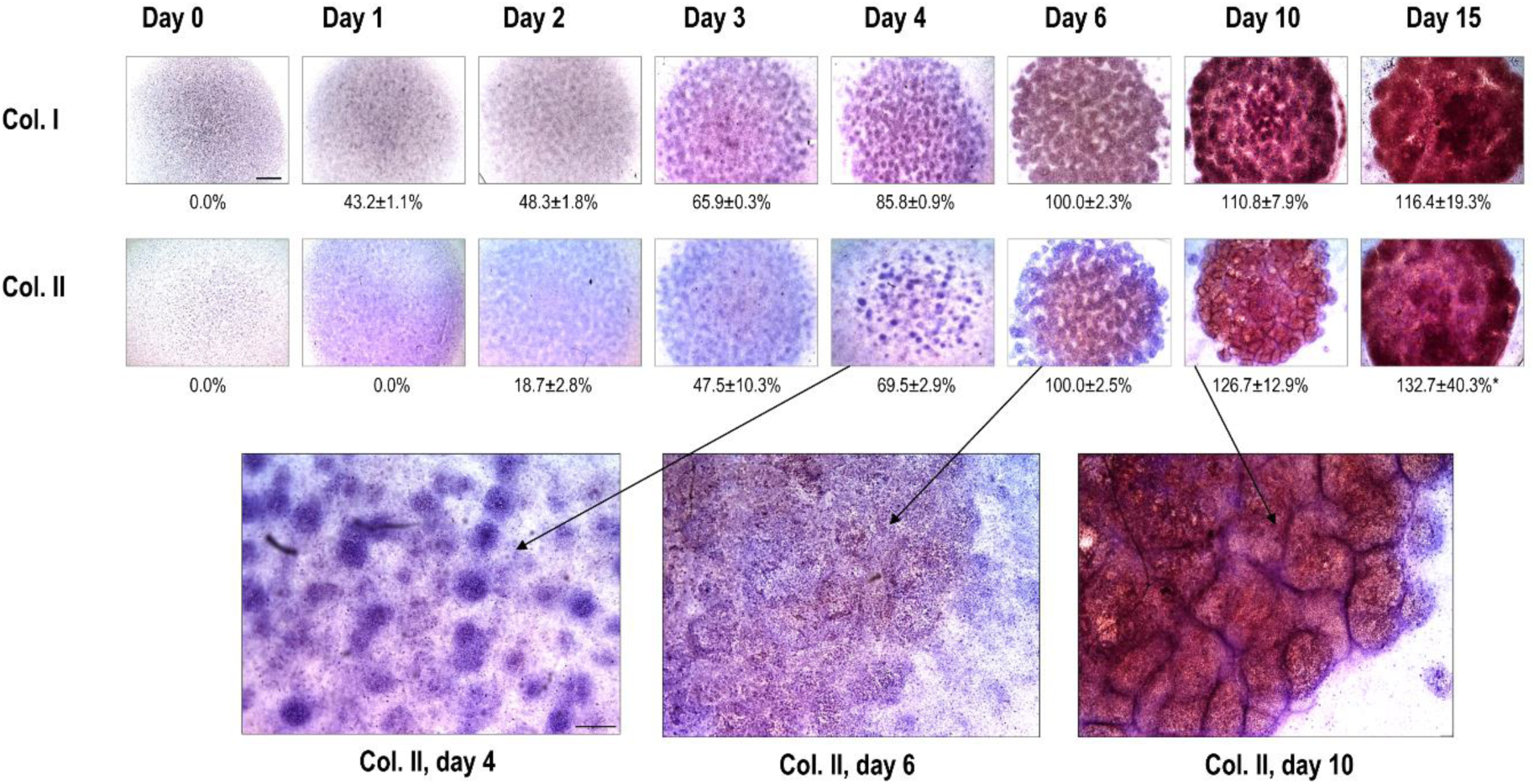
Collagen type I and II immunohistochemistry visualised by DAB chromogen reaction in chondrifying micromass cultures of various ages. Scale bars, 500 μm (thumbnail images in the upper panel) or 200 μm (inserts in the lower panel for more mature (*i.e*., day 6, 10, and 15) cultures). The brown deposits represent immunopositive signals for collagen type I or II, respectively, within the chondrogenic nodules of the micromass cultures. Values below images reflect the image analysis of immunopositive areas, normalised to haematoxylin nuclear staining. Data are expressed as mean ± SEM, compared to day 6 (100%). Representative photomicrographs are shown out of 3 biological replicate experiments.

Having confirmed chondrogenic differentiation and metachromatic ECM deposition in the micromass cultures, we performed PCA of the RNA-seq data to explore their interrelationships and visualise the correlation between the samples (Fig. 4A–B). Biological replicates within different time points were clustered together, showing marked changes in the transcriptome over time in culture, as the chondrogenic cells underwent spontaneous chondrogenesis in the micromass cultures. The transcriptomes of mature chondrocytes obtained from articular cartilage clustered together and were distinct from those of the micromasses. There was an overall trend from less mature (undifferentiated) cells on day 0 to more mature (differentiated) micromass cultures on day 10 and towards mature articular chondrocytes, indicating a progressively different transcriptome during chondrogenesis. The first component (PC1) on the *x*-axis accounted for 52.46% of the variance. There was a gradual shift along the *x*-axis as chondroprogenitor cells differentiated into chondroblasts and chondrocytes in the micromasses (between days 0 and 10), with articular chondrocytes positioned at the far end. However, the day 15 samples seem to have lost this trend, and it seems that these cultures are positioned backward and closer to the immature cultures. The second and third components (PC2 and PC3) on the *y*-axis explain 15.84% and 11.93% of the variance, respectively (Fig. 4A–B). Along the hyperplanes of the first *versus* third principal components, samples obtained on day 15 (*i.e.,* cells in more mature micromass cultures) clustered separately from the other sample groups, again closer to less mature cultures. Mature articular chondrocytes clustered separately from micromasses, indicating differences in their transcriptomes. The slightly greater scatter on days 6, 10, and 15 along the *y*-axis (Fig. 4A) may indicate that the biological replicates were slightly out of phase at this stage of chondrogenic differentiation, resulting in greater inter-culture and inter-cellular variability. However, PCA demonstrated that the variance between time points was generally still greater than the variance between biological replicates.

**Figure 4.**
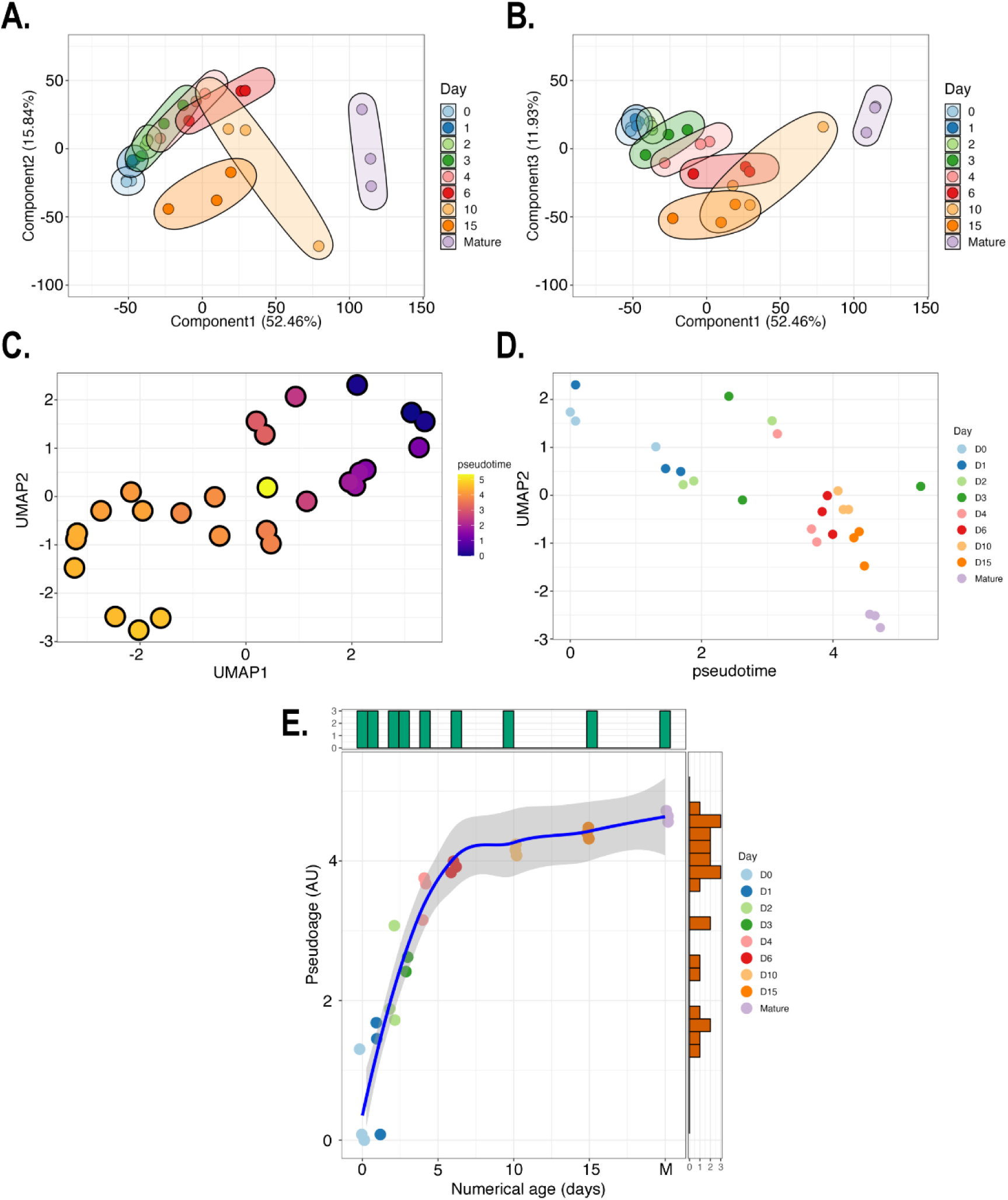
Low-dimensional space representation, including principal component analysis (PCA; panels **A–B**) and uniform manifold approximation and projection (UMAP; panels **C–E**), of the RNA-seq data. PCA was performed on the normalised expression data. The first component (PC1) explains 52.46% of variability, the second component (PC2) explains 15.84% (**A**), and the third component (PC3) explains 11.93% (**B**). Each point represents an experimental sample; colours indicate different time points (days) of chondrogenesis, or mature articular cartilage. Samples clustered together by time points (culturing days/maturity), and there was minor variation between biological replicates, except for day 10. Following UMAP dimension reduction, RNA-seq data were ordered with reverse graph embedding and pseudotime calculated for each culture of known latent development time (**C**). Pseudotime plotted against UMAP component 1, showing that cultures progress systematically through pseudotime, but there is little separation once they reach 4.5 (**D**). Numerical time *vs*. pseudotime plot (M, mature articular chondrocytes) shows the (pseudotime) developmental saturation occurring from approximately day 6 (**E**).

To investigate whether the transcriptome progresses through a distinct developmental trajectory throughout the entire dataset, we used the computational approach of trajectory analysis using the *Monocle 3* R package. Following UMAP dimension reduction (Fig. 4C), micromass culture sequence data were ordered using reverse graph embedding (37), and pseudotime was calculated for each culture of known latent development time (using *Monocle3*, see Methods). Fig. 4D shows the plot of pseudotime against UMAP component 1, and it is clear that cultures progress systematically through pseudotime, but there is little separation once they reach 4 (arbitrary units). A comparison of the numerical time *vs*. pseudotime plot graphically shows that (pseudotime) developmental saturation occurred from approximately day 6 (Fig. 4E).

We then generated heatmaps based on the normalised expression data. Hierarchical clustering based on the global transcriptome expressed in chondrogenic micromass cultures and mature articular chondrocytes also confirmed the segregation of samples, primarily according to the age of the cultures (Fig. 5A). The transcriptomic signatures in samples from the early stages of chondrogenesis (days 0, 1, and 2) clustered effectively between biological replicates and between different time points. The transcriptomes of chondrocytes from mature articular cartilage also clustered together, and they shared a considerable overlap with day 10 and day 15 cultures, representing mature and probably hypertrophic colonies. Similarly, colonies from culturing days 3, 4, and 6, predominantly containing differentiating chondroblasts and chondrocytes, were not characterised by distinct profiles and clustered together, indicating that differentiating chondroblasts and chondrocytes shared a very similar transcriptome on these culturing days. When we compared all time points to day 0, the fold changes (both upregulated and downregulated genes) gradually increased (Fig. 5B), indicating that the transcriptome of more differentiated cultures gradually became more distinct than that of undifferentiated (day 0) colonies. The transcriptome of adult chondrocytes was distinct and showed considerable similarity to that of the most mature (day 10 and 15) micromasses.

**Figure 5.**
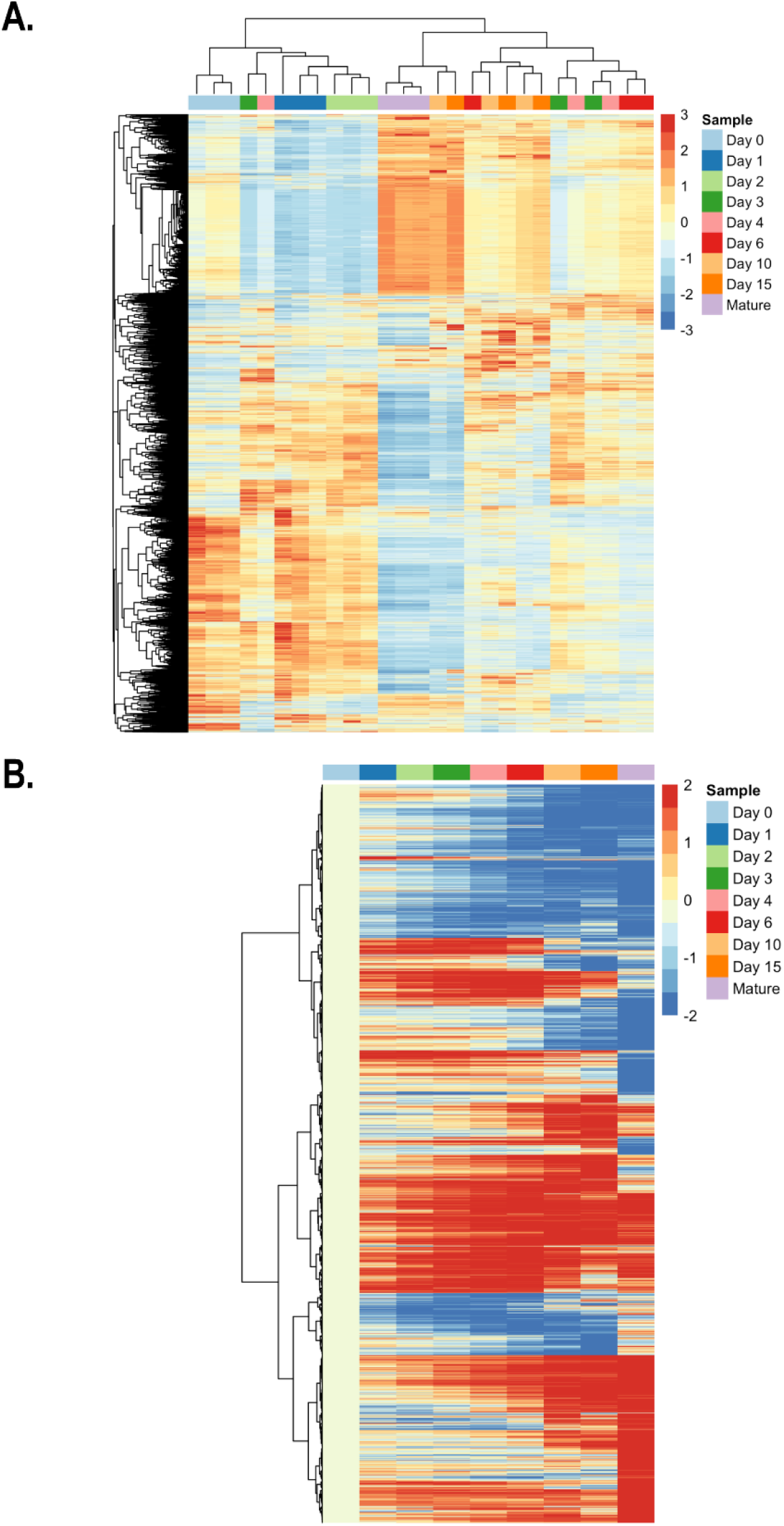
Hierarchical clustering (column dendrogram) of the global transcriptome of chondrogenic cells during the differentiation programme (days 0-15) and that of mature articular chondrocytes (**A**). Data for all 3 biological replicates are shown. Whilst most time points clustered separately (day 0, 1, 2 micromasses, and mature chondrocytes from 35-day-old broilers), days 3, 4, and 6, as well as days 10 and 15 clustered together, indicating overlapping transcriptomes at these time points. Heatmap showing differentially expressed genes (DEGs) normalised to day 0 (**B**). Average expression between biological replicates was used for calculating log2 fold change (LFC) values. A gradual increase in up and downregulation was observed during the course of differentiation towards mature articular chondrocytes.

### The Top 20 Most Abundantly Expressed and Differentially Expressed Genes on Each Culturing Day Include Transcripts Known to be Involved in Chondrogenic Differentiation

The top 20 most abundantly expressed genes for each culture day are listed in Supporting Information (Table S2 in Supplementary file 1). Genes involved in chondrogenesis and cartilage ECM synthesis (*COL1A1*, *COL1A2*, *COL2A1*, *COL9A1*, *COL11A1*, *FN1*, *ACAN*), cytoskeleton (*ACTB*, *TUBA1A1*, *TUBB*), ribosomal proteins and RNAs (*NCL*, *RPS2*, *RPS3A*, *RPS6*, *RPS12*),glycolysis and energy metabolism (*GAPDH*, *ENO1*), transcription factors (*YBX1*), translation initiation and elongation factors (*EIF4G2*, *EEF1A1*), ribonucleoproteins (*HNRNPA1*, *HNRNPA3*), heat shock factors (*HSP90AA1*, *HSPA2*, *HSPA5*, *HSPA8*, *HSPB9*), and lncRNA transcripts (*LOC776816*, *LOC101747587*) were enriched in the list for each culture day.

Notably, the proportion of genes involved in cartilage development (GO:0051216) gradually increased until day 10 but then declined again in day 15 micromass cultures. Unsurprisingly, the highest proportion of genes involved in cartilage development and homeostasis was observed in the mature articular chondrocytes (Table S2).

Many genes were found to be significantly differentially regulated, as revealed by comparing the transcriptomic profiles of consecutive culture days. In total, there are 3470 DEGs in this dataset. An overview of the downregulated and upregulated genes for each comparison, and the number of genes that were unique for each comparison between consecutive days, is shown in Table 1. The top 20 DEGs obtained from pairwise comparisons between consecutive culture days (*i.e*., day 0 *vs*. 1, day 1 *vs*. 2, etc.), ranked according to their LFC values, are shown in Table 2. Full lists of DEGs between consecutive culturing days are shown in the Supporting Information (Supplementary file 2 – ‘*DEG*-*full-list’ worksheet*). The number of DEGs gradually declined between pairwise comparisons of consecutive culture days until days 3, 4, and 6, indicating a very high abundance of genes modulated early during chondrogenic differentiation. For example, while 590 genes were differentially regulated between days 0 and 1, only 14 were differentially expressed between days 3 *vs*. 4. Again, there were more DEGs between pairwise comparisons of the later stages of chondrogenesis, and the number of modulated genes was very high between 15-day-old micromasses and mature chondrocytes (Table 1A). It is also worth highlighting that while most DEGs were upregulated during the early stages of chondrogenesis (458 upregulated *vs*. 132 downregulated DEGs between days 0 and 1), the trend was reversed in more mature micromasses (84 upregulated *vs*. 324 downregulated DEGs between days 10 and 15) (Table 1B).

**Table 2.**
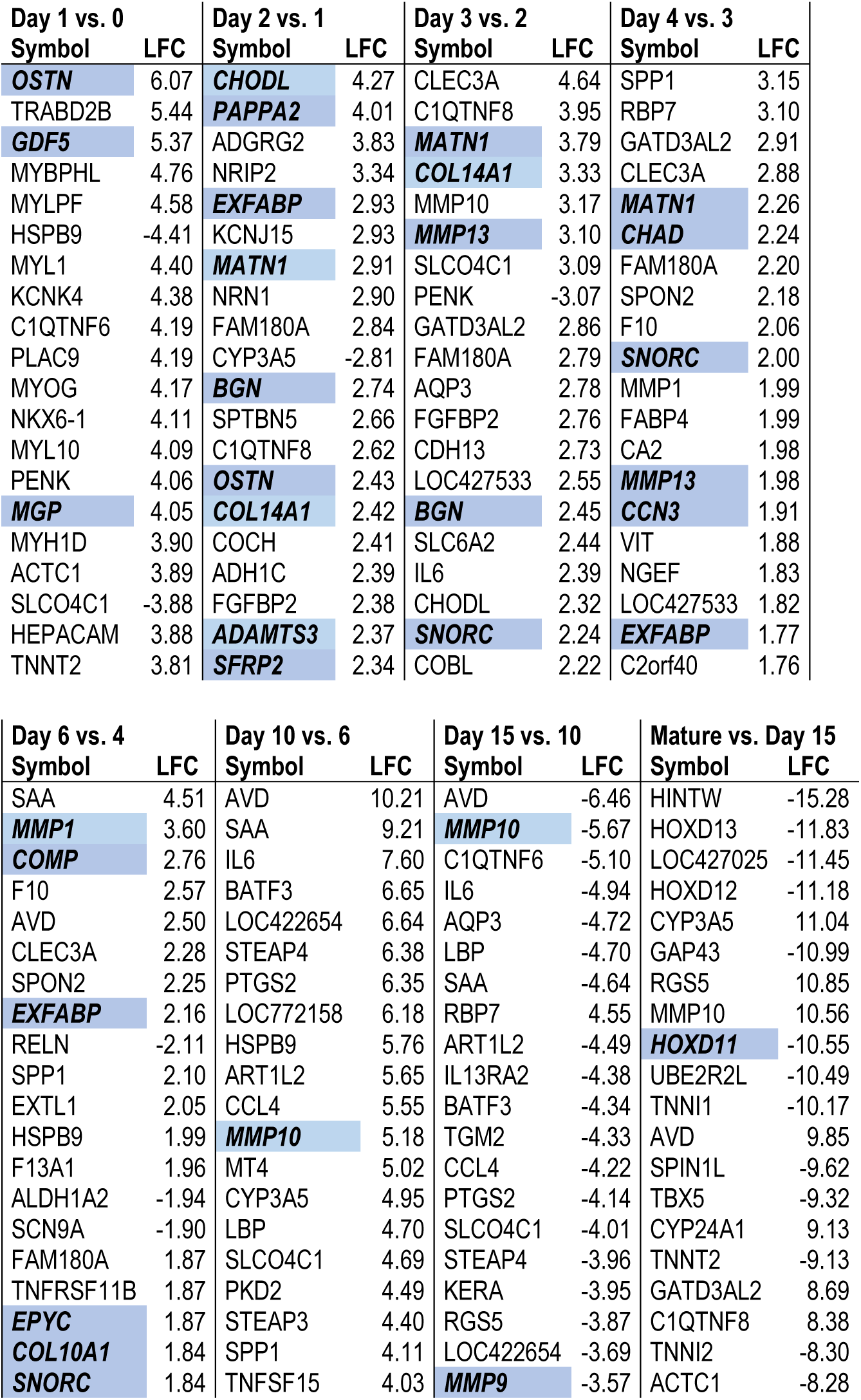
Top 20 significant DEGs in pairwise comparisons between different time points during *in vitro* chondrogenesis, ranked according to log2 fold change (LFC) values. LFC cut-off: 2.0. Gene name in **bold** over a blue background have a known role during cartilage development (GO:0051216).

In line with the bulk analysis of the differential expression of the transcriptome, several genes involved in chondrogenesis (GO:0051216) were among the top 20 DEGs between pairwise comparisons of consecutive culture days, such as *GFD5*, *MATN1*, and *COMP* (Table 2). Cartilage development-associated genes were over-represented among the top 20 DEGs until day six.

### The Top 1000 Most Abundantly Expressed Genes on Each Day Form a Network of Interconnected Entities

We exported the top 1000 most abundant transcripts at each time point, including mature articular chondrocytes, to Cytoscape and analysed the data using the STRING plugin. The analysis results (including the nodes, interactions, expected interactions, and cluster coefficients) are summarised in Table 3. Interestingly, day 0 was characterised by the highest number of interactions, followed by a steady decline in this parameter during chondrogenic differentiation (until day 6), whereas the trend was reversed in more mature micromasses and articular chondrocytes. On each day of culture, transcripts with chaperones (*CCT2, HSP90AA1, HSP90AB1, HSPA4, HAPS5, HSPA8*), ribosomal (*RPLP0*, *RPS27A*, *RP3*), translational (*EEF1A1, EEF2, EFTUD2*), cytoskeletal (*ACTB*), ubiquitination (*UBA52*), and glycolytic (*GAPDH*) functions and catenin beta involved in adherens junctions (*CTNNB1*) had the highest number of connections and closeness centrality values. In mature chondrocytes, mitogen-activated protein kinase 3 (*MAPK3*), interleukin-6 (*IL6*), prolyl 4-hydroxylase subunit beta (*P4HB*), and phosphatase and tensin homolog (*PTEN*) were among the those with the highest closeness centrality values.

**Table 3.**
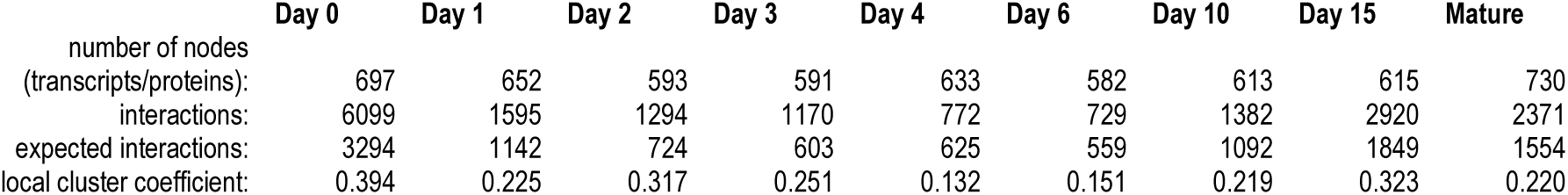
The overall analysis of nodes and edges of networks amongst the top 1000 most abundant transcripts at each time point using the STRING plugin of Cytoscape.

### GO and IPA Network Analysis of DEGs Revealed Pathways Related to Chondrogenic Differentiation

*Gene Ontology (GO) analysis.* By analysing the most abundant transcripts as well as the most highly regulated DEGs between the different time points, we identified genes that were already known to be involved in chondrogenesis and/or expressed in chondrocytes according to the literature (see Table 2 and Table S2 in Supplementary file 1). We then analysed in more detail those GO terms (focusing on molecular function and biological pathway categories) that were over-represented in differentially expressed sets of genes between consecutive time points. The DEGs between days 0 and 1, mainly containing undifferentiated chondrogenic LMPs, were enriched with general terms such as “tissue development,” “system development,” “developmental process,” and “anatomical structure development” (Fig. 6A), which correlate with the onset of cartilage formation. While there were no significantly enriched GO terms in DEGs between days 1, 2, 3, 4, and 6 after Benjamini-Hochberg correction, more specific terms such as “cartilage development,” “connective tissue development,” “positive regulation of cartilage development,” and “collagen binding” were still ranked forward in these comparisons. The major enriched pathways in DEGs between days 6 and 10 included terms related to signal transduction pathways, “growth factor activity,” and “cytokine activity,” but more notable terms such as “positive regulation of cartilage development” were also enriched (Fig. 6B). When we looked at enriched GO terms for DEGs between day 10 and day 15 micromasses, pathways related to cartilage ECM organisation and cell surface receptor signalling were significantly over-represented, with terms such as “skeletal system development,” “extracellular matrix structural constituent” (Fig. 6C). When we observed a subset of genes significantly differentially regulated between day 15 micromasses and articular chondrocytes, the following GO terms were over-represented: “G protein-coupled receptor signaling pathway,” “anatomical structure development,” “anatomical structure formation involved in morphogenesis,” “animal organ morphogenesis,” “tissue development,” “extracellular matrix structural constituent,” and “skeletal system development,” all relevant to cartilage formation (Fig 6D). We also studied DEGs between the earliest (day 0) and latest stage (day 15) micromass cultures and found the following relevant GO terms to be enriched: “cartilage development,” “extracellular matrix organization,” “tissue development,” “extracellular matrix structural constituent,” and “collagen fibril organization” (Fig. 6E).

**Figure 6.**
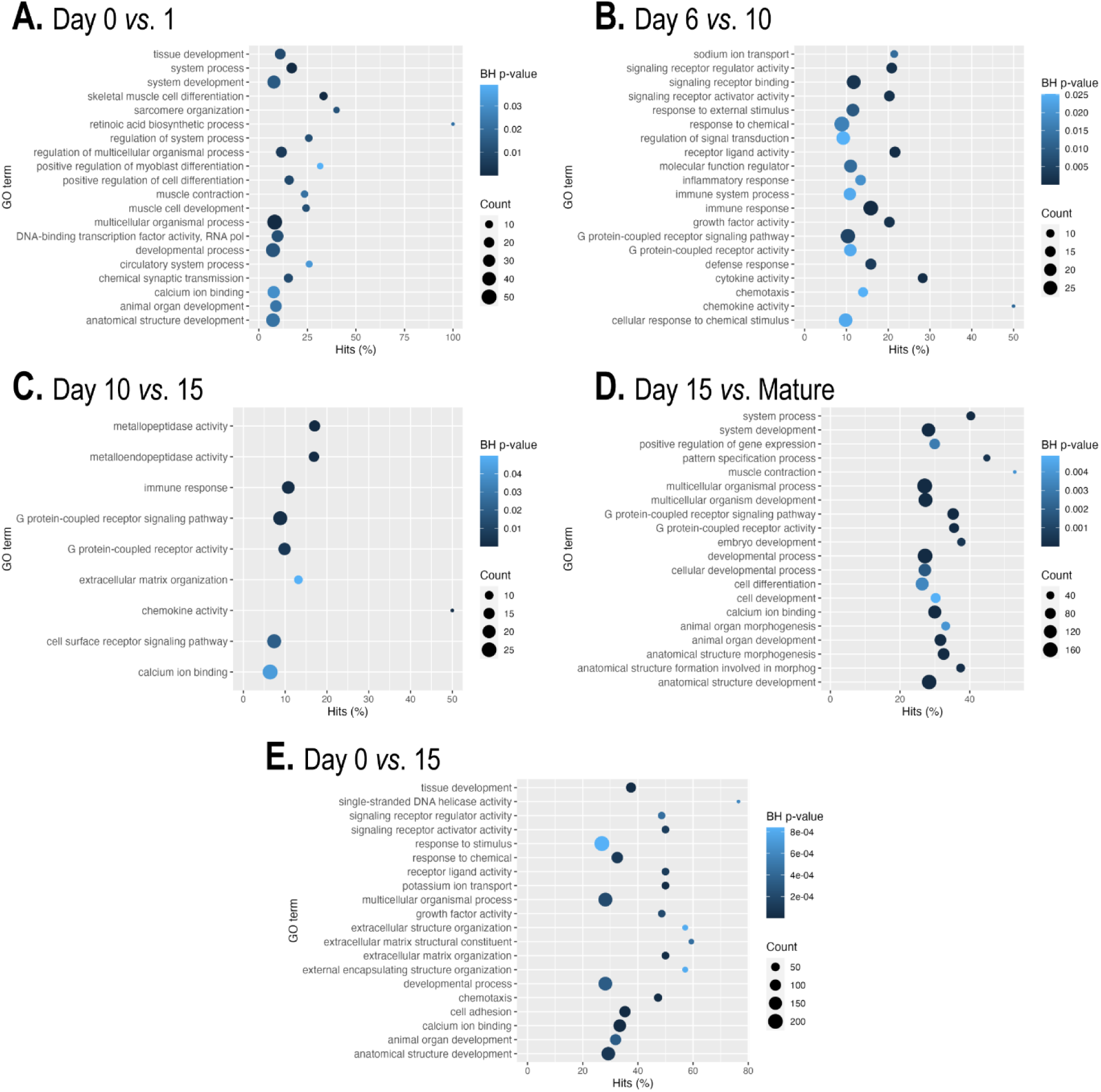
Significant GO term (molecular function, biological process) enrichment (Benjamini-Hochberg (BH) adjusted *p*-value) in DEGs between pairwise comparisons of consecutive culturing days and mature articular chondrocytes (**A–E**). If not shown, there are no significantly enriched GO categories in that comparison.

*IPA analysis.* We also submitted DEGs (LFC > ±2) from pairwise comparisons of time points to the Qiagen IPA knowledgebase and analysed the top significant diseases and functions. The overrepresented networks are summarised in the Supporting Information (Table S3 in Supplementary file 1). When we compared the early time points (day 0 vs. day 1, and day 1 vs. day 2), the following networks were identified: “Embryonic Development, Organismal Development, Tissue Development,” “Cancer, Cell-To-Cell Signaling and Interaction, Organismal Injury and Abnormalities,” “Amino Acid Metabolism, Cell-To-Cell Signaling and Interaction, Molecular Transport,” “Cardiovascular Disease, Hereditary Disorder, Skeletal and Muscular System Development and Function,” and “Connective Tissue Disorders, Organismal Injury and Abnormalities, Skeletal and Muscular Disorders”.

When comparing the DEGs between days 2 and 3, the following networks were over-represented: “Cell to Cell Signaling and Interaction, Cellular Growth and Proliferation, Hematological System Development and Function,” “Connective Tissue Development and Function. Nervous System Development and Function, Reproductive System Development and Function,” and “Connective Tissue Disorders, Developmental Disorder, Hereditary Disorder.”

In addition to the more generic terms, the terms “Connective Tissue Disorders, Protein Synthesis, RNA Post-Transcriptional Modification,” “Connective Tissue Disorders, Developmental Disorder, Hereditary Disorder,” and “Cell Cycle, Connective Tissue Development and Function, DNA Replication, Recombination, and Repair,” which are more closely related to chondrogenesis, were identified in the day 2 *vs*. day 3, day 3 *vs*. day 4, and day 4 *vs*. day 6 comparisons.

The terms “Cell to Cell Signaling and Interaction, Cellular Movement, Hematological System Development and Function” and “Cell to Cell Signaling and Interaction, Drug Metabolism, Molecular Transport” were present in the comparisons at the later time points, indicating that the genes involved in mediating intracellular signalling pathways were enriched. In a similar way, terms such as “Cellular Development, Cellular Movement, Hepatic System Development and Function,” “Cell Death and Survival, Cellular Growth and Proliferation, Gastrointestinal Disease,” and “Connective Tissue Disorders, Organismal Injury and Abnormalities, Skeletal and Muscular Disorders” were enriched in the comparisons between the more mature stages of the chondrogenic cultures. When we compared 15-day-old micromass cultures with mature chondrocytes, the following terms were enriched: “Cancer, Cell-to Cell Signaling and Interaction, Nervous System Development and Function” and “Cellular Development, Cellular Growth and Proliferation, Developmental Disorder.”

### Unsupervised K-Means Clustering Analysis

To identify gene clusters according to their expression dynamics, six groups were defined using the K-means algorithm (Fig. 7). The gene lists for each cluster are available in the Supporting Information (Supplementary file 2 – ‘C*lusters’ worksheet*). A detailed analysis of each cluster using the STRING plugin in Cytoscape (nodes, interactions, expected interactions, and cluster coefficients) is presented in Table 4.

**Figure 7.**
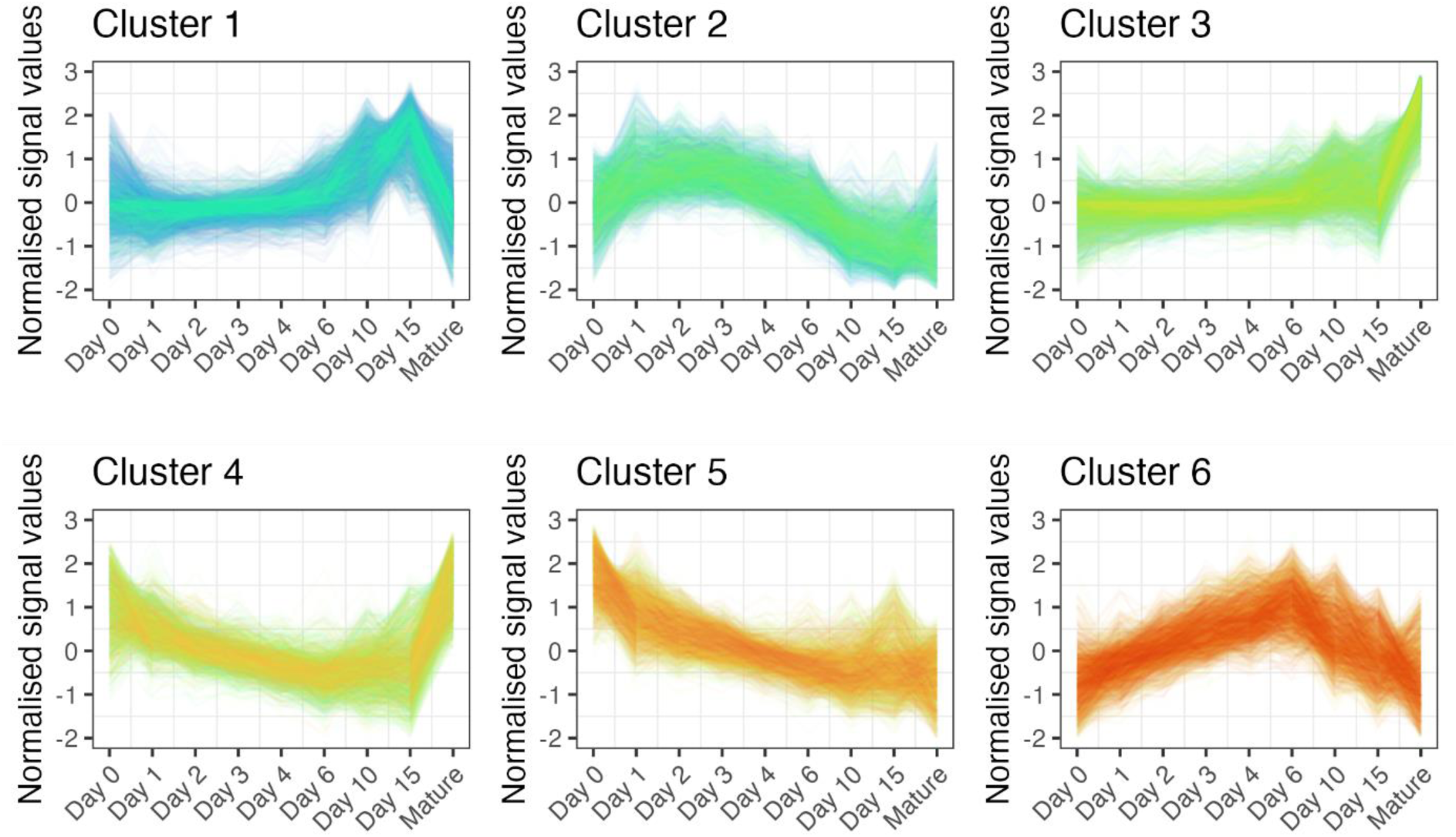
Unsupervised clustering analysis of genes, based on their normalised expression values (“Signal”) during chondrogenesis using the K-means algorithm, defined 6 groups of genes. The expression dynamics of each cluster are visible in clusters 1–6. *x*-axis, time points (days of culturing).

**Table 4.** The overall analysis of nodes and edges of networks in each unsupervised cluster of transcripts using the STRING plugin of Cytoscape.

|  | Cluster 1 | Cluster 2 | Cluster 3 | Cluster 4 | Cluster 5 | Cluster 6 |
| --- | --- | --- | --- | --- | --- | --- |
| number of nodes (transcripts/proteins): | 1824 | 1381 | 1741 | 987 | 1444 | 1104 |
| interactions: | 18,396 | 4861 | 11,781 | 4824 | 16,500 | 2836 |
| expected interactions: | 14,443 | 3828 | 8563 | 3920 | 10,699 | 2418 |
| local cluster coefficient: | 0.267 | 0.183 | 0.209 | 0.228 | 0.333 | 0.156 |

Genes in **cluster 1** were characterised by flat but gradually increasing expression levels over time with peak expression on days 10 and 15, followed by a massive downregulation in mature articular chondrocytes (Fig. 7). There were 2731 genes in this cluster, making it the largest group. The over-represented GO terms in this cluster were related to RNA processing,

RNA binding, ribosome biogenesis, gene expression, transcription, and translation, as well as ‘cellular metabolic process,’ ‘lipid biosynthetic process,’ ‘carboxylic acid metabolic process,’ and ‘oxoacid metabolic process’ (Fig. 8). Despite the lack of GO terms relevant to cartilage and chondrocytes, this cluster included 46 genes relevant to chondrogenic differentiation, including *CCN2*, *FGF18*, *FGFR3*, *RUNX2*, *TGFB1* and *TGFB2*, *WISP2* (*CCN5*). The key nodes with the highest closeness centrality values in this group were chaperones (*HSP90AA1*, *HSPA4*, *HSPA8*), *GAPDH*, *MYC*, ribosomal proteins (*RPLs* and *RPSs*), and *HDAC1*.

**Figure 8.**
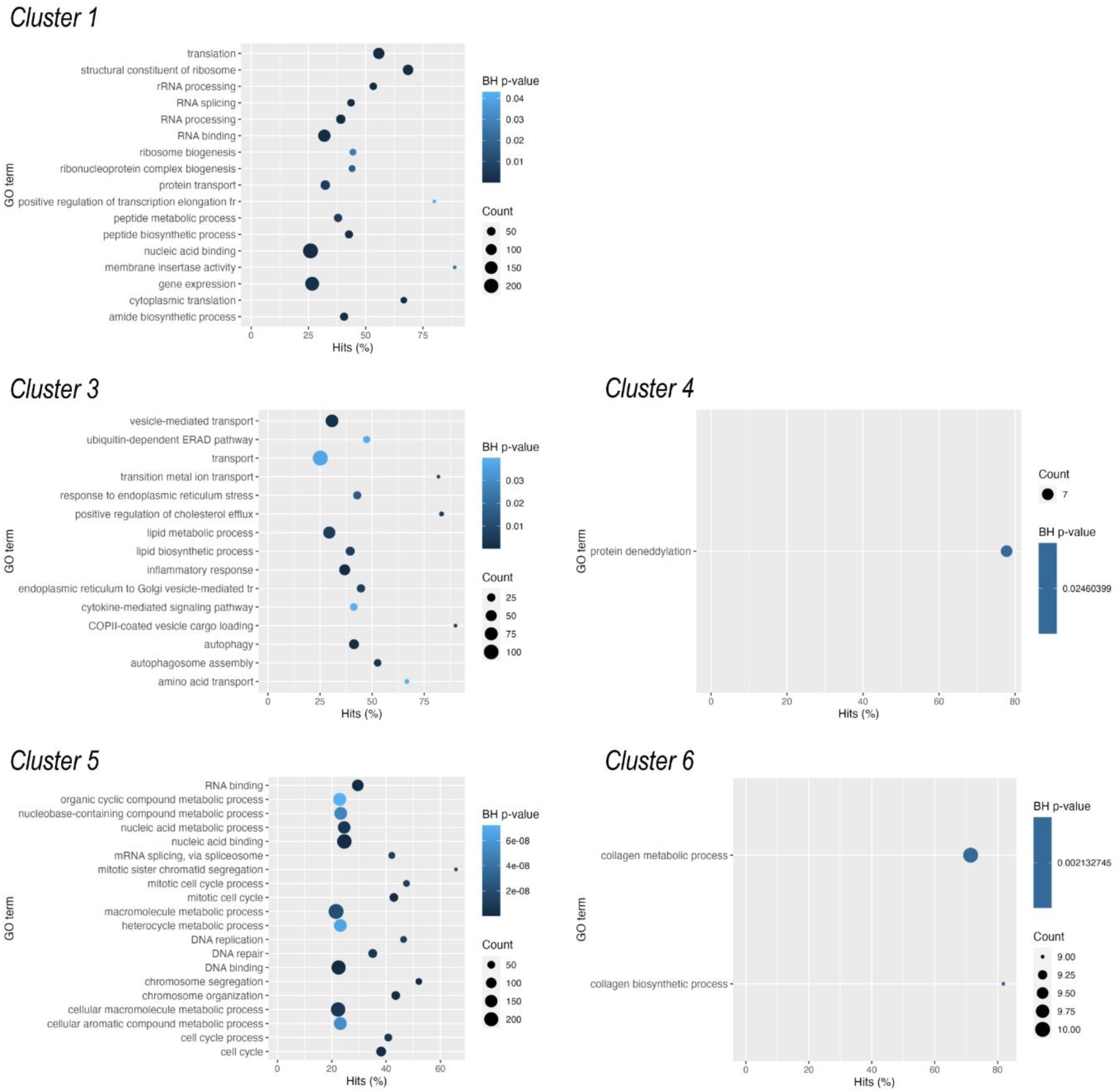
Significantly (Benjamini-Hochberg (BH) adjusted *p*-value) over-represented GO terms (molecular function, biological process) for the 6 clusters of genes defined by K-means algorithm. There were no significantly enriched GO categories in cluster 2.

**Cluster 2** contained 1986 genes. The genes in this cluster were characterised by a low expression level on day 0, and there was a trend of upregulation at early time points (days 1, 2, 3, and 4), followed by downregulation at later time points (day 10 and 15 micromasses, and mature chondrocytes; Fig. 7). There were no significantly enriched GO categories in this list after Benjamini-Hochberg adjustment; however, terms such as ‘embryonic skeletal system morphogenesis’ or ‘positive regulation of chondrocyte differentiation’ were ranked forward. There are 28 known chondrogenesis-related genes in this cluster, including genes encoding ADAMTSs (*ADAMTS7* and *12*), *DNMT1*, *HOXA3*, *HOXA11*, *HOXD3*, *SOX6*, and *SMAD5*. The following genes were characterised by the highest number of known connections and closeness centrality values in this cluster: epidermal growth factor receptor (*EGFR*), Erb-B2 receptor tyrosine kinases 2 and 4 (*ERBB2* and *ERBB4*), *SMAD3*, *DNMT1*, *NCAM1*, and *WNT1*.

There were 2328 genes in **cluster 3**. This group is characterised by a steady low-level expression profile from day 0 to day 6 cultures, and then starts to pick up and display peak expression in mature articular chondrocytes (Fig. 7). Significantly over-represented GO terms include generic terms such as ‘transport,’ ‘endoplasmic reticulum to Golgi vesicle-mediated transport,’ ‘lipid biosynthetic process,’ and ‘lipid metabolic process,’ but notable terms such as ‘chondroitin sulfate biosynthetic process’ and ‘extracellular matrix structural constituent’ were also on top of the list ranked according to BH-adjusted *p*-values (Fig. 8). As can be inferred from the expression pattern, this cluster had the highest proportion of genes (71 genes) with a known function in chondrogenesis according to the cartilage development (GO:0051216) list. This group of genes contained, among many others, key transcription factors (*SOX5*, *RUNX1, RUNX3*, *HIF1A*) and most of the genes encoding ECM constituents, including *ACAN*, alpha chains of various collagens (*COL1A1*, *COL2A1*, *COL3A1*, *COL4A1*, *COL6A1*, *COL8A1*, *COL9A1*, *COL10A1*, *COL15A1*, *COL20A1*, *COL28A1*), *COMP*, *MATN1* and *3*, and *TRPV4*. The following genes had the highest number of edges and closeness centrality values in this cluster: protein kinases (including *MAPK3* and *JUN*), *FN1* (fibronectin), key transcription factors (*HIF1A*), signal transducer and activator of transcription 3 (*STAT3*), and hyaluronate receptor *CD44*.

**Cluster 4** comprised 1313 genes. This cluster is characterised by a biphasic expression pattern: genes in this group initially (day 0 and day 1 cultures) show high expression values, then there is a trend of downregulation until days 10 and 15, but expression levels in mature chondrocytes are once again very high (Fig. 7). There was only one significantly over-represented GO term (protein deneddylation) after Benjamini–Hochberg adjustment in this cluster (Fig. 8), but terms such as ‘protein serine/threonine kinase activity,’ ‘cellular response to vascular endothelial growth factor stimulus,’ ‘NF-kappaB binding,’ ‘hippo signalling,’ ‘regulation of calcineurin-NFAT signaling cascade,’ and ‘Notch binding’ were also enriched. Genes with the highest numbers of edges and closeness centrality values included members of the PI3K/Akt pathway (*AKT1*, *MTOR*, *PIK3C3*, *PIK3CA*), translation initiation and elongation factors (*EIF4G1* and *EEF2*), cyclin-dependent kinase *CDC42*, and proteasome subunits (*PSMA3*, *PSMD2*, *PSMD14*, *PSMC6*). There were 27 genes in this cluster that were included in the GO:0051216 list for cartilage development. For example, this cluster contains *BMP2, BMPR1A*, *SMAD1*, and *WNT9A*, which are expressed in chondrocytes.

The 2002 genes in **cluster 5** initially showed high expression levels and were then characterised by a trend of downregulation towards the later time points, particularly in mature articular chondrocytes (Fig. 7). This pattern likely includes genes essential for regulating proliferation-related processes during the early stages of chondrogenesis. Indeed, GO terms (biological process, molecular function) related to the cell cycle, DNA replication or DNA repair such as ‘chromosome organization,’ ‘cell cycle,’ ‘mitotic sister chromatid segregation,’ ‘single-stranded DNA helicase activity,’ ‘catalytic activity, acting on DNA,’ and ‘regulation of nucleobase-containing compound metabolic process’ are significantly enriched in this cluster (Fig. 8). There were 34 genes common between this cluster and the cartilage development GO list, including *HOXA5*, *WNT5A*, *BMP4*, *BMP7*, and *BMPR2*. Genes with the highest known connections and closeness centrality values were β-actin (*ACTB*), β-catenin (*CTNNB1*), *NOTCH1*, *HDAC2*, *SIRT1*, and *FOXM1*.

**Cluster 6** contained 1726 genes. Genes in this cluster displayed a robust upregulation pattern from days 0 to 6, followed by a massive decline in mature micromass cultures and articular chondrocytes (Fig. 7). These genes likely play important roles in primary chondrogenesis but are downregulated in mature micromasses and articular chondrocytes. In line with this, the following relevant GO terms are significantly over-represented in this cluster: ‘collagen biosynthetic process’ and ‘collagen metabolic process’ (Fig. 8), but other chondrogenesis-related terms are also ranked forward such as ‘cartilage development,’ ‘cartilage condensation,’ ‘connective tissue development,’ ‘mesenchymal cell differentiation,’ ‘skeletal system development,’ and ‘limb development.’ Transcripts with the highest number of edges and closeness centrality values included *PRRC1*, a protein kinase A regulator, *NUDT13*, which is involved in NADH homeostasis, *SNCA* (synuclein alpha); the voltage-gated calcium channel subunit *CACNA1C,* and the calcium-calmodulin dependent kinase *CAMK2A*. There were 54 genes in this cluster (second only to cluster 3) relevant to cartilage formation, including various collagens (*COL5A1*, *COL5A2*, *COL8A2*, *COL12A1*, *COL13A1*, *COL14A1*, *COL16A1*, *COL22A1*, *COL23A1*, *COL27A1*), *FGFR1* and *FGFR2*, *FOXC1*, *GDF2*, *HOXD11*, *SEMA3D*, *TFGBI*, *WNT2B*, *WNT5B*, and *SOX9*.

### The Temporal Expression Pattern of Chondrogenic Genes and Collagens During Chondrogenic Differentiation

We performed a targeted *in silico* analysis of the NGS data to determine the profiles of common chondrocyte-specific markers in chondrifying cultures. To this end, we generated a list of genes with functions in chondrogenesis and chondrocytes by merging the entities of the following GO categories: GO:1990079 (cartilage homeostasis); GO:0051216 (cartilage development); GO:0060536 (cartilage morphogenesis); GO:0001502 (cartilage condensation); and GO:0061975 (articular cartilage development) (see Supplementary file 2 – ‘C*hondrogenic’ worksheet;* Fig. 9A). As expected, the “classical” hyaline chondrocyte marker genes showed different temporal patterns between time points. *COL2A1* had the highest normalised expression level on days 6 and 10 (over 300,000), followed by a steady increase along with chondrogenesis, and peaked in mature chondrocytes (see cluster 3). The same temporal expression pattern and high normalised expression values were observed for some of the other ECM constituents, such as classical collagen alpha chains (*COL1A1*, *COL6A1,* and *COL9A1*), *ACAN, COMP,* and *MATN1, 2* and *3* (all in cluster 3). Other highly abundant (>3,000) transcripts include *FGFR3*, *HAPLN1*, *NCAM1*, *NFATC1*, *3,* and *5*, *SOX9*, and *SMAD6*. In contrast, some of the “classical” cartilage markers, such as the growth differentiation factor (GDF) genes, generally showed low (<1,000) normalised expression levels. Most of the genes in this group displayed a trend of upregulation at later time points, but some other key factors (e.g., *BMP7*, *GDF11*, *NCAM1*, *PAX7*, *SMAD1,* and *WNT7A*) showed the opposite trend (Fig. 9A). Markers of chondrocyte hypertrophy (45), including *RUNX2*, *GADD45B*, *IL8*, S100 proteins (cluster 1), *NFATC1*, *SMAD3*, *SMAD5* (cluster 2), *CEBPB*, *SMAD2*, *DDR2*, *MATN3*, *TGM2* (cluster 3), *SMAD1* (cluster 4), catenin beta (*CTNNB1*), and *SIRT1* (cluster 5), showed variable expression patterns, but most of them were upregulated in day 15 micromass cultures.

**Figure 9.**
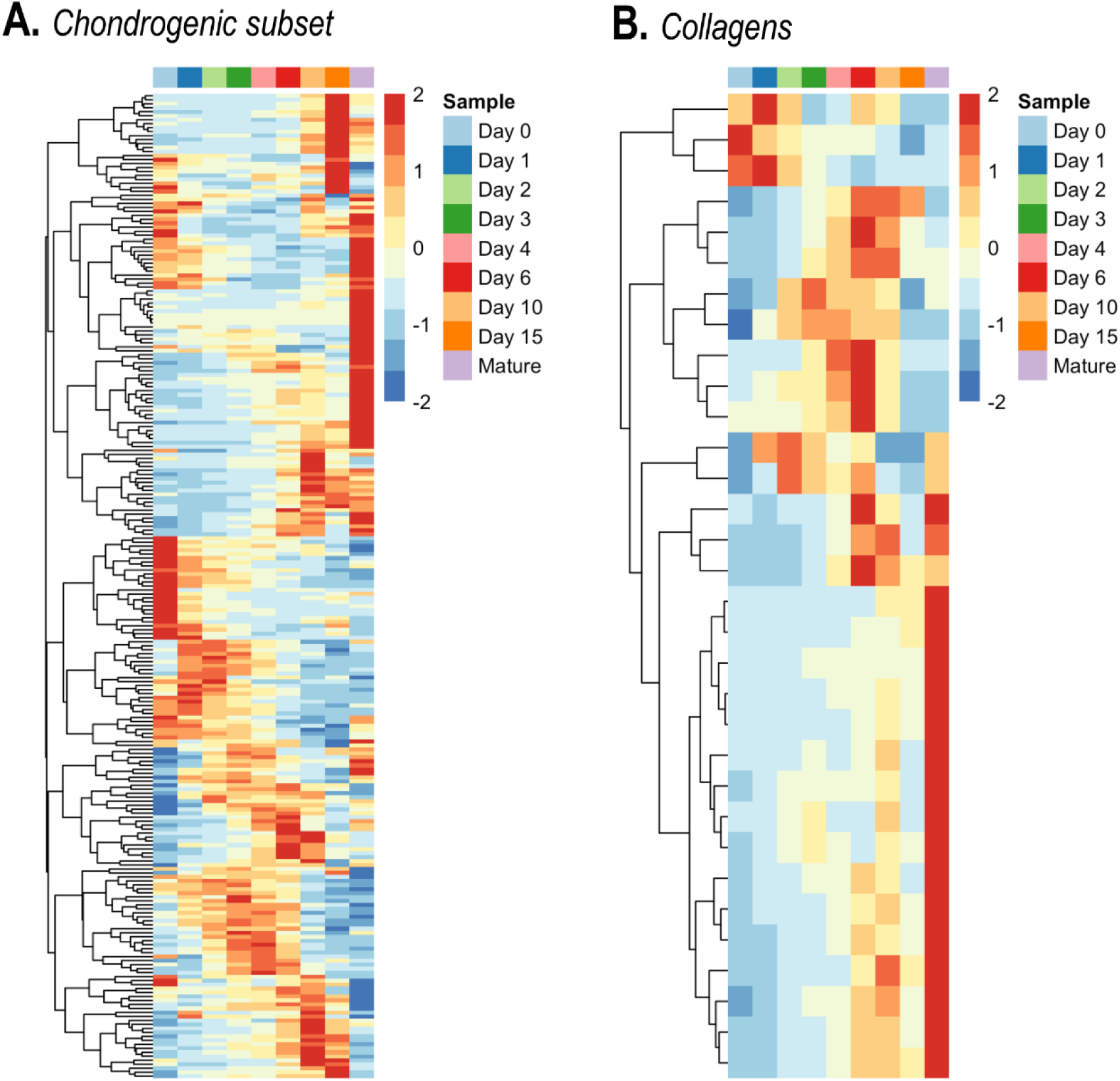
Hierarchical clustering of the (**A**) chondrogenic subset, and (**B**) collagens in differentiating cells of micromass cultures undergoing chondrogenesis during days 0-15, as well as mature articular chondrocytes, based on normalised expression values. Average values for the 3 biological replicates are shown.

Because various collagen types have been described as major or accessory components of cartilage ECM, we investigated which collagen-coding transcripts were expressed in chondrifying micromass cultures (Table S4 in Supplementary file 1; Fig. 9B). We detected genes coding for alpha chains of almost all known collagen types (I–XXVIII), except for types VII, XI, XIX, XXI, XXV, XXVI, and XXIX. While the genes coding for collagen types displayed variable expression patterns, the majority were grouped into clusters 3 and 6, indicating an upregulation trend. Some collagens, such as *COL17A1* and *COL18A1* displayed a downward trend towards the more aged micromasses, whereas others, including *COL5A1*, *COL5A2*, *COL8A2*, *COL12A1*, *COL13A1*, *COL14A1*, *COL16A1*, and *COL27A1* exhibited a peak-like expression pattern (Fig. 9B). *COL2A1* had the largest overall (∼250,000) and absolute (∼240,000 on day 10) average normalised expression values; moreover, its levels were extremely high (approximately 1,500,000) in mature chondrocytes. In addition to *COL1A1*, transcripts of “classical” cartilage-specific collagens (genes coding for collagen type 6 and 9 subunits, such as *COL6A2* and *COL9A1*) also had a high relative normalised expression. The genes coding for minor but still abundant (<10,000 relative expression value) collagen types included *COL3A1*, *COL5A1*, *COL12A1*, *COL14A1*, *COL16A1*, and *COL27A1*.

### Gene Co-Expression Networks

Using unsigned weighted gene correlation network analysis (WGCNA), we first generated a dendrogram and a trait heatmap of the samples using the three classical chondrogenic markers *SOX9*, *COL2A1,* and *ACAN*, as well as age as ‘traits’ (Fig. 10). As seen earlier (see Fig. 5), hierarchical clustering showed a separation between groups of biological replicates according to age and the three marker genes for 0-, 1-, and 2-day-old micromasses, whereas day 3 and 4 cultures, as well as day 6, 10, and 15 cultures, clustered together (Fig. 10A). We then searched for subsets of genes that were highly correlated with the expression patterns of the three marker genes and age, referred to as ‘traits’. To this end, genes were clustered into 48 modules of correlated expression patterns, with each module designated an arbitrary colour (Fig. 10B). We selected the top three modules (*sienna3*, *thistle2*, and *pink*) for detailed analysis because they showed the highest positive correlation with all four investigated traits. The three modules combined contained 622 genes (see Supporting Information, Supplementary file 2 – ‘*WGCNA’ worksheet*). GO pathway analysis performed on this list revealed that the following terms relevant to chondrogenesis and ECM production were significantly enriched: ‘collagen trimer,’ ‘extracellular matrix,’ ‘signal transduction,’ and ‘cell communication.’ Because these pathways are highly relevant to chondrogenic differentiation, these modules can be classified as functional chondrogenic modules. However, out of the 622 genes on this list, there were only 17 common entities with genes known to be involved in chondrogenesis/chondrocyte functions (GO:0051216). These included *CHADL*, *RUNX1*, *COL3A1*, *COL9A1*, *CSGALNACT1*, *MATN1*, and *TGFBR2*. We identified the following 10 transcription factors in this list: *ATOH8*, *FOXO1*, *HBP1*, *HOXA10*, *HOXA11*, *HOXB3*, *POU5F3*, *RUNX1*, *SOX3*, and *TBX2*, of which only *HOXA11*, *HOXB3*, and *RUNX1* were listed under the GO:0051216 term.

**Figure 10.**
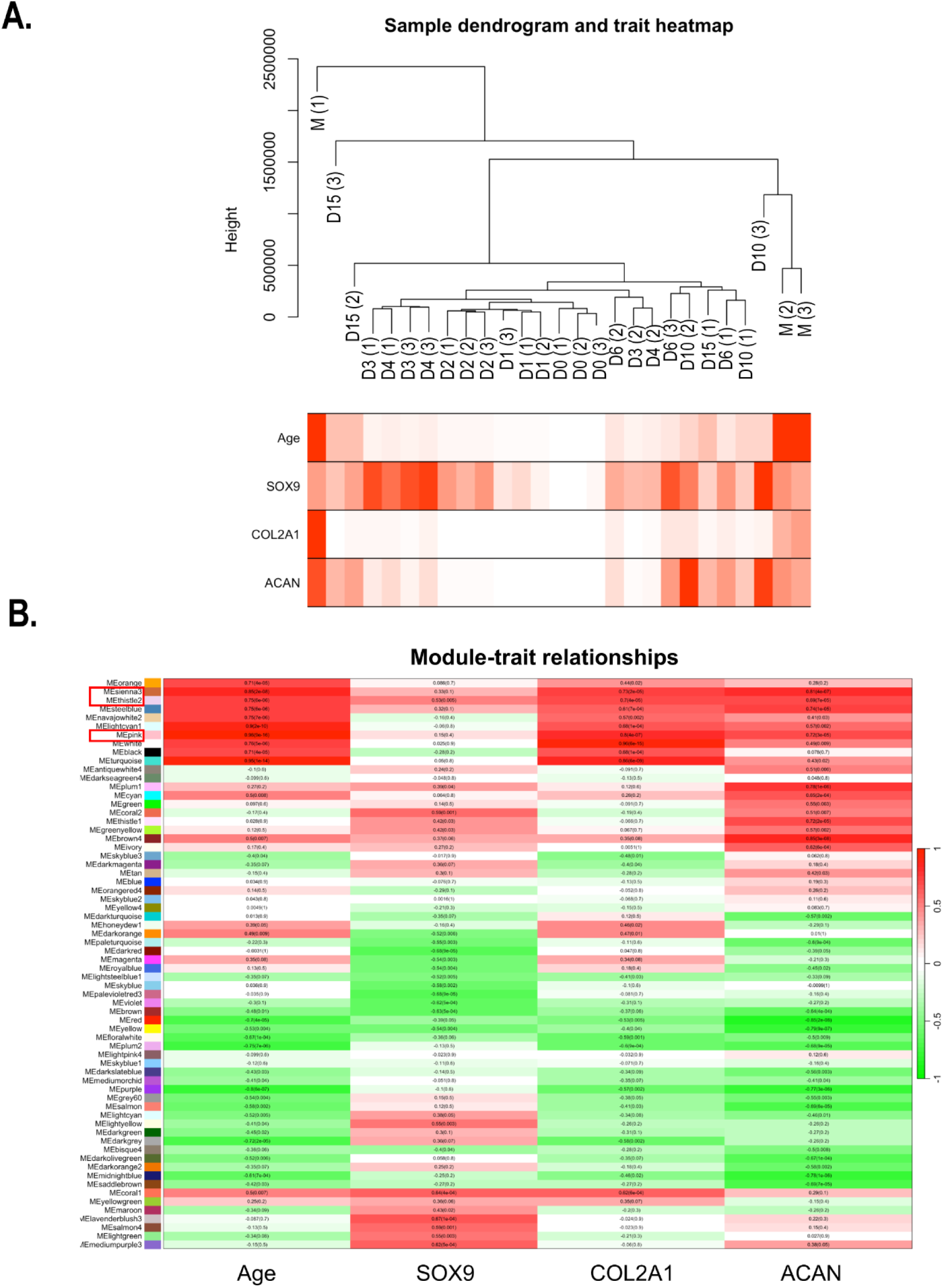
Unsigned WGCNA was used to identify subsets of genes that were highly correlated with the following traits: time points (age), *SOX9*, *COL2A1*, and *ACAN* expression patterns. Genes were then clustered into modules designated with arbitrary colours. (**A**) Dendrogram of RNA-seq samples (codes: D, day; M, mature; numbers in brackets indicate biological replicates) and corresponding changes in traits. The lowest values are shown in white; the highest values are depicted in red. (**B**) Modules whose eigengenes are highly correlated with age (days in culture) and the expression patterns of *SOX9*, *COL2A1*, and *ACAN* are highlighted by a red frame (*sienna3*, *thistle2*, *pink*); their corresponding Pearson correlation values are shown.

We then extracted the genes from the merged contents of the top three modules to Cytoscape and selected the top 50 most highly correlated genes based on closeness centrality values. To identify the key driving molecules in the network, the genes were sorted according to closeness centrality values and the connections between them were visualised (Fig. 11). *CHADL*, *IGFR2*, and *RUNX1* were among the genes with the highest numbers of connections.

**Figure 11.**
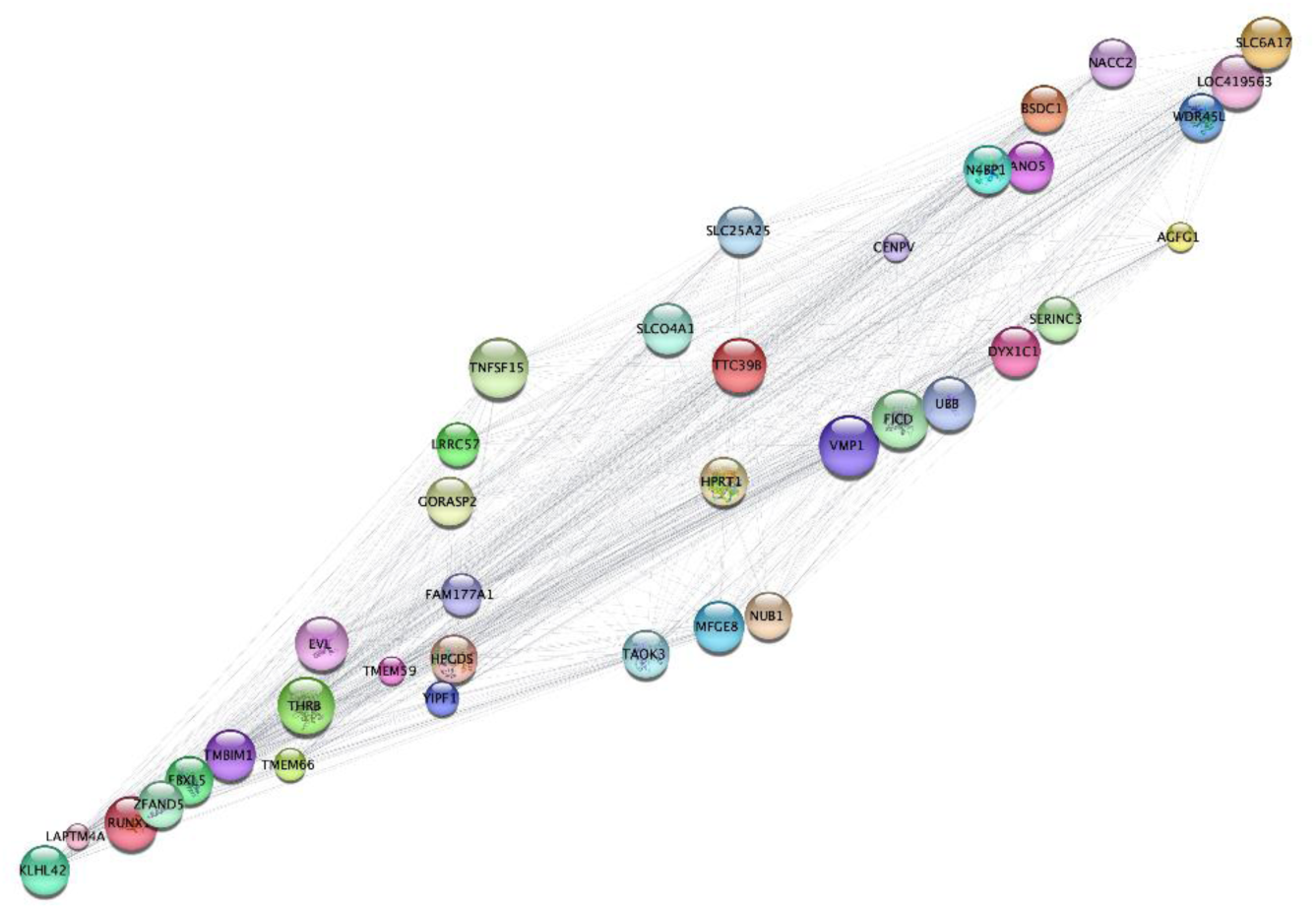
The edge data of the top ∼500 genes from the *sienna3, thistle2,* and *pink* modules from the WGCNA analysis were exported to Cytoscape. The genes were then sorted according to closeness centrality values, and the connections between the top ∼50 were visualised. Node size indicates closeness centrality values; edge length represents the strength of the correlation between the respective nodes.

### Transcription Factors in Chondrogenesis

Using the GO term GO:0003700: ‘transcription factor activity, sequence-specific DNA binding,’ we searched for TFs in our dataset. Of the 1175 genes in this GO category, 483 transcripts had detectable expression levels following normalisation of our dataset (Fig. 12A; see list in Supporting Information, Supplementary file 2 – ‘*TFs-in-dataset’ worksheet*). We then excluded transcripts that did not meet the following criteria from further analysis: LFC > 1.8 or < –1.8; geometric mean normalised expression values > 500 at least at the time point immediately after upregulation/before downregulation. We matched the entries on this list, containing 170 differentially expressed TFs with the entities of the GO term ‘cartilage development’ (GO:0051216). Only 27 entries (∼16%) of our list of TFs were previously annotated under the GO:0051216 term (Supplementary file 2 – ‘*TFs-in-dataset’ worksheet*). With the strictness of our selection criteria in mind, we advocate that the remaining 84% of TFs in our list should be considered for further exploration, and possibly as an addition to the list of genes involved in cartilage development as either positive or negative regulators. It should also be noted that 63 (∼37%) of the 170 genes (Supplementary file 2 – ‘*TFs-in-dataset’ worksheet*) were also differentially expressed during the chondrogenic differentiation of human MSC cultures (hMSC data were retrieved from Gene Expression Omnibus, <u>GSE109503</u>), and only 11 of these (*HIF1A, NFIB, NKX3-2, PRRX1, RARB, SCX, SMAD3, SNAI1, SOX5, SOX9 and TRPS1*) were previously annotated under the GO term cartilage development (GO:0051216).

**Figure 12.**
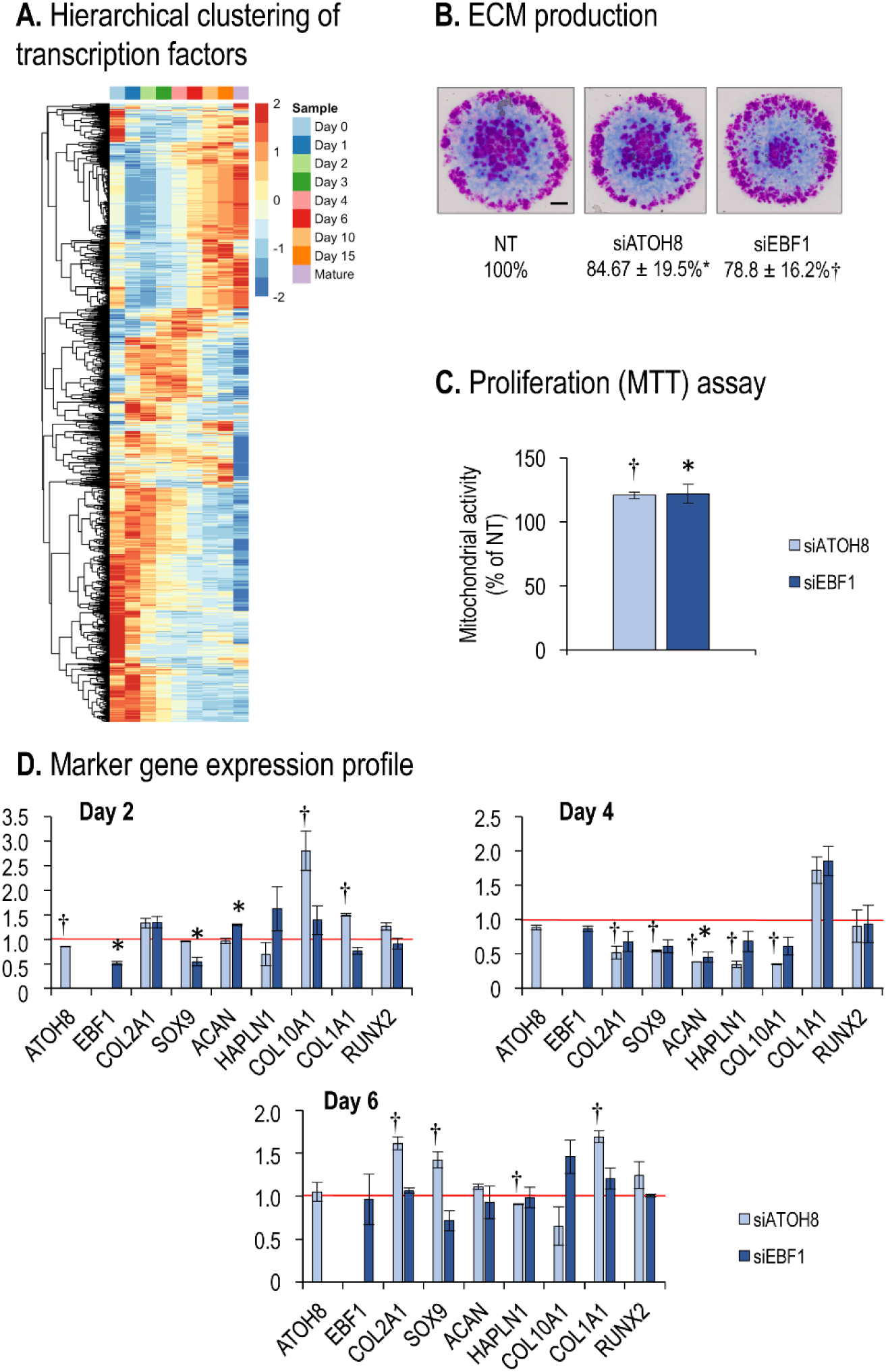
Transcription factors in chondrogenesis of micromass cultures. (**A**) Hierarchical clustering of the transcription factors in differentiating cells of micromass cultures undergoing chondrogenesis during days 0-15, as well as mature articular chondrocytes, based on normalised expression values. Average values for the 3 biological replicates are shown. (**B**) Metachromatic cartilage areas after DMMB staining of 6-day-old micromass cultures following electroporation with *ATOH8* or *EBF1* siRNA, or non-targeting (NT) control. Original magnification was ×4. Scale bar, 1 mm. Values are results of a MATLAB-based image analysis of metachromatic areas. Data are expressed as mean ± SEM, compared to NT (100%). Representative data of 3 biological replicates. (**C**) MTT assay results carried out 6 days post electroporation with siATOH8 or siEBF1, normalised to the NT control. Average data of 3 biological replicates. (**D**) Gene expression data of selected osteo/chondrogenic markers 2, 4, and 6 days after transient gene silencing. Relative gene expression values are calculated by normalising to *RPS7*. Data are expressed as mean ± SEM (n=3). For panels **C–E**, significant differences (*p* < 0.05) in relative gene expression data relative to NT controls are indicated by a dagger sign (†, in case of *ATOH8*) or an asterisk (*, in case of *EBF1*).

When looking separately at our selection criteria for TFs (see lists in Supporting Information, Supplementary file 2 – ‘*TFs-in-dataset’ worksheet*), 255 TF genes have an mRNA expression that reaches a normalised relative expression of > 500 at least at one time point of sampling; 35 (∼14%) of these entries are annotated under GO:0051216 (Supplementary file 2 – ‘*TFs-GO0015216-1’ worksheet*). Some TFs are steadily expressed during the entire cartilage differentiation period and are necessary for the non-stage-dependent processes of chondrogenesis and cartilage homeostasis. We identified 68 entries in the chicken model that had a persistent normalised expression value of > 500 at all time points for sampling (including mature articular cartilage). Ten of these (∼15%) were also annotated under the ‘cartilage development’ GO term (GO:0051216; Supplementary file 2 – ‘*TFs-GO0015216-2’ worksheet*). If our full list of TFs expressed at detectable levels at least at one time point (with 483 entries) was aligned with the list for cartilage development (GO:0051216), 49 entries (∼10%) emerged as matches (Supplementary file 2 – ‘*GO3700*-*GO15216-2’ worksheet*). These results signify a relative enrichment of genes with chondrogenic associations among TFs with a higher normalised read count (∼15% vs. ∼10%). In hMSC cultures undergoing chondrogenic differentiation, a very similar proportion of differentially expressed TFs (under the same criteria defined above) was annotated under the GO:0051216 term (22 entries from a total of 205, ∼11%; see Supplementary file 2 – *‘GSE109503-GO’ worksheet*), the total number of entries that fit our criteria was also quite similar (205 *vs*. 170 in chicken).

Of the 63 genes that were differentially expressed in both chicken micromass cultures and chondrogenic hMSC cultures (see Supplementary file 2 – *‘Chicken-vs-GSE109503’ worksheet*) and were not annotated under the GO:0051216 term, only 11 genes were regulated in the same direction and at comparable time stages of chondrogenesis (*BARX1*, *BHLHE41*, *CEBPD*, *ETS2*, *FOS*, *FOXA2*, *HES1*, *HLX*, *NR4A2*, *PITX2* and *SOX8*). However, *BARX1* and *NR4A2* only exceeded our cut-off value (normalised expression value of > 500) in mature articular cartilage samples, somewhat diminishing their potential role in chondrogenic development.

With a common set of DEGs that have relatively understudied roles in chondrogenesis, we aimed to place emphasis on those that are more perplexing and have a dissimilar/converse expression pattern in hMSCs and chicken LMPs (see Supplementary file 2 – *‘Chicken-vs-GSE109503’ worksheet*). This is of particular relevance because hMSCs have a stronger tendency to produce fibrocartilage-like tissue instead of hyaline cartilage *in vivo* according to clinical trial reports (46, 47). Due to the technical circumstances of siRNA transfection in the micromass model, it is ideal to target TFs that are upregulated early (with FC > 1.8) in the chicken model, while downregulated early (with FC < –1.8) in hMSC cultures. Six genes showed expression patterns that matched the above criteria: *ATOH8*, *EBF1*, *NFIA*, *NFIX*, *PLSCR1* and *STAT1*. Of the six genes listed above, *ATOH8* and *EBF1* were selected for further experiments because they were also identified by other analyses (see above).

We performed *in silico* signalling network analysis for all six genes using the SIGNOR 3.0 public repository (48), mainly looking for curated interactions where they are present as regulators not restricted to any species. Data are available for *NFIA*, *NFIX* and *STAT1*. For the other genes, for which sufficient SIGNOR data were not available (*ATOH8*, *EBF1* and *PLSCR1*), STRING interaction networks were analysed. Following the extraction of the top 25 interaction partners for each gene (retrieved from *Homo sapiens* data for best data abundance), their expression levels were analysed and matched to the original gene of interest. In all cases, only interacting partners with normalised expression levels of > 500 at least at one time point were considered for the analysis. All fold change values at individual time points were compared to that of the gene of interest, and partners with the lowest average difference in FC ratios were considered the best matches. The three interacting partners with the most similar expression patterns for each of the six genes above (*TWIST2*, *SMARCB1*, and *ARID1A* for *ATOH8*; *ZNF423*, *CREBBP*, and *ZNF521* for *EBF1*; *NFIX*, *WNT5A*, and *MYOD1* for *NFIA*; *WNT5A*, *ROBO1*, and *ETV5* for *NFIX*; *SHC1*, *LZTR1*, and *PRKCD* for *PLSCR1*; and *RELA*, *IRF2*, and *S100A10* for *STAT1*) are shown in Supplementary file 1, Fig. S2.

### Transient gene silencing of *ATOH8* or *EBF1* reduces chondrogenesis in micromass cultures

Next, we transiently silenced the expression of *EBF1* or *ATOH8* in chondrogenic micromass cultures by electroporation prior to inoculation as micromass cultures (day 0). In addition to the criteria described above, *EBF1* was chosen because it is an upstream regulator of SOX9 expression (Table S5 in Supplementary File 1). *ATOH8* was a particularly interesting candidate, as it was also identified in our WGCNA analysis (Supplementary File 2 – *‘WGCNA’ worksheet*). Transient gene silencing of *ATOH8* or *EBF1* on day 0 resulted in a significant (84.67 ± 19.5%, or 78.8 ± 16.2% of non-targeting (NT) control, respectively) reduction in cartilage ECM deposition by day 6 (Fig. 12B). At the same time, the rate of cell proliferation was significantly increased (120.8 ± 2.4%, or 122.1 ± 7.3% of the NT control, respectively) following transfection with siRNA by day 6, as inferred from the results of MTT assays (Fig. 12C). We then analysed the expression levels of key osteochondrogenic marker genes 2, 4, and 6 days post-transfection using RT-qPCR (Fig. 12D). In 2-day-old cultures, gene silencing resulted in approx. 15% and 50% downregulation of *ATOH8* and *EBF1*, respectively. This corresponds well with the higher expression levels of *ATOH8* in micromass cultures, because target gene abundance influences the efficiency of siRNA-mediated gene silencing (49). siEBF1 significantly downregulated *SOX9* expression (to 55% of the NT control), whereas siATOH8 significantly upregulated *COL10A1* and *COL1A1*. By day 4, although the silencing has diminished in case of both *ATOH8* and *EBF1* expression levels, significant downregulation of the chondrogenic markers *COL2A1*, *SOX9*, *ACAN*, and *HAPLN1*, as well as *COL10A1* was observed in siATOH8 cultures, whereas siEBF1 downregulated *ACAN*. This is likely attributable to the perturbation of the complex downstream chondrogenic signalling cascade and indicate that even slight changes in the expression of transcription factors can have a significant impact. In 6-day-old cultures, when *ATOH8* levels are normally very low, transient gene silencing resulted in a compensatory upregulation of *COL2A1*, *SOX9*, and *COL1A1*, and downregulation of *HAPLN1* expression. These data indicate that the two transcription factors are indeed involved in the regulatory network of chondrogenesis, and that they differentially regulate the expression of osteochondrogenic marker genes in micromass cultures.

## DISCUSSION

In this study, we examined gene expression patterns during cartilage formation in micromass cultures of embryonic limb bud-derived progenitor cells using RNA sequencing, and used the transcriptome of chondrocytes isolated from chicken articular cartilage as a control. We detected a progressively different and distinct transcriptome during key stages of chondrogenesis. We confirmed the involvement of the top DEGs in chondrogenic differentiation by pathway analysis. We identified several chondrogenesis-associated transcription factors (TFs) and collagen subtypes that were not previously linked to cartilage formation. Specifically, we confirmed that *ATOH8* and *EBF1* are transcription factors involved in chondrogenesis because transient gene silencing attenuates chondrogenesis and deregulates the expression of osteochondrogenic marker genes.

*In situ* hyaline cartilage formation in the embryo is a multistep, dynamic, and sequential process controlled by several TFs, soluble mediators, ECM, cell-cell, and cell-matrix interactions, as well as epigenetic and miRNA-mediated mechanisms (50–52). In stark contrast, the standard *in vitro* culture conditions for the differentiation of MSCs into cartilage are simple and have remained largely unchanged since the original description (53). Current cartilage regeneration options, even the “gold standard” method using a combination of growth factors, scaffolds, and MSCs, do not reliably produce hyaline cartilage in favour of biomechanically inferior fibrous cartilage (54). Therefore, there is a critical need for more effective hyaline cartilage regeneration therapies, for which a thorough understanding of the molecular events underlying chondrogenic differentiation is essential. To the best of our knowledge, this is the first study to examine the global temporal transcriptome landscape of *in vitro* chondrogenesis by high-throughput RNA sequencing using a well-established embryonic limb bud-derived 3D micromass system that closely recapitulates the process *in vivo*.

High-throughput transcriptomic technologies, such as microarrays and RNA sequencing are tools for studying the expression patterns of thousands of transcripts, allowing the identification of differentially expressed genes between two or more conditions and describing altered biological pathways. Studies on unbiased global transcriptomic profiling of the key regulators involved in chondrogenesis are scarce, and those that are available are performed using different model systems. Microarrays have previously been used to examine temporal gene expression patterns during *in vitro* chondrogenic differentiation in a mouse micromass culture systems (55). Global gene expression profiling analysis of microdissected chondrogenic tissues derived from tibial and fibular pre-condensed mesenchyme of mouse hind limbs using microarrays was the first *in vivo* transcriptomic study of cartilage development (56). Temporal changes in key molecular components during the formation of hyaline-like cartilage from infrapatellar fat pad-derived MSCs in micromass cultures have been studied using microarray gene expression analysis (57). Microarrays have also been used to analyse and compare global gene expression (58) and miRNA expression profiles of chondrocytes derived from human induced pluripotent stem cells (hiPSCs) (59). However, microarrays have limited ability to quantify the expression levels of a set of predetermined transcripts printed on a given chip. Therefore, recent studies have employed ultrahigh-throughput RNA sequencing (RNA-seq), which has several advantages over conventional microarrays.

Detailed unbiased NGS data on chondrogenic differentiation are scarce. Recently, a dataset of high-throughput RNA sequencing of pellet cultures generated from primary human bone marrow-derived MSCs induced towards the chondrogenic lineage at six different time points (day 0 to day 21) was published (6). The authors identified a chondrogenic gene subset containing 1172 entities, whose functional characterisation promises to harness the potential of MSCs for cartilage tissue engineering. The same laboratory recently published a paper on chondrogenic differentiation of hiPSCs using bulk and single-cell RNA sequencing, and identified gene regulatory networks regulating chondrogenesis (60). Temporal changes in the transcriptome of human embryonic stem cells (ESCs) during key stages of chondrogenic differentiation have also recently been reported (61).

All the above studies were conducted on cells that do not spontaneously generate cartilage *in vitro*, and existing protocols rely on the addition of growth factors and other compounds, such as insulin-transferrin-selenium, dexamethasone, ascorbic acid, L-proline, TGF-β3, GDF5, FGF2, and NT4 to achieve chondrogenesis *in vitro*. The main advantage of the embryonic limb bud progenitor-derived micromass model is that, unlike most other differentiation techniques, this protocol does not require additional growth factors and other reagents to achieve high-quality chondrogenic differentiation and ECM similar to hyaline cartilage.

The chicken limb bud-derived micromass model is conventionally used to study the early stages of chondrogenic differentiation until day six of culture (16, 19). A low-dimensional space representation of the RNA-seq data, including PCA and UMAP, was used to reconstruct the differentiation trajectory of the chondrogenic model. In the PCA plots, samples from early chondrogenic cultures (between days 0 and 6) displayed a gradual shift along the *x*-axis as chondroprogenitor cells differentiated into chondroblasts and chondrocytes, with mature articular chondrocytes positioned at the far end. However, the day 15 samples seem to have lost this trend, suggesting that these cultures are positioned “backward,” closer to the immature cultures. Therefore, we aimed to confirm this apparent differentiation trajectory by using a dedicated method. Following UMAP dimension reduction using the Monocle 3 R package, RNA-seq data were ordered with reverse graph embedding, and pseudotime was calculated for each culture of known latent development time. When pseudotime was plotted against UMAP component 1, there was an obvious trend for cultures to progress systematically through pseudotime at the early time points (days 0–4), but there was little separation once they reached 4.5, where more mature micromass cultures were clustered. When plotting numerical time *vs*. pseudoage, this “developmental saturation,” occurring from approximately day six, was even more obvious. These data indicate that the gene network undergoes significant rearrangement mainly until day 6, whereas more aged micromasses are characterised by less profound changes at the global transcriptomic level.

### Chondrogenic Gene Expression Profile

Most studies on MSC-derived cartilage regeneration have relied on the analysis of a selected subset of common cartilage marker genes. To this end, we prepared a custom-made list of “classical” markers for hyaline cartilage by merging the genes annotated with relevant GO terms. We found that all genes on these lists were expressed in the micromass model but with different temporal dynamics (i.e., were assigned to all six unsupervised clusters) and expression levels. For example, SOX9, a pivotal TF in chondrogenesis and mature cartilage, along with its partners SOX5 and SOX6, followed a typical “chondrogenic” pattern, displaying gradually higher transcript levels in more mature cultures (assigned to clusters 2 and 6). In addition to securing and maintaining the commitment of skeletogenic progenitor cells to the chondrocyte lineage, SOX9 is critical to ensure adult articular cartilage maintenance and function by transcriptionally activating genes (i.e., *COL2A1* and *ACAN*) essential for cartilage ECM and repressing factors and pathways that favour non-chondrocyte lineages (62). In contrast to an earlier microarray-based study, which only managed to detect *SOX4*, *SOX5*, *SOX8*, *SOX9,* and *SOX11* in normal and OA cartilage (63), we identified transcripts for 12 SOX transcription factors, with *SOX9*, *SOX6*, *SOX5*, *SOX11*, *SOX8,* and *SOX4* displaying the highest read numbers, which confirms the much higher sensitivity of the RNA-seq-based methodology. Of these, only the SOX-trio (*SOX9*, *SOX5,* and *SOX6*) followed the typical “chondrogenic” pattern. We also checked the profiles of TFs known to control *SOX9* expression (Table S5 in Supplementary file 1). Among these TFs, *HIF1A*, *JUND*, *ATF3*, *CHD2,* and *HDAC2* were most abundant. Moreover, *EBF1*, *FOS*, and *MYC* were identified as the TFs of key importance in our detailed analysis.

The most abundantly expressed transcripts in chondrogenic cultures included ribosomal proteins (indicating preparation for intense protein synthesis), chaperones, cytoskeletal components (members of the actin cytoskeleton and the microtubular system), ECM components such as collagen (*COL1A1*, *COL2A1*, *COL9A1*), fibronectin (*FN1*), aggrecan (*ACAN*), multifunctional heterogeneous nuclear ribonucleoproteins (hnRNPs) mediating mRNA metabolism, transcription factors, initiation and elongation factors (*EEF1A1* and *EIF5B*), and glycolytic enzymes (*GAPDH* and *ENO1*). As chondrocytes are highly glycolytic cells, the fact that these genes were found in high abundance corroborates the literature (64). The above functional classification of the highly abundant transcripts was also confirmed by GO pathway analyses. These results are in perfect agreement with previously published data (14, 65).

When we examined the DEGs by pairwise comparisons between time points, we identified several genes with important roles in skeletal development. In the early stages of chondrogenic differentiation (transition from chondroprogenitors to ECM-producing chondroblasts), *GDF5*, osteocrin (*OSTN*), biglycan (*BGN*), and cellular communication network factor-3 (*CCN3*), which are key components of chondrogenesis, were among the top DEGs (14). We also noted robust differential expression of transcripts involved in ECM production or remodelling, such as chondrolectin (*CHODL*), cartilage-derived C-type lectin (*CLEC3A*), matrix metalloproteinases (*MMP1*, *MMP9, MMP10*, *MMP11, MMP13*), *ADAMTS3*, collagens (*COL10A1*, *COL14A1*), matrilin-1 (*MATN1*), and chondroadherin (*CHAD*), which is also in line with the literature (66).

To further characterise the genes directly involved in chondrogenic differentiation, we employed gene co-expression analysis using WGCNA, based on the assumption that co-expressed genes are functionally related and co-regulated (6). The rationale behind choosing the three classical chondrogenic markers *SOX9*, *COL2A1,* and *ACAN,* as well as age (maturity), as traits in our WGCNA analysis, was that the genes showing a highly correlated expression pattern would likely be involved in governing chondrogenic differentiation and ECM production in this model. The ‘chondrogenic functional modules’ (*sienna3*, *thistle2*, *pink*) contained 622 genes with a very high correlation to the above traits. Genes in this module included voltage-gated calcium and potassium channel subunits and other transporters, cadherins, integrins, carbohydrate sulfotransferases, collagen alpha 1 chains, growth factors, TFs with known involvement in chondrogenesis, and lncRNAs and miRNAs. Specifically, we selected TFs, including *ATOH8*, *FOXO1*, *HOXA10*, *HOXA11*, *HOXB3*, *POU5F3, RUNX1*, *SOX3*, and *TBX2*, and signalling molecules, such as FGFs and FGFRs, semaphorins (*SEMA7A*), *TGFBR2*, and WNTs (*WNT8B* and *WNT11B*). These results are in perfect agreement with the published data (66, 67). Reactome pathway enrichment analysis revealed that the following pathways were over-represented: ‘collagen chain trimerization,’ ‘extracellular matrix organization,’ ‘assembly of collagen fibrils and other multimeric structures,’ and ‘integrin cell surface interactions’ (Fig. S3A in the Supplementary file 1).

When we compared the genes between our chondrogenic module and those obtained in the hMSC-based chondrogenic model using a similar methodology (6), 42 common entities were identified (Supplementary file 2 – ‘*WGCNA’ worksheet*). These genes represent the ‘core’ chondrogenic subset between the two chondrogenic models, which include characteristic markers for cartilage ECM (*CHADL*, *COL3A1*, *COL9A2*), transporters and channels (*SLC1A2, SLC37A3*), the *NR1H3* nuclear receptor, and cell surface receptors (*IGF2R*). Enriched Reactome pathways with the involvement of common entities include ‘NCAM1 interactions,’ ‘nuclear receptor transcription pathway,’ and ‘collagen biosynthesis and modifying enzymes’ (Fig. S3B, Supplementary file 1). The majority of the common genes have not yet been implicated to play a role in chondrogenesis, and they therefore set the ground for more targeted studies to understand what justifies their presence in the list of core chondrogenic genes.

### Collagen Expression Profile

As SOX9 is a key regulator of genes coding for ECM components during chondrogenic differentiation, we examined the gene expression profiles of collagens. The importance of collagen in cartilage ECM cannot be overstated (68). The biological properties of cartilage primarily rely on its unique and extensively cross-linked collagen network, which exhibits regional differences in articular cartilage (69). Given the complex ultrastructure created during cartilage development, there appear to have little capacity for chondrocytes or induced MSCs to recapitulate the original collagen architecture following injury (2). For this reason, detailed knowledge of collagen subtypes, including minor collagens, expressed by *in vitro* models of chondrogenesis, is critical.

Numerous collagen subtypes have been identified in articular cartilage, including types II, III, IX, X, XI, VI, XII, XIII, and XIV (2). Collagen fibrils in articular cartilage mostly consist of type II collagen, accompanied by a lower number of minor collagens, which provide cartilage with tensile strength and contribute to the physical properties of the matrix. In our model, we identified alpha-chain transcripts for most of the known collagen types (I–XXVIII). Initially, *COL1A1* and *COL1A2* had the highest transcript numbers. However, as early as day 1, *COL2A1* took over and maintained steadily increasing expression dynamics throughout chondrogenesis, reaching over 200,000 reads by day 6. Although *COL2A1* levels declined again in more mature (10- and 15-day-old) micromass cultures, it was high expressed in mature articular chondrocytes. In contrast to our results, when bone marrow-derived MSCs were induced toward the chondrogenic lineage, the gene coding for the alpha2 chain of collagen type I (*COL1A2*) had the highest expression levels throughout, and *COL2A1* was the second most abundant collagen transcript by day 21 (mature chondrocyte stage) (6). Additionally, genes coding for collagen types XVII, XIX, XX, and XXIII were not expressed in MSC-derived chondrocytes. The genes encoding the alpha chain of type III collagen were the third most abundant collagen gene type by day 21 in that model, and the substantially high levels of collagen type X (4^th^ highest level) indicated the formation of hypertrophic chondrocytes. In contrast, type X collagen remained downregulated by day 6 in the limb bud-derived micromass model, and it only started to pick up in 10-day-old micromass cultures, but was present at low expression levels.

Looking at the collagen expression profile of human iPSCs during chondrogenesis, *COL1A1* had the highest levels by day 60, followed by *COL1A2* and *COL3A1*, with relatively low *COL2A1* levels (60). Interestingly, iPSCs did not express the genes encoding collagen types XVII and XX. This dataset also included gene expression profiles for adult, juvenile, embryonic, and 17-week human articular cartilage, which showed variable collagen gene expression profiles. One of the adult cartilage samples expressed *COL12A1* most strongly and *COL3A1* levels were also quite high. The adolescent cartilage samples showed very high *COL6A1* expression, whereas *COL2A1* was only moderately expressed. A high number of *COL2A1* transcripts are present in embryonic and week-17 cartilage samples (60).

In addition to the major collagens, several minor collagens were detected at high (>1000) read numbers in our chondrogenic model. Collagen type IV is network-forming collagen, which is known to be limited to the pericellular matrix of articular cartilage, where it may be involved in maintaining chondrocyte phenotype and viability (69). Its expression peaked in differentiating chondrocytes (day 3) and on day 10 in micromass cultures, with much higher transcript levels in articular chondrocytes. The alpha chain of collagen type V, a dominant regulator of collagen fibrillogenesis, has been documented to accumulates as articular cartilage matures (70). This is exactly what we observed in our model: *COL5A1* levels peaked in more mature cultures (days 6, 10, and 15). Furthermore, α1(V) chains were described to be cross-linked to α1(XI) chains in cartilage ECM to form V/XI polymers (70). Collagen type I can form fibrils by itself, but it can also form heterofibrils with collagen types III and V (68). Little is known about the role of non-fibrillar, short-chain collagen type VIII in cartilage, other than its presence in cartilage ECM (71). We detected the highest *COL8A1* expression levels in day 10 cultures. Collagen XIV, a FACIT collagen, is a member of the cross-linked collagen network of cartilage ECM and is predominantly expressed in late embryonic and differentiated cartilage (68, 69). Type XVIII collagen displays unique expression dynamics in that it has peak expression in immature (day 0) micromasses and then steadily declines thereafter. It is a non-fibrous collagen that has mainly been implicated in the epithelium and basement membrane in eye tissues (72) and inhibits angiogenesis (73). Thus, collagen type XVIII may play a role in the maintenance of the avascular nature of cartilage. Type XXVII collagen is prominently located at the cartilage–bone interface and surrounding proliferative chondrocytes in the epiphyseal growth plate. It plays a role in the transition of cartilage to bone during skeletogenesis (69). This was reflected by its upregulation in the 10-day-old micromass cultures.

Overall, the chondrogenic model derived from primary limb buds produced a more hyaline cartilage-like ECM than hMSCs did, at least at the transcriptional level. Although the specific role of minor collagens in chondrogenesis has not yet been explored, targeted studies may contribute to a deeper understanding of the dynamics of cartilage matrix production, which could facilitate the development of new biomarkers for joint health and drug development in OA.

### Transcription Factor Expression Profile

The human genome encodes more than 3000 TFs, several which are known to control the chondrocyte phenotype at the genomic level (67). To identify TFs that were associated with chondrogenic differentiation, but their specific roles are incompletely understood, we narrowed down the TFs identified in chondrogenic cells (GO:0003700) based on a set of criteria (fold-change > 1.8 (–1.8 in case of downregulation); geometric mean-normalised expression values > 500 at the time point following upregulation or immediately before downregulation). We then matched the chicken data with the chondrogenic human MSC transcriptome (retrieved from <u>GSE109503</u>). Of the 52 TF genes that were differentially expressed in both models and were not annotated under the GO:0051216 ‘cartilage development’ term, only 6 genes met our selection criteria. *ATOH8* and *EBF1* were chosen for transient gene silencing, as discussed below. *NFIA* and *NFIX* belong to the NF1 class of transcription factors. Although they are hypothesized to be important in the formation of the skeletal system, their specific roles in chondrogenesis remain elusive (67). NFIA is one of the genes potentially regulated by miRNAs during bone marrow-derived MSC chondrogenesis (74). *STAT1* is a member of the family of signal transducers and activators of transcription (STATs), and although it plays prominent roles in chondrocyte differentiation, we are still far from understanding its specific functions in chondrogenesis (67). STAT1 was recently identified as a molecular target of IGFBP-3 in chondrogenesis (75). Phospholipid scramblase (PLSCR) has not been implicated directly in chondrogenesis; however, it interacts with extracellular matrix protein 1 (ECM1), which is involved in cartilage formation (76).

While some of the TFs identified in this study are known to interact with *ATOH8, EBF1, NFIA, NFIX, PLSCR1* or *STAT1* have well-established roles in chondrocytes, others have not yet been described in the context of chondrogenesis. These positively correlated genes without known roles in cartilage formation include *SMARCB1* and *ARID1A* for *ATOH8*, *ROBO1* and *ETV5* for *NFIX, SHC1* and *LZTR1* for *PLSCR1*, and *RELA* for *STAT1*. Given that these entities are among the top interacting partners of key chondrogenic TFs, elucidation of their specific role(s) in hyaline cartilage differentiation warrants further studies.

### Transient Gene Silencing of *ATOH8* or *EBF1* Blocks Chondrogenesis

Since atonal homolog 8 (*ATOH8*) and early B-cell factor 1 (*EBF1*) have not yet been reported in the context of primary chondrogenesis in detail, we studied the effects of their transient knockdown at the beginning of chondrogenic differentiation. *ATOH8*, a member of the basic helix-loop-helix (bHLH) transcriptional regulators (77), was also identified in our WGCNA analysis because its expression dynamics followed that of *ACAN*, *COL2A1*, and *SOX9*. *EBF1* belongs to the EBF family of helix loop helix transcription factors (78). It is controlled by Shh signalling and is an upstream regulator of SOX9 expression. Transient gene silencing of *ATOH8* or *EBF1* significantly attenuated primary chondrogenic differentiation in micromass cultures. *EBF1* downregulated SOX9 expression 2 days post-electroporation with siRNA, and downregulated *ACAN* in 4-day-old micromasses. Simultaneously, upregulation of several chondrogenic (*COL2A1* and *SOX9*) and osteogenic (*COL1A1*) markers was observed in 6-day-old micromass cultures. These data indicated that both *ATOH8* and *EBF1* are upstream regulators of chondrogenic differentiation.

Chondrocyte-specific knockdown of *ATOH8* has been reported to reduce the size of the skeleton of axial and appendicular bones in mouse embryos *via* blocked proliferation (79). In this model, the effects of *ATOH8* gene silencing were slightly different. It significantly downregulated chondrogenic marker genes in chondroblasts, which is probably responsible for attenuated matrix production by day 6. The upregulation of *SOX9*, *COL1A1* and *COL2A1* at later time points is probably a result of a compensatory mechanism, as the effects of the siRNA diminished. In addition to bone formation, ATOH8 also regulates myogenesis and skeletal muscle maintenance (77), neurogenesis, kidney and pancreas development, and placental formation (80). ATOH8 is a direct target of the Smad-dependent TGF-ß/BMP signaling pathway, and the Alk1/Smad/ATOH8 axis regulates hypoxic response by binding to hypoxia-inducible factor 2α (HIF-2α) (80), all of which are key pathways involved in chondrogenic differentiation. Judging from the prominent shift in the expression of osteogenic and chondrogenic marker genes following gene silencing, ATOH8 likely plays a key role in determining the pathway of terminal differentiation. While there is evidence for this role (79), there are still unexplored areas to be exploited in the context of stem cell-based cartilage regenerative medicine, for which ATOH8 is a promising candidate, according to our results.

While primarily known to operate as a transcription factor to activate B cell-specific genes (81), *EBF1* is expressed in the medial region of the sclerotome and around the vertebral cartilage anlagen (78). EBF1 is co-expressed with EBF2 and 3 in the mesenchyme of developing limb buds of chicken embryos and in the perichondrium (82). Following the introduction of the dominant-negative form of EBF1 into the developing wing, chondrocyte maturation and hypertrophy are affected, resulting in shorter, thicker, and partially fused bones. Simultaneously, *SOX9*, *COL2A1* and *AP* are significantly downregulated (82). Consistent with these observations, *EBF1* gene silencing significantly downregulated *SOX9* and *ACAN* expression in our model (although at different time points). Therefore, gene silencing of *ATOH8* and *EBF1* has confirmed their involvement in regulating chondrogenic differentiation.

### Conclusions

Here, for the first time, we provide an in-depth quantitative transcriptomic landscape of *in vitro* chondrogenesis in avian embryonic limb bud-derived micromass cultures using RNA sequencing. We identified several subsets of genes, including novel TFs, associated with chondrogenesis. In addition to revealing transcriptomic signatures that are already known to control chondrogenic differentiation in other models, we provide a refined, integrated view of inter-regulatory networks that operate in addition to conventional early stage chondrogenic pathways. The data presented could be used to advance the knowledge of chondrogenic differentiation and cartilage pathology and contribute to better cartilage regeneration techniques. Transcriptomic data obtained from the model, following *in silico* analyses, may be suitable for identifying candidate genes that drive chondrogenesis to the permanent hyaline cartilage phenotype characteristic of articular cartilage rather than the transient hypertrophic phenotype.

#### Strengths of the Study

The study of human cartilage development is difficult for ethical reasons. Therefore, we used a chicken embryonic limb bud-derived micromass system to model chondrogenesis *in vitro*, which is considered to better recapitulate the normal development of permanent cartilage than MSCs. Most of the published global high-throughput transcriptome data on the regulation of human chondrogenesis are from MSC or iPSC differentiation, which have the potential disadvantage of the type of cartilage produced (fibrous cartilage) and lack of spontaneous chondrogenic differentiation. The advantage of an embryonic LMP-based micromass model system is that it does not require additional growth factors or other reagents to achieve better chondrogenic differentiation than MSCs or iPSCs. Having performed a detailed analysis of the genes identified in this dataset, we not only confirmed the known components of chondrogenesis in this model but also provided an integrated view of refined inter-regulatory networks between the chondrogenic subset of genes.

#### Limitations of the Study

There are known differences between chicken and mammalian cartilage (83), which should be considered when applying these data to mammalian or human chondrogenesis. Careful *in silico* analyses should be conducted when selecting candidate genes from this pool. Furthermore, while we studied the effects of transient gene silencing of two transcription factors (*ATOH8* and *EBF1*) using siRNA electroporation, we acknowledge that not ideal transfection efficiency in micromass cultures, as well as the gradually diminishing effects of transient gene silencing over time could have influenced our results. For a more complete picture, it is important to monitor the effects of *ATOH8* and *EBF1* overexpression in this setting. Furthermore, it should be noted that this was an *in vitro* model of cartilage formation along with its inherent limitations. Therefore, the targets identified using this model must be validated *in vivo*.

### Perspectives

The ultimate goal of successful cartilage tissue engineering is to produce cartilaginous tissue with characteristics reminiscent of articular cartilage. This is a complex challenge; in addition to finding the right cell source, appropriate and time-dependent stimuli with exogenous factors are required. However, these cannot be successfully applied until the complex regulatory networks that control hyaline cartilage formation have been fully explored. The results of this study can provide a solid basis for fine-tuning and improving the current chondrogenic differentiation protocols.

## DATA AVAILABILITY

The entire dataset has been deposited and published in the BioProject database (http://www.ncbi.nlm.nih.gov/bioproject/). BioProject IDs: PRJNA817177 (day 0–day 6 micromass cultures); PRJNA938813 (day 10 and day 15 micromass cultures; mature articular chondrocytes). All other data generated or analysed during this study are included in this published article (and its supplementary information files).

## SUPPLEMENTARY DATA

Supplementary Data (Supplementary Files 1 and 2) are available online.

## AUTHOR CONTRIBUTIONS

Roland Á. Takács: Conceptualization, Formal analysis, Methodology, Validation, Visualisation, Writing—original draft. Judit Vágó: Conceptualization, Formal analysis, Methodology, Validation, Visualisation, Writing—original draft. Szilárd Póliska: Formal analysis, Methodology, Validation, Visualisation, Writing—original draft. Peter N. Pushparaj: Formal analysis, Methodology, Validation, Writing—original draft. László Ducza: Formal analysis, Methodology. Patrik Kovács: Formal analysis, Methodology, Visualisation. Eun-Jung Jin: Formal analysis, Writing—review & editing. Richard Barrett-Jolley: Formal analysis, Software, Methodology, Visualization, Writing—review & editing. Róza Zákány: Funding acquisition, Writing—review & editing. Csaba Matta: Conceptualization, Formal analysis, Funding acquisition, Methodology, Supervision, Validation, Writing—original draft, Writing—review & editing.

## List of abbreviations

AB: Alcian blue
BSA: bovine serum albumin
DEG: differentially expressed gene
DMMB: dimethyl methylene blue
ECM: extracellular matrix
FDR: false discovery rate
GO: gene ontology
HDC: high density culture
iPSC: induced pluripotent stem cell
LMP: limb bud mesenchymal progenitor
LFC: log2 fold change
MSC: mesenchymal stem cell
NGS: next-generation sequencing
NT: non-targeting control
OA: osteoarthritis
PBS: phosphate-buffered saline
PCA: principal component analysis
SafO: safranin-O
QC: quality check
RNA-seq: RNA sequencing
TF: transcription factor
TPBS: Tris-phosphate buffered saline
UMAP: uniform manifold approximation and projection
WGCNA: weighted gene correlation network analysis.

## ACKNOWLEDGEMENTS

The authors thank Mrs. Krisztina Biróné Barna for technical assistance. The authors are exceptionally grateful to Ms. Éva Szabó and Mrs. Ilona Huszár-Újvári Istvánné, and especially to Mrs. Katalin Váriné Debrecen (Katika Néni) for their kind assistance in handling and sacrificing broilers for articular cartilage collection. Without their involvement, a significant part of this study would not have been possible. We also wish to thank Dr. Peter Nagy at the Department of Biophysics and Cell Biology, University of Debrecen, for developing MATLAB-based image analysis software. We wish to thank Dr. Gergely Nagy at the University of Debrecen for useful comments, discussions, and critical feedback on the manuscript before submission. We are indebted to Ms. Ebeid Rana Abdelsattar Mansour for proofreading this manuscript. We would also like to thank Mr. Ferenc Henrik from Kelet-Grain Kft. poultry farm in Hajdúböszörmény (Hungary) for providing us with broilers for this study. Ethics License Number: HB/15-ÉLB/03743-2/2022.

## Footnotes

1 https://apr2022.archive.ensembl.org/Gallus_gallus/Info/Annotation

2 http://cole-trapnell-lab.github.io/monocle-release/docs_mobile/ (Last accessed: 24 February 2023)

