## Supplementary File 1 for "The temporal transcriptomic signature of cartilage formation"

### This PDF file includes:


- **Figure S1.** Time course of mitochondrial activity (cell viability) of micromass cultures throughout the culturing period
- **Figure S2.** Three top interacting partners with the most similar expression patterns to the transcription factors EBF1 and ATOH8, as retrieved from the STRING database
- **Figure S3.** Reactome pathway enrichment analysis on the set of genes identified using WGCNA analysis

- **Table S1.** Forward and reverse primer sequences, gene accession numbers and amplicon length for each primer pair employed in this study
- **Table S2.** The top 20 most abundantly expressed genes for each culturing day during chondrogenic differentiation
- **Table S3.** Qiagen IPA analysis carried out on pairwise comparisons of the entire dataset for the significantly overrepresented non-directional networks
- **Table S4.** Temporal expression profiles of genes encoding collagen subunits during *in vitro* chondrogenesis and mature articular chondrocytes
- **Table S5.** Temporal expression profiles of transcription factors known to regulate *SOX9* transcription during chondrogenesis

### Time course of mitochondrial activity (cell viability) of micromass cultures throughout the culturing period

#### *Materials and Methods*

For mitochondrial activity (cell viability) assays, micromass colonies cultured in 96-well plates were used. 10 µL of MTT reagent (3-[4,5-dimethylthiazolyl-2-yl]-2,5-diphenyltetrazolium bromide; Cat. No. 0793-1G; VWR, Avantor; 5 mg MTT/1 mL PBS) was pipetted into each well on different culturing days (0, 1, 2, 3, 4, 6, 10 and 15). Cells were incubated for 2 h at 37°C in a cell culture incubator, and following the addition of 500 µL MTT solubilizing solution, optical density was measured at 570 nm (Chameleon, Hidex, Finland). The assays were carried out on three biological replicate experiments, measuring 6 cultures each time. Data shown are mean ±SD. Statistical analysis between consecutive culturing days was carried out using Student's unpaired *t*-test. Asterisk (*) denotes significant (*$P<0.005$) difference between consecutive days.

Our results show that there is a rapid increase in cell numbers during the first 4 days of culturing, as reflected by a significant elevation of the MTT assay readings between consecutive culturing days. The rate of proliferation then declines, reaching a plateau between days 4 and 10, followed by a significant reduction in cellular mitochondrial activity by day 15 *(Figure 1)*.

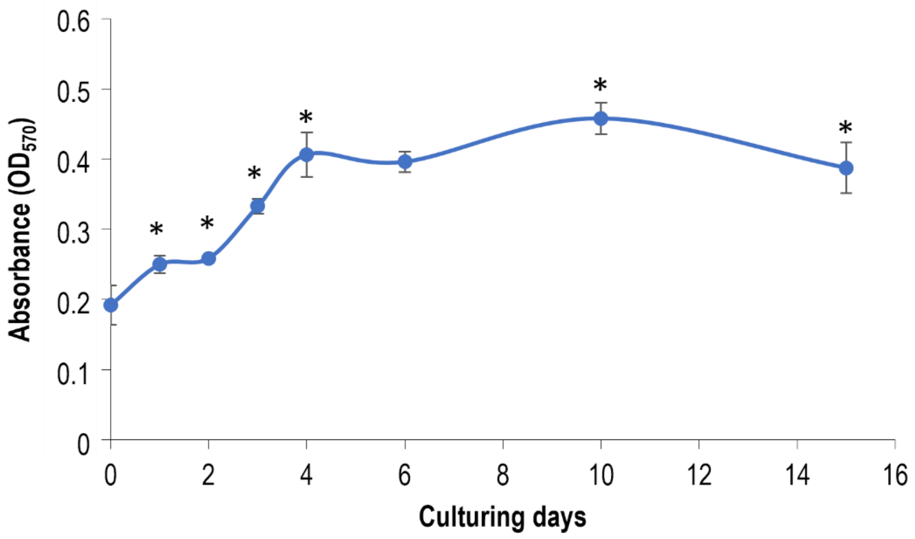


**Figure 1.** Time course of mitochondrial activity (cell viability) of micromass cultures on different days of culturing. Data points are average values of 6 biological replicates ±SD.

*Figure S2*. Three top interacting partners with the most similar expression patterns to the transcription factors *ATOH8*, *EBF1*, *NFIA*, *NFIX*, *PLSCR* and *STAT1*, as retrieved from the Signor (*NFIA*, *NFIX* and *STAT1*) and STRING (*ATOH8*, *EBF1* and *PLSCR1*) databases.

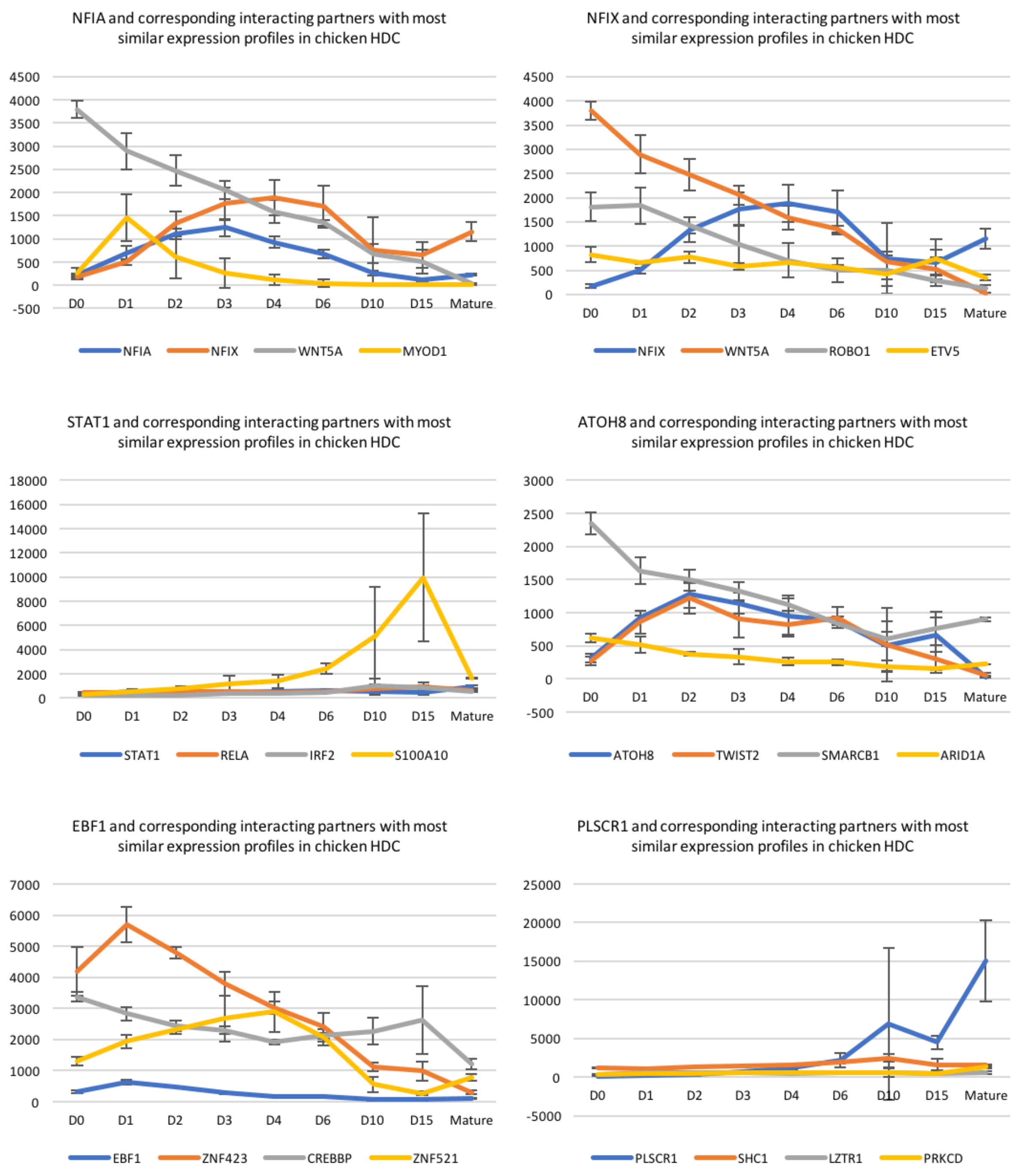

*Figure S3.* (**A**) Reactome pathway enrichment analysis results on the set of genes identified using WGCNA analysis. (**B**) Reactome pathway enrichment analysis results on the common entities (referred to as the 'core' chondrogenic subset) between the two chondrogenic models. Enrichment analysis was carried out using WebGestalt[1].

**A.**

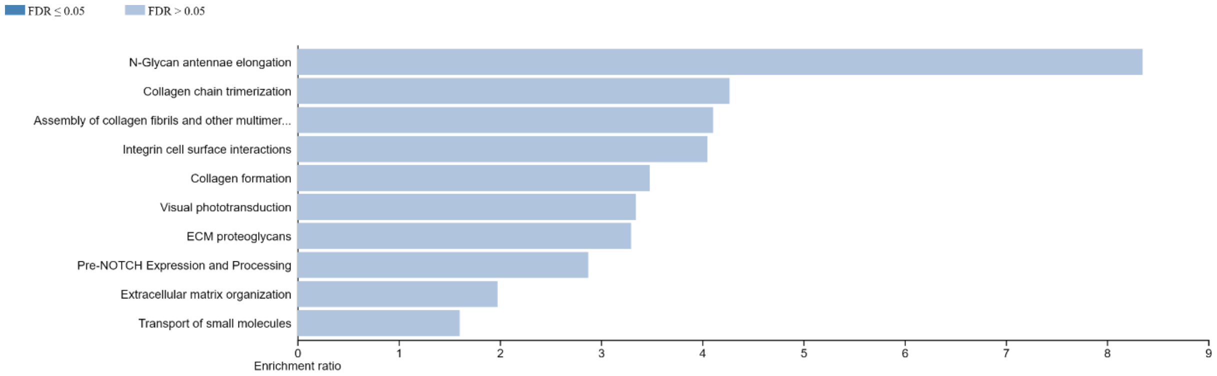


**B.**

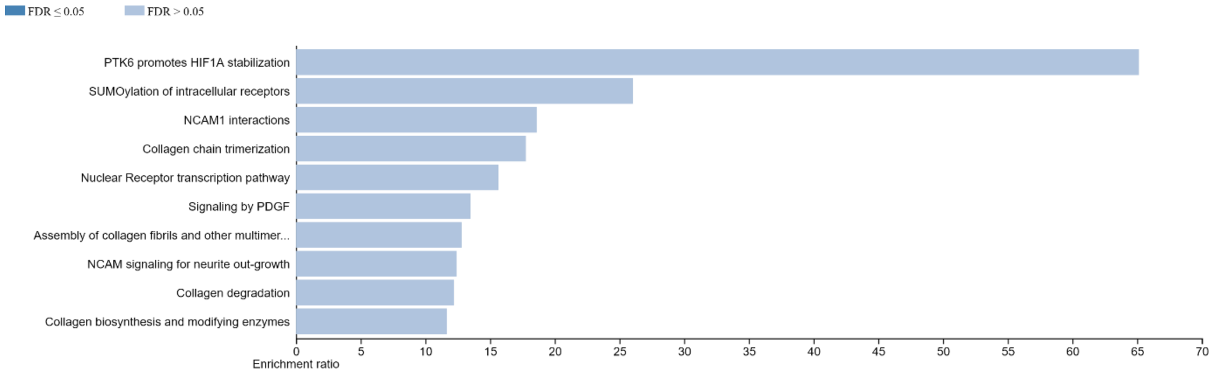


[1] WebGestalt (WEB-based Gene SeT AnaLysis Toolkit); www.webgestalt.org

*Table S1.* Forward and reverse primer sequences, gene accession numbers and amplicon length for each primer pair employed in this study.

| Gene symbol, species | Accession number | Primer sequence | Product length (bp) |
|---|---|---|---|
| ***RPS7*** *(Gallus gallus)* | XM_004940515.4 | FW: CGGAGTGCCGCGAAAGG<br>REV: TGTGATGTTCAACTCCCGCA | 157 |
| ***YWHAZ*** *(Gallus gallus)* | NM_001031343 | FW: GTTCCCTTGCAAAAACGGCT<br>REV: GAGGCAGACGGAAGTTGGAA | 199 |
| ***PPIA*** *(Gallus gallus)* | NM_001166326.1 | FW: GAGCTCTTCGCTGACAAGGT<br>REV: GCGTAAAGTCACCACCCTGA | 139 |
| ***HPRT1*** *(Gallus gallus)* | NM_204848.1 | FW: TGGTGGGGATGACCTCTCAA<br>REV: TCCAACAAAGTCTGGCCGAT | 190 |
| ***RPL4*** *(Gallus gallus)* | NM_001007479 | FW: TGTTTGCCCCAACCAAGACT<br>REV: CTCCTCAATGCGGTGACCTT | 137 |
| ***RPL13*** *(Gallus gallus)* | NM_204999.1 | FW: GGCCCGTGTTATCTCAGAGG<br>REV: CCGCTTCTTTGGCACGTTTT | 113 |
| ***SOX9*** *(Gallus gallus)* | NM_204281.1 | FW: TTTCCGAGACGTGGACATCG<br>REV: GTACCGCTGTAGGTGGTGAC | 150 |
| ***ACAN*** *(Gallus gallus)* | NM_204955 | FW: AGCAGTAGATGCACTGGGAC<br>REV: GCCAGGTCGATCTCACACAG | 153 |
| ***COL2A1*** *(Gallus gallus)* | NM_204426.1 | FW: GGGACCTCAAGGCAAAGTCG<br>REV: TTCCAGGCTCACCATTAGCG | 140 |
| ***HAPLN1*** *(Gallus gallus)* | XM_046934898.1 | FW: TTTCCCTCAATCAGGTCCTCT<br>REV: TCACAAGAATCTTCTTCACTTGTGT | 194 |
| ***COL10A1*** *(Gallus gallus)* | NM_001396427.1 | FW: TCCACAACATTTGAGGACGGA<br>REV: TTCACCCCTCATCTGCACAC | 192 |
| ***COL1A1*** *(Gallus gallus)* | NM_001396622.1 | FW: TACTGCAACATGGAGACGGG<br>REV: CCGCCGTACTCAAACTGGAA | 152 |
| ***RUNX2*** *(Gallus gallus)* | NM_204128 | FW: GAGTCAGATTACAGACCCCAGG<br>REV: AAATGGGCCCAGCTCGGAA | 199 |
| ***ATOH8*** *(Gallus gallus)* | XM_040671707.2 | FW: CTGGCCAGACTAGCAGACCT<br>REV: GTAAGGTCCTAGTGCACTGCT | 88 |
| ***EBF1*** *(Gallus gallus)* | XM_046926830.1 | FW: GCGACGATTTCAGGTTGTGG<br>REV: GCATGTTCCAGATAAGAAGGCG | 156 |

*Table S2.* The top 20 most abundantly expressed genes for each culturing day during chondrogenic differentiation, excluding not yet identified transcripts. Mean raw normalised expression data shown. Gene names in ***bold italics*** over a blue background have a known role in cartilage development (GO:0051216).

| ***Day 0*** | | ***Day 1*** | | ***Day 2*** | | ***Day 3*** | | ***Day 4*** | | ***Day 6*** | |
|---|---|---|---|---|---|---|---|---|---|---|---|
| **Gene symbol** | **Expression value** | **Gene symbol** | **Expression value** | **Gene symbol** | **Expression value** | **Gene symbol** | **Expression value** | **Gene symbol** | **Expression value** | **Gene symbol** | **Expression value** |
| ACTB | 104874.01 | ACTB | 76682.98 | ACTB | 73632.77 | ***COL2A1*** | 75390.25 | ***COL2A1*** | 123751.95 | ***COL2A1*** | 238898.16 |
| EEF1A1 | 68229.04 | EEF1A1 | 42814.84 | GAPDH | 55874.90 | ACTB | 74391.11 | ***COL1A1*** | 69871.14 | ***COL1A1*** | 149234.53 |
| LOC776816 | 63438.22 | LOC776816 | 40131.43 | EEF1A1 | 44357.99 | GAPDH | 54539.84 | ACTB | 63497.41 | ***COL1A2*** | 70243.33 |
| YBX1 | 36022.98 | GAPDH | 34126.15 | ***COL2A1*** | 43847.94 | ***COL1A1*** | 48997.35 | GAPDH | 59476.00 | GAPDH | 59150.92 |
| NPM1 | 35274.93 | YBX1 | 22678.09 | LOC776816 | 43562.14 | LOC776816 | 44292.92 | ***PTN*** | 50089.40 | ***COL9A2*** | 56094.60 |
| EEF2 | 33655.45 | EIF4G2 | 22241.27 | ENO1 | 27525.70 | EEF1A1 | 39951.72 | ***COL1A2*** | 41891.96 | ACTB | 50600.52 |
| GAPDH | 32425.10 | EEF2 | 21606.37 | YBX1 | 26247.79 | PTN | 33872.99 | LOC776816 | 40783.76 | LECT1 | 47403.40 |
| EIF4G2 | 30031.09 | HNRNPA3 | 19656.51 | ***COL1A1*** | 24522.39 | ***COL1A2*** | 31592.26 | EEF1A1 | 35100.54 | ***COL9A1*** | 46777.57 |
| HNRNPA3 | 29386.52 | ***COL2A1*** | 18971.79 | ***PTN*** | 21718.57 | ENO1 | 28158.53 | ENO1 | 32030.89 | HSPA5 | 46090.14 |
| TUBA1A | 28331.81 | TUBB | 18874.47 | RPS2 | 21494.02 | YBX1 | 27719.89 | LECT1 | 30558.46 | FTH1 | 45947.39 |
| NCL | 27493.24 | ENO1 | 18641.04 | RPS3A | 21146.14 | MDK | 22718.83 | YBX1 | 28453.24 | CST3 | 41480.22 |
| HSP90AA1 | 27469.91 | CHGB | 18351.30 | EEF2 | 20750.29 | RPS3A | 22110.30 | RPS3A | 27908.53 | ***ACAN*** | 40587.91 |
| PPIA | 27397.40 | FSTL1 | 18303.51 | RPS6 | 20590.98 | ***FN1*** | 21985.02 | ***COL9A2*** | 27228.59 | ***SPARC*** | 38280.12 |
| TUBB | 26043.80 | RBM3 | 17386.09 | TUBB | 19993.14 | RPS2 | 21756.07 | FKBP9 | 25823.61 | ***PTN*** | 37349.72 |
| RPS3A | 26019.32 | HSPA8 | 17306.93 | H3F3B | 19504.65 | RPS6 | 21330.40 | SERPINH1 | 25343.13 | FKBP9 | 36434.41 |
| RPS2 | 25109.70 | RPS3A | 16813.27 | ***COL1A2*** | 19275.63 | FSTL1 | 20748.00 | FTH1 | 24588.54 | EXFABP | 36273.92 |
| RPSAP58 | 24311.93 | RPS6 | 16723.76 | EIF4G2 | 18065.15 | MYL6 | 20206.74 | MDK | 24445.92 | SERPINH1 | 35671.97 |
| RBM3 | 24153.97 | NPM1 | 16645.47 | FSTL1 | 18019.00 | RPS12 | 19625.07 | ***COL9A1*** | 24243.56 | LOC776816 | 35179.68 |
| RPS6 | 23158.93 | H3F3B | 16644.55 | HNRNPA3 | 17617.66 | H3F3B | 19259.02 | RPS6 | 24136.15 | HSP90B1 | 34316.00 |
| HNRNPA1 | 22610.10 | PPIA | 16031.69 | TUBA1A | 17197.45 | EEF2 | 18911.18 | RPS12 | 23343.51 | RPS3A | 32627.60 |

| ***Day 10*** | | ***Day 15*** | | ***Mature*** | |
|---|---|---|---|---|---|
| **Gene symbol** | **Expression value** | **Gene symbol** | **Expression value** | **Gene symbol** | **Expression value** |
| SAA | 322365.47 | HSPA2 | 266896.48 | ***COL2A1*** | 1419262.40 |
| ***EXFABP*** | 240747.94 | FTH1 | 229047.25 | SAA | 720922.95 |
| ***COL2A1*** | 198944.63 | HSP90AA1 | 176898.66 | ***SPARC*** | 488806.06 |
| FTH1 | 197572.53 | HSPB9 | 139746.30 | AVD | 350749.92 |
| ***COL1A1*** | 129367.40 | HSPA5 | 101269.10 | ***EXFABP*** | 341789.89 |
| HSPA5 | 94803.96 | ***COL2A1*** | 91235.87 | ***COL1A1*** | 316190.19 |
| HSPA2 | 91205.07 | LOC101747587 | 87306.70 | FTH1 | 305357.61 |
| LOC101747587 | 85068.42 | PRRC2C | 75354.07 | HSPA5 | 299344.97 |
| GAPDH | 84769.15 | GAPDH | 74156.31 | ***FN1*** | 267380.94 |
| ***ACAN*** | 77578.68 | ***COL1A1*** | 73460.24 | MMP10 | 220744.73 |
| SPP1 | 74638.70 | RPS3A | 69913.49 | LOC101747587 | 211266.60 |
| IL8L2 | 70642.08 | ***SPP1*** | 69657.21 | EEF1A1 | 159232.35 |
| ***COL6A3*** | 59496.75 | EEF1A1 | 67160.13 | ***COL1A2*** | 141878.95 |
| HSPB9 | 59200.89 | ACTB | 63226.85 | SPP1 | 137723.37 |
| PTGS2 | 57887.17 | PNRC1 | 58667.96 | ***DCN*** | 132089.72 |
| ***COL9A1*** | 57855.93 | HSPA8 | 55281.88 | HSPB9 | 129236.76 |
| ***COL9A2*** | 57042.83 | EIF5B | 52908.45 | ***COL9A1*** | 103560.60 |
| ***COL1A2*** | 55431.53 | PABPC1 | 52569.23 | ***COL9A2*** | 92938.82 |
| HSP90AA1 | 54449.96 | RPS12 | 52559.78 | HSPA2 | 86708.39 |
| LDHA | 48782.14 | TMSB4X | 52538.43 | ***SERPINE2*** | 80261.53 |

*Table S3.* Qiagen IPA analysis carried out on pairwise comparisons of the entire dataset for the significantly overrepresented non-directional networks.

| Top diseases and functions | Molecules in Network |
|---|---|
| *Day 0 vs 1* | |
| Embryonic Development, Organismal Development, Tissue Development | ACTBL2,ALDH1A2,ALDH1A3,Alpha tubulin,ANKRD33,BCAS1,BHMT2,BMP8A,CASQ2,Ck2,FAM3D, FOXC1,Insulin,ISL2,ITGBL1,LMO1,MEF2C,MICAL1,MYO1F,OLIG2,OSTN,RNA polymerase II, SLC2A9,SLC35F3,SRL,TAL1,TAL2,Tgf beta,TRPM8,TRPV2,TUBB3,VGLL2,VGLL4,VWA1,WNT16 |
| Cancer, Cell-To-Cell Signaling and Interaction, Organismal Injury and Abnormalities | ADAMTS7,Adaptor protein 2,ADGRD1,ADGRF5,ADORA1,AREG,ARR3,BTN1A1,chemokine, CHIA,EDN2,EFEMP1,EGFR ligand,Erbb1 dimer,GASK1B,Girk,Gpcr,GPER1,GPR171,HAVCR1, KCNJ12,Metalloprotease,Mmp,MMP27,NDP,NMBR,NPY4R/NPY4R2,NR4A3,RGS20,SCTR,SMC1B, Tnf (family),UTS2R,Ve Cadherin,Vegf |
| Amino Acid Metabolism, Cell-To-Cell Signaling and Interaction, Molecular Transport | ADAM32,ATP12A,C1QB,CCR6,CLDN8,FA2H,GADL1,GAS6,GPR182,GPRC5D,HSPA2,KCNA2, KCNE3,KCNG1,LAPTM4B,MALL,MLNR,NTSR1,ODF4,OPN1LW,OPRL1,RARS2,REEP6,SCT, SLC30A2,SLCO4C1,SYNE4,SYT2,TJP3,TM9SF1,TMEM182,TMEM71,TRARG1,VIPR1,WFDC11 |
| Cardiovascular Disease, Hereditary Disorder, Skeletal and Muscular System Development and Function | ACTC1,ATPase,CHRDL2,CMYA5,Collagen type III,CRYAB,EEF1A,EEF1A2,G Protein I,GDF5,HSP, Hsp70,HSPB2,HSPB9,Kallikrein,LDL-cholesterol,LRRC56,MYBPC3,MYL1,MYL11,MYL2,MYL3, MYOM1,NEU4,P38 MAPK,PLAT,RBM24,RGS5,Rlc,SERPINB5,TNNI1,TNNT1,TNNT2,tubulin (family),VLDL-cholesterol |
| Cancer, Gastrointestinal Disease, Hepatic System Disease | ACTA2,ARRDC4,CER1,DDX4,DEUP1,EOMES,FGFR3,FOXA1,Histone h2a,Histone h3,Histone h4,Hsp90,Ifn gamma,Ikb,IKK (complex),IL2RB,IRX2,KCNK4,KDR,KIT,NR4A1,OTP,PCK1,PEPCK, PLAAT1,PTK,PTN,Rsk,SP9,SYTL5,TFAP2A,Ubiquitin,Wnt,WNT2,ZIC1 |
| *Day 1 vs 2* | |
| Behavior, Cardiovascular Disease, Organismal Injury and Abnormalities | Actin,ARC,BGN,Calcineurin protein(s),caspase,Collagen type I (complex),collagen type I (family), CTLA4,ERK,F Actin,FAM83B,FGFBP2,Fibrinogen,FOXD1,GLRX,H4C4,Hdac,HISTONE,Ige,IgG, Immunoglobulin,LCP2,LDB3,MB,NFAT (complex),NFkB (complex),PAX7,PP2A,Rap1,SCIN, SFRP2,TCR,TIMD4,TREM2,VWF |
| Connective Tissue Disorders, Organismal Injury and Abnormalities, Skeletal and Muscular Disorders | ADAMTS3,AHSG,Akt,BRINP2,CA4,Casein,CETP,COCH,collagen,Collagen type II,Collagen type III, Collagen type IV,Collagen(s),Ecm,EPYC,Fibrin,HDL-cholesterol,Integrin,INTERLEUKIN,KCNE3,LDL, LDL-cholesterol,Metalloprotease,Mmp,MMP1,MMP10,MMP27,MMP7,PAPPA2,PDGF BB,PLC, PLD,PRKAA,secreted MMP,TRPV4 |
| Carbohydrate Metabolism, Molecular Transport, Small Molecule Biochemistry | ADCY,Ap1,AVP,BDKRB2,CAV3,CD3,CG,chemokine,Creb,cytokine,ERK1/2,G protein alpha i,Gpcr,IL-1R,IL1,IL12 (complex),IL12 (family),IL17D,IL1RL1,Lh,Mapk,MYF5,Nfat (family),Nos,NRIP2,P38 MAPK, PADI3,Pka,Pro-inflammatory Cytokine,RAMP3,RGS3,Rxr,Tgf |
| Cellular Assembly and Organization, Humoral Immune Response, Inflammatory Response | AFAP1L1,AGXT,C1q,C1QC,C1QL1,C1QTNF8,C3,Ck2,COL14A1,COMP,Complement component 1, Complement System,F13B,FCN3,FDX2,FNDC1,HOXC12,HPX,inosine,LAIR2,LOXHD1,MATN1,MCF2L, GGGCUGG),MYL2,PINLYP,PPP1R11,PPP1R1C,RASGEF1C,RSPH6A,SCX,SYCN,ZNF575 |
| Gastrointestinal Disease, Hepatic System Disease, Liver Cholestasis | 2-hydroxyestrone,acetaldehyde,ADH1C,C6orf58,CCL17,chymotrypsin,CONNEXIN,CRLR-RAMP3 (complex),CYP3A5,CYTIP,Cytokeratin,GJB2,Histone h3,IL17R,INPPL1,Insulin,Jnk,KCNJ15,LRP, myosin-light-chain kinase,OSTN,p85 (pik3r),PI3K |
| *Day 2 vs 3* | |
| Cell to Cell Signaling and Interaction, Cellular Growth and Proliferation, Hematological System Development and Function | ADAM19, ADH6, BOLL, C1QTNF8, CAPN9, CHODL, CSF3, FAM180A, FGFBP2, FUT9, GABA receptor, GABRA4, HBZ, HELT, IKZF3, IL4, MATN1, NRIP2, PI16, R3HDML, RGS11, SLC26A4, SP8, Timp, UPK3A, XK, ZFPM1 |
| Connective Tissue Development and Function. Nervous System Development and Function, Reproductive System Development and Function | ADIPOQ, Alp, AMPK, BGN, C1q, CG, chemokine, COCH, Collagen Alpha1, Creb, CRH, FSH, GJB2, GPER1, HBZ, HDL, IFN Beta, IgG,IL12 (complex), IL23, IL6, Immunoglobulin, Lh, NfkB1-RelA, Nos, ORMI, PENK, PEPCK, PINLYP, Pka, PRKAA, PRLH, Pro-inflammatory Cytokine, Sod, voltage-gated calcium channel |
| Infectious Diseases, Organismal injury and Abnormalities, Psychological Disorders | Ap1, AQP3, BCR (complex), CA12, CA4, Carbonic Anhydrase, CD3, CDKN2B, cytokine, ERK, FRY. H4C4, HISTONE, Histone h3, IFN alpha/beta, Ige, Igm, Ikb, IKK (complex), IL12 (family), Interferon alpha, JAK3, JUN/JUNB/JUND, NFkB (complex), Notch, PI3K (complex), RNA Polymerase II, SLC6A2, SOX21, STAT, STAT5a/b, TCR, TLR7, Tnf(family), TNFSF11 |
| Connective Tissue Disorders, Developmental Disorder, Hereditary Disorder | ALKAL2, Alpha catenin, AQP9, BANCR, Calmodulin, caspase, COL14A1, COLEC10, collagen type I (family), cysteinyl-leukotriene, FIBRINOGEN (family), FOXA1, Histone h4, HTR1A, IL15R Insulin, IRS1/2. Jnk, Kng1/Kng2, MAP2K1/2, Mapk, marinobufagenin, MASP1, Mek, NMDA Receptor,NPTX1, P38 MAPK, PI3K p85, Pkc(s), POMC, RAS, Smad2/3, Sos, SOX7, Vegf |
| *Day 3 vs 4* | |
| Endocrine System Disorders, Gastrointestinal Disease, Hereditary Disorder | ADGRG7, AQP5, BRDT, Calcineurin protein(s), Calmodulin, Ck2, EOMES, FBX041, FOXA2, GALR3, GATA4, Gpcr, GPR35, GRM5, Hedgehog, HISTONE, Histone h3, IKZF1, Insulin, KCNJ11, KEL, LEFTY1, MLKL, NFkB (complex), PIFO, RGS13, RNA polymerase II, RXFP3, SLC8A2, SPP1, TYR, Wnt, WNT7B, WNT8A, XK |
| Cell to Cell Signaling and Interaction, Cellular Movement, Hematological System Development and Function | CHAD, CTXN3, DEFB4A/DEFB4B, ERK1/2, GZMK. HES6, IFN, IFN Beta, IgA, IgG, IgM, IL1, IL12 (complex), IL12 (family), IL20, IL21R, IL36B, JAK, JAK3, KL, LDL, MARCO, OMD, PIGR, PRKAA, Pro-inflammatory Cytokine, Serine Protease, SPINK5, STAT, STAT4, TFAP2E, Tgf beta, TMPRSS2, Tnf (family), TRAT1 |
| Cell to Cell Signaling and Interaction, Drug Metabolism, Molecular Transport | ACP3, ADCY, AKAP14, Akt, ANGPTL3, ASAH2, CA12, cAMP-dependent protein kinase, Carbonic anhydarase, CG, CGA, CNGA1, Creb, CRH,DUOX1, estrogen recepror, FILIP1L, FOXG1, FSH, GNRH, Growth Hormone, Hdac, LDL-cholesterol, Lh, IPDGF BB, Pka, Pka catalytic subunit, PLC, POU1F1, POUF4F1, RIT2, SLC5A8, TCP11, TH, TSH |
| Inflammatory Response, Lipid Metabolism, Small Molecule Biochemistry | ANKRD60, BARHL2, CAMK4, CD14, CELA1, CIB3, CREB1, ganglioside GD1a, GPR26, HBEGF, histamine, IGSF9B, IL17C, KY, Ldha/RGD1562690, MRCHF4, MATN1, NOD2, OVOL3, PLA2G12B, PLD4, PRDM12, prostaglandin D2, REL, resolvin D2, RNA fragment, SFTPD, SIAH3, SPHK1, SYCP2L, TACR3, TLR6, TRAF3IP2 |

*Table S3.* (continued)

| | |
|---|---|
| *Day 3 vs 4 (continued)* | |
| Cell to Cell Signaling and Interaction, Hematological System Development and Function, Immune Cell Trafficking | ADAM17, CAMK4, CARD6, CD14, CD200R1L, CLDN18, CLEC3A, CNDP1, CTSL, CTSS, ECEL1, FAM180A, GALNTL6, HBEGF, IL17C, IL2, Integrin alpha 5 beta 3, Integrin alpha V beta 3, KCNV1, Mlc, PPP3CA, prostaglandin D2, RIGI, RNF212, SLC13A1, SPON2, SPP1, SRC, TLR5, TLR6, TSPAN32, UNC5D, Villin |
| *Day 4 vs 6* | |
| Nervous System Development and Function, Neurological Disease, Organismal Injury and Abnormalities | ARHGEF38, CBLN3, CCDC85C, CDH33, CELSR3, CLEC3A, CPEB2, DIS3L, DMRTA2, DMRTB1, EXTL1, FAM180B, GBX1, Gef, GRB2, KCNA6, KRT81, KRTAP19-5, LCN15, Muc1, NOTO, OTX1, PCDH12, PRR22, SLC15A1, SLC3SF1, SPRY3, SV2C, TMCC3, TPBGL, UBL5, ZMAT5 |
| Cell-To-Cell Signaling and Interaction, Decreased Levels of Albumin, Tissue Morphology | ADCY, ADGRF4, APLNR, BDKRB1, BDKRB2, Calmodulin, Casein, CG, Creb, CRH, ERK, ERK1/2, G protein alpha i, Gpcr, GPR50, Gsk3, Hdac, IL12 (family), Integrin, Lh, LRRN3, Mapk, MC1R, Metalloprotease, Mmp, MMP7, Pka, RGS20, RGS3, SCN2A, STAT, TCF, TLR7/8, TRPM3 |
| Cellular Development, Cellular Movement, Hepatic System Development and Function | ANKRD22, Ap1, ATP6VOD2, CCR7, COMP, cytokine, ELOVL3, F10, Rbrinogen, GOT, HISTONE, IFN Beta, Ifn gamma, Ifnar, Iga, Ige, IgG, Igm, IL12 (complex), 1L15, Immunoglobulin, Interferon alpha, KCND3, MHC Class II (complex), NFkB (complex), PROX1, RIMS4, RIPK4, Rxr, SFRP5, SPP1, Tgf beta, Tlr, Tnf (family), transcription factor |
| Cellular Function and Maintenance, Cellular Movement, Tissue Morphology | Akt, Alp, C/EBP, C4A/C4B, CD55, CLSTN2, Collagen type II, Collagen(s), collagenase, CSF, elastase, GHRL, Growth hormone, HDL, HDL-cholesterol, IL-1R, IL1, IL22, IL23, 1L4I1, LDL, MMP1, NfkB1-RelA, Nr1h, ORM1, PDGF BB, PRKAA, Pro-Inflammatory Cytokine, RELN, SAA, SAA1, Secretase gamma, SLC15A1, Sod, UCN |
| *Day 6 vs 10* | |
| Cell Signaling, Cellular Development, Embryonic Development | ADGRB1, ANKRD1, ARHGAP22, BDKRB1, BNC1, C6orfS8, DENND2D, Endothelin, ERO1A, GDF2, GHSR, GJA4, Gpcr, GRPR, HAND1, HHEX, HOPX, HRH3, HTR1E, HTR6, KIT, LPAR6, MMRN1, NKX2-3, NMBR, OSR1, OSR2, PGF, RASL11B, Relaxin, RXFP1, RXFP2, SCGN, SPDEF, Vegf |
| Cell Morphology, Cellular Assembly and Organization, Cellular Development | 26s Proteasome, ASIC2, C15orf39, CHL1, CP, CPNE7, CRHBP, DAB1, DIPK1C, DPEP2, ESRP2, FAM81A, FOXF2, GABRA2, GADL1, GCH1, Hsp70, Hsp90, HSPA2, IKKA/B, KCNAB1, MLKL, NFKBIA, NLGN3, Nos, NWD2, P glycoprotein, Proteasome, RIPK3, SGK2, SNCA, SV28, TAL2, TC2N, Ubiquitin |
| Cell Death and Survival, Cellular Growth and Proliferation, Gastrointestinal Disease | ACSL1, AGR2, B3GAT1, B1RC3, C1QB, CARD11, caspase, CCDC85C, CCR2, CRNN, CYB561, DUSP8, EGLN3, ENO2, FTH1, Ikb, IKK (complex), ITM2C, LYZ, MAP1LC3, MBP, MFSD12, MHC Class II (complex), MHC II, MPP2, PARP14, PLP1, REEP6, RGS1,Sapk,SFRP2, SlCO4C1, SQSTM1, TH2 Cytokine, TNFSF15 |
| Cell-To-Cell Signaling and Interaction, Hematological System Development and Function, Tissue Morphology | ADORA2B, ANKRD22, Ap1, ATF3, BATF3, CD83, CEBPB, CEBPD, Collagen(s), DUSP1, FHL2, GPRIN2, Hif1, IFN alpha/beta, IFN Beta, IL12(family), ITGA2B, KERA, Laminin (family), MAPK, MAL, MMP1, MMP9, NELL2, NFkB (family), NFKBIZ, RNA polymerase II, SDC4, SELE, Smad2/3, SPP1, STEAP3, Tgfbeta, TMEFF1, TXNRD1 |
| *Day 10 vs 15* | |
| Cell-To-Cell Signaling and Interaction, Cell Movement, Hematological System Development and Function | 26s Proteasome, Alpha catenin, carboxypeptidase, CDHS, Collagen(s), CPA6, CTSA, DPEP1, EMB, EPYC, FBLNS, GDFtt, GDF2, HDL, Hsp70, KERA, HYZ, MMP9, MXRA5, NCAN, NFKBIA, Nos, PLEKHNt, PRKAA2, PTGSZ, RNA polymerase ll, HELE, STEAPJ, STEAP4, SYK/ZAP, SYT4, TAL2, THBS2, Trypsin, Ubiquitin |
| Dermatological Diseases and Conditions, Immunological Disease, Inflammatory Disease | Alp, C1QL3, CALML3, CYGB, cytochrome C, Cytokeratin, DOCK11, fgf, FGFR, GTPase, HEPHl1, ITGB1BP2, Keratin, KRT10, KRTS, KRT6A, KRT8, MEF2, ONECUT1, PHEX, P13K (complex), PKP1, PRKAA, PROTEASE, PTPRNZ, Raf, Rb, RSP01, SGIPt, SLC9A2, SUSOJ, TNFRSF19, TRIM29, UGT1A1, WNT11 |
| Connective Tissue Disorders, Organismal Injury and Abnormalities, Skeletal and Muscular Disorders | ACE, Adam, ADAM19, ADAM20, ADAMDEC1, ADAMTS17, ADAMTS18, ADAMTSJ, ADAMTS7, ADAMTS8, AQUAPORIN, chymase, Collagen type Ill, Collagen type IV, Collagen type V, Ecm, ERBB4, GABA A receptor, HAVCR1, IL1B, Integrin, laminin1, lamininS, LRG1, MAOA, Metalloprotease, MMP10, MMP27, NID1, NOX3, NRG (family), SLC01B1, somatostatin receptor, TCF/LEF,THBS4 |
| Gastrointestinal Disease, Neurological Disease, Organismal Injury and Abnormalities | ACKR3, ADGRF5, ADGRL3, ADRA1B, B3GALT1, CCR7, CCRLZ, CHODL, CHRDL1, CRHBP, DRD5, FAM180A, FAM83C, FOX13, GABBR2, GPR132, GPR5O, HRH1, HTR1B, IT1H5, KCNG3, KCNS1, KCNT2, MTX1, NMBR, NTMT2, PTGDR, PTH2R, SLC04C1, SSTR2, SSTR4, TMEM196, TMEMZ33, ZNG1C/2NG1F |
| Connective Tissue Disorders, Organismal Injury and Abnormalities, Skeletal and Muscular Disorders | AMPA/kainate receptor, Cadherin, CDH11, CDH23, CHRNA1, CHRNA3, COL13A1, COL17A1, COL1A2, COL28A1, COL8A1, COL8A2, collagen, Collagen type XVIII, FBLN2, glutamate receptor (family), GP11B-IIA, GRI, GRIA3, GRIK1, GRIK2, Hsp27, JINK1/2, LDB3, LDL cholesterol, MARCKSL1, Mmp, MMPtJ, NACHR, NEU4, P38 MAPK, Relaxin, SLC6A2, Tgf beta, Tnf (family) |
| *Day 15 vs Mature* | |
| Embryonic Development, Hematological System Development and Function, Lymphoid Tissue Structure and Development | ARHGAP22, BFSP1, BTBD17, CCR7, CD2, CUX2, EPS1SL1, FCHS02, FNBP4, GDA, HS3ST1, ITSN1, KDM1B, KIF168, Kif188, Kif3C, KPNA2, MDC51, MSL1, PRDM16, RAB17, RAB28, RUNDC3A, S100A14, S100A16, SIOOA6, SLC27A2, SPATA17, SRSF11, TMOD4, TOX, TOX2, UBN1, ZIC1, ZNF821 |
| Cancer, Gastrointestinal Disease, Organismal Injury and Abnormalities | AMPD3, ARAP3, ARF6, C11orf52, C1orf35, CCl4, CEMIP2, DACH1, DDX60, DEF8, DENND2D, EHBP1, EPB41L3, GCHFR, GSF3, 1TGA9, KIAA1549, MAP7D2, MDK, MLLT3, MSC, MYCT1, NEBL, NSUN7, PLEKHM3, PNISR, PNN, POLE4, PPIP5K2, RTF1, SIX2, SMOC1, TAFAS, TRIM55, VAX1 |
| Cancer, Cell-to Cell Signaling and Interaction, Nervous System Development and Function | ADD2, AHNAK2, ATP5MG, BAZ1B, CGN, COLEC10, FAM169A, FBXW7, FECH, IGF2BP1, IGF2BP3, INHBE, KIAA0753, LARP7, LRRCC1, MCU, MRPL18, MRPL58, MRPS5, MTG1, MTRFR, MY01B, MY01F, PIN4, PRKCB, RPLP2, SARS2, SKA3, SMC4, TOP2A, TPM1, TUBAL3, USF3, ZBTB47, ZNF746 |
| Cellular Development, Cellular Growth and Proliferation, Developmental Disorder | ANKFY1, ANXA1, ARHGAP36, ARHGAP39, BASP1, CENPA, CFAP70, CSAD, CTNNA3, DBF4B, DDX55, DPY30, EIF5B, EPB41L4A, FRK, H3·5, HDGF, HMGA2, IL2RA, KLHL6, MAP1B, MYCBP, OBSL1, PEBP1, PROSER2, PSIP1, RACGAP1, RC3H2, ROR2, SFN, SPIRE1, STMN2, THOC2, TRPC5, VOPP1 |

*Table S3.* (continued)

| | |
|---|---|
| *Day 0 vs Day 15* | |
| Developmental Disorder, Hereditary Disorder, Organismal Injury and Abnormalities | ALG9, ANO6, ATF6B, BYGNT7, CANX, CCPG1, CCR7, CLGN, CNGA3, CNPY3, DPY19L1, ECEL1, EMF1, ESF1, FAM177A1, GHDC, GPC3, GPR174, 1RAG2, LRRTM3, NHSL1, NRN1, NSDHL, PKD2, SEC11C, SE62, SOGA3, SPCS1, SPOCK1, SPOCK2, SPOCK3, SPTBN5, THSD7B,TMX2, UXS1 |
| Cancer, Cell Cycle, Cellular Development | AKR1D1, AURKB, CAMKV, CAPN6, CD4, CKMT2, DTL, ECT2, FOLR1, GART, GEN1, KBTBD3, KIF20A, KIF23, MAD2L1BP, MME, NUP210L, PCBP3, PCNX2, PFDN1, RBMXL3, RFESD, RIF1, RPL21, RPL38, RPLP1, RPLP2, SRSF3, SUGP2, TMEM140, TMEM182, TOP2A, TRIP13, VRK1, ZBTB14 |
| Cardiac Dilation, Cardiac Dysfunction, Cardiac Enlargement | A1CF, ADAMTSL1, ANO3, ARID4B, C10orf71, CCBE1, CCL4, CLSTN2, FBN2, FHL2, GRN, HAND2, 1RX5, ITGA9, KCNAB2, KIF21B, KLHDC88, LBX1, LOC1027Z4428/SIK1, MFAP5, MICALL2, MSC, MYL11, MYPN, NANOS1, NPAS2, P-TEFb, PALLD, PDLIM5, SPEG, SPINK2, TIMM10, UNC93A, WDCP, YAF2 |
| Cell Morphology, Cellular Assembly and Organization, DNA Replication, Recombination, and Repair | alcohol group acceptor phosphotransferase, ASCL3, ASTN2, BAIAP2L1, BBX, BFSP1, C12orf75, CADPS2, CDCA3, CDKN3, CENPE, DEPDC1, FOXM1, GRID2IP, KIF18B, KIF24, KIF2C, KIF3A, KIFC1, MTFR2, MYOD1, NEK2, NFATC2, NUSAP1, PAPLN, PARD6B, PLK1, RNF170, S100A6, SCGN, SPAG1, TMOD4, TRIM2, TRIP11, VPS36 |
| Cell Cycle, Digestive System Development and Function, Gene Expression | ABHD8, ACTL6A, BCL11B, C1QTNF8, CCNJ, CCNJL, CEBPB, COLEC10, DCLK1, DEPDC1B, DUT, ESPL1, FAM110B, Igh (family),INO80C, LRRN1, MELK, NETO1, NGEF, NMNAT3, PARD3B, PCDH17, PHLDB2, RAB31P, RICBB, SATB2, SCUBE3, SERPINH1, SIM2, STAG1, TIAM1, TRIM47, TUBAL3. VSX1, VWA2 |
| Cancer, Connective Tissue Disorders, Organismal Injury and Abnormalities | ADIPOQ, ARRDC4, BCKDHB, C1QTNF3, CAPN13, CCDC107, CD99, CKAP2, CLDN3, DSP, ECSCR, EMB, FADS6, FAM107B, FLI1, GBX1, HMGB2, HOXC8, HPD, IgG, LEF1, LIX1, MAP9, MBNL2, MBNL3, MEF2B, OSTN, PAX7, PLOD1, PLOD2, POU2AF1, POU3F1, RIPOR2, STXBP4, ZC3H1ZD |

*Table S4.* Temporal expression profiles (normalised average expression values) of genes encoding collagen subunits during *in vitro* chondrogenesis and in mature articular chondrocytes.

| Symbol | Day 0 | Day 1 | Day 2 | Day 3 | Day 4 | Day 6 | Day 10 | Day 15 | Mature |
|---|---|---|---|---|---|---|---|---|---|
| *COL1A1* | 12435.65 | 13731.73 | 24522.39 | 48997.35 | 69871.14 | 149234.53 | 129367.40 | 73460.24 | 316190.19 |
| *COL1A2* | 10277.92 | 12629.87 | 19275.63 | 31592.26 | 41891.96 | 70243.33 | 55431.53 | 13807.64 | 141878.95 |
| ***COL2A1*** | **5124.71** | **18971.79** | **43847.94** | **75390.25** | **123751.95** | **238898.16** | **198944.63** | **91235.87** | **1419262.40** |
| *COL3A1* | 520.47 | 1547.80 | 3374.46 | 3665.71 | 3883.68 | 6796.37 | 8038.21 | 4638.09 | 11164.50 |
| *COL4A1* | 957.98 | 1370.99 | 1694.11 | 2337.63 | 1648.97 | 1535.28 | 2749.17 | 1610.88 | 6033.96 |
| *COL4A2* | 558.78 | 723.58 | 883.45 | 1111.57 | 708.62 | 579.46 | 1415.55 | 667.37 | 2892.33 |
| *COL4A4* | 3.39 | 13.63 | 15.99 | 12.65 | 9.09 | 11.30 | 3.54 | 3.29 | 12.67 |
| *COL5A1* | 3354.38 | 6653.15 | 8581.59 | 13006.58 | 15960.85 | 22972.10 | 23911.12 | 19489.95 | 5915.29 |
| *COL5A2* | 3239.05 | 9148.61 | 14107.28 | 14828.89 | 15346.17 | 14203.64 | 13050.12 | 6892.85 | 6800.77 |
| *COL6A1* | 209.83 | 456.72 | 958.06 | 1531.47 | 2256.01 | 4691.38 | 8377.26 | 4378.88 | 35927.92 |
| *COL6A2* | 379.43 | 841.16 | 1493.59 | 2161.68 | 3493.79 | 10129.61 | 21534.14 | 8850.29 | 80238.39 |
| *COL6A3* | 517.26 | 1576.95 | 3110.91 | 6015.25 | 11650.26 | 26606.21 | 59496.75 | 31343.50 | 78331.47 |
| *COL8A1* | 29.40 | 52.82 | 124.47 | 515.43 | 860.51 | 1742.23 | 2380.11 | 417.43 | 2133.97 |
| *COL8A2* | 172.91 | 471.98 | 1400.09 | 1830.54 | 1304.47 | 1344.90 | 1087.56 | 240.36 | 745.65 |
| *COL9A1* | 487.37 | 2904.18 | 8900.67 | 12689.18 | 24243.56 | 46777.57 | 57855.93 | 26838.08 | 103560.60 |
| *COL9A2* | 695.59 | 3192.39 | 9143.09 | 15740.89 | 27228.59 | 56094.60 | 57042.83 | 33962.69 | 92938.82 |
| *COL9A3* | 170.38 | 1339.24 | 3826.45 | 6133.44 | 10639.87 | 19812.78 | 23702.90 | 11269.41 | 58400.92 |
| *COL10A1* | 6.43 | 15.96 | 13.96 | 39.86 | 125.65 | 450.54 | 1336.33 | 2551.24 | 10588.65 |
| *COL12A1* | 2047.31 | 2095.84 | 4630.80 | 9913.81 | 13137.12 | 19214.44 | 20272.06 | 7897.80 | 7001.24 |
| *COL13A1* | 103.89 | 175.37 | 233.99 | 289.25 | 404.83 | 533.26 | 259.59 | 44.32 | 39.14 |
| *COL14A1* | 304.58 | 121.36 | 650.23 | 6523.20 | 13200.84 | 15752.35 | 5095.64 | 896.92 | 344.34 |
| *COL15A1* | 55.71 | 421.52 | 592.38 | 741.19 | 617.45 | 586.35 | 844.13 | 447.29 | 2537.33 |
| *COL16A1* | 475.96 | 1104.45 | 1937.75 | 3863.33 | 6889.11 | 15555.74 | 12544.44 | 6428.85 | 8801.93 |
| *COL17A1* | 114.53 | 66.25 | 45.58 | 36.66 | 38.68 | 33.51 | 18.82 | 4.02 | 28.15 |
| *COL18A1* | 1130.23 | 1194.76 | 817.19 | 604.08 | 437.58 | 336.02 | 393.62 | 489.87 | 404.67 |
| *COL20A1* | 4.15 | 3.22 | 9.25 | 13.40 | 22.64 | 72.99 | 26.02 | 10.12 | 71.86 |
| *COL22A1* | 6.16 | 65.69 | 152.79 | 124.88 | 94.03 | 142.42 | 52.92 | 24.20 | 117.13 |
| *COL23A1* | 23.36 | 21.51 | 21.92 | 29.16 | 32.85 | 63.78 | 26.54 | 7.30 | 9.48 |
| *COL24A1* | 228.11 | 311.41 | 231.28 | 139.17 | 168.73 | 238.87 | 205.05 | 149.98 | 155.71 |
| *COL27A1* | 415.23 | 1813.25 | 3539.92 | 6578.20 | 10046.94 | 17431.51 | 11028.69 | 4995.78 | 3374.92 |
| *COL28A1* | 3.06 | 7.21 | 7.97 | 12.33 | 8.48 | 16.74 | 35.94 | 0.29 | 75.20 |

*Table S5*. Temporal expression profiles of transcription factors known to regulate *SOX9* transcription during chondrogenesis. ***EBF1*** was analysed further during our detailed TF analysis.

| Symbol | Day 0 | Day 1 | Day 2 | Day 3 | Day 4 | Day 6 | Day 10 | Day 15 | Mature |
|---|---|---|---|---|---|---|---|---|---|
| *ATF3* | 1915.32 | 689.34 | 450.15 | 270.81 | 595.16 | 818.89 | 8756.30 | 11440.28 | 17187.04 |
| *CEBPB* | 733.07 | 427.92 | 455.04 | 506.58 | 821.18 | 1873.39 | 7858.09 | 3933.55 | 13422.36 |
| *CEBPD* | 168.72 | 105.25 | 180.95 | 349.61 | 458.60 | 1071.17 | 5620.70 | 2625.20 | 9433.36 |
| *CHD2* | 3184.88 | 2382.41 | 2290.59 | 2748.74 | 3822.15 | 6104.76 | 8487.92 | 10317.49 | 2295.84 |
| *CREB1* | 655.62 | 792.99 | 599.40 | 523.63 | 391.41 | 277.53 | 304.94 | 244.18 | 494.64 |
| *CTCF* | 2324.43 | 2553.47 | 2246.06 | 2046.54 | 1744.63 | 1489.62 | 1288.90 | 1254.72 | 1620.65 |
| *E2F4* | 943.78 | 716.58 | 659.12 | 621.09 | 650.54 | 662.01 | 810.10 | 1578.41 | 607.47 |
| ***EBF1*** | **322.46** | **621.49** | **470.41** | **281.98** | **169.88** | **167.89** | **85.93** | **71.62** | **113.59** |
| *ELF1* | 652.99 | 412.79 | 339.29 | 414.00 | 496.48 | 802.09 | 1697.18 | 2560.93 | 1409.98 |
| *EP300* | 2419.27 | 2006.20 | 1447.91 | 1306.41 | 1068.59 | 1234.00 | 1169.41 | 1605.95 | 1115.46 |
| *EZH2* | 1379.85 | 1239.75 | 1063.57 | 1023.81 | 792.61 | 539.38 | 616.65 | 802.76 | 255.82 |
| *FOS* | 3103.13 | 444.49 | 298.41 | 325.41 | 614.10 | 1719.95 | 6518.75 | 2450.57 | 4693.71 |
| *FOSL2* | 190.52 | 38.94 | 30.50 | 50.77 | 64.63 | 119.00 | 466.75 | 397.03 | 1064.65 |
| *HDAC2* | 7948.10 | 6172.39 | 4814.27 | 4078.07 | 3810.38 | 3041.97 | 2957.87 | 2875.93 | 2096.28 |
| *HIF1A* | 3285.12 | 4713.90 | 4997.83 | 7390.27 | 11740.85 | 16975.14 | 29052.46 | 22257.59 | 71849.48 |
| *HNF4A* | 1.27 | 0.00 | 0.58 | 0.47 | 0.79 | 0.96 | 0.00 | 0.00 | 1.29 |
| *JUND* | 7638.38 | 3072.50 | 3807.85 | 3617.80 | 3050.72 | 3943.18 | 6110.79 | 5529.67 | 6373.74 |
| *MAFK* | 527.92 | 142.79 | 144.55 | 190.54 | 286.44 | 537.85 | 2307.69 | 3419.07 | 1214.07 |
| *MAX* | 533.61 | 398.20 | 469.02 | 445.11 | 469.49 | 454.41 | 430.54 | 514.75 | 433.73 |
| *MXI1* | 349.96 | 463.63 | 474.84 | 496.44 | 531.42 | 516.23 | 1861.61 | 1800.75 | 1777.41 |
| *MYBL2* | 1010.20 | 815.03 | 719.64 | 611.95 | 404.73 | 215.67 | 129.19 | 125.09 | 184.40 |
| *MYC* | 1210.76 | 471.46 | 362.67 | 397.74 | 523.28 | 670.73 | 1573.43 | 2612.71 | 963.77 |
| *NFIC* | 370.09 | 664.16 | 1047.23 | 1715.88 | 2060.71 | 3323.19 | 3827.22 | 5104.45 | 655.42 |
| *PBX3* | 613.01 | 810.35 | 947.34 | 1195.40 | 1222.81 | 1012.81 | 547.79 | 414.74 | 321.59 |
| *RAD21* | 5030.29 | 3580.88 | 3145.46 | 3092.87 | 3115.79 | 2773.81 | 2385.64 | 2401.29 | 2242.36 |
| *RBPJ* | 395.63 | 359.82 | 313.75 | 342.23 | 372.51 | 349.75 | 438.97 | 345.05 | 801.50 |
| *RELA* | 478.81 | 488.36 | 525.97 | 520.38 | 507.82 | 571.41 | 831.03 | 999.74 | 616.26 |
| *RXRA* | 142.36 | 195.68 | 215.33 | 248.82 | 210.35 | 267.04 | 566.96 | 707.19 | 291.66 |
| *SIN3A* | 1087.32 | 836.22 | 744.63 | 639.31 | 531.28 | 525.41 | 623.10 | 455.24 | 697.96 |
| *SP1* | 2288.09 | 1416.20 | 1494.05 | 1545.84 | 1551.36 | 1852.43 | 2491.66 | 3513.57 | 1813.89 |
| *SPI1* | 8.80 | 3.21 | 5.60 | 2.86 | 6.42 | 4.21 | 15.31 | 18.11 | 242.56 |
| *SRF* | 4385.40 | 1638.06 | 1322.58 | 1089.93 | 815.67 | 721.40 | 872.90 | 1403.32 | 949.99 |
| *TBL1XR1* | 979.88 | 791.77 | 749.69 | 786.74 | 805.71 | 947.30 | 1922.47 | 4545.72 | 797.52 |
| *TCF7L2* | 1607.11 | 1594.56 | 1404.38 | 1379.76 | 1455.94 | 1632.88 | 2693.22 | 3959.88 | 339.12 |
| *USF1* | 1575.52 | 1767.72 | 2092.99 | 1873.89 | 1532.55 | 1297.71 | 1161.04 | 1391.39 | 965.08 |
| *WRNIP1* | 152.34 | 336.72 | 423.10 | 466.91 | 453.70 | 408.59 | 316.82 | 315.81 | 366.63 |
| *ZBTB7A* | 530.14 | 402.33 | 412.18 | 489.63 | 507.67 | 751.80 | 991.99 | 960.84 | 474.59 |
